# Megakaryocyte emperipolesis arms neutrophils via intracellular provisioning

**DOI:** 10.64898/2026.08.04.742780

**Authors:** Julia K. Kuehn, Geon Ho Bae, Roxane Darbousset, Frank Y. Huang, Rebecca M. Hall, Josephin E. Schiffer, Dhruv Miglani, Spoorthi Balu, Alexandra Wactor, Dorsaf Ghaloussi, Olga Barreiro, Li Guo, Andrew S. Weyrich, Simon J. Cleary, Matthias Gunzer, Yoichiro Iwakura, David P. Hoytema van Konijnenburg, Joseph E. Italiano, Mark R. Looney, Éric Boilard, Wolfgang Bergmeier, Pierre Cunin, Peter A. Nigrovic

**Affiliations:** Department of Pediatrics, Division of Immunology, Boston Children’s Hospital, Harvard Medical School; Boston, MA, USA; Department of Biochemistry and Biophysics, and Blood Research Center, School of Medicine, University of North Carolina at Chapel Hill; Chapel Hill, NC, USA; Department of Immunology, Harvard Medical School; Boston, MA, USA; HMS Center for Immune Imaging, Harvard Medical School; Boston, MA, USA; Program in Molecular Medicine and Department of Internal Medicine, University of Utah; Salt Lake City, UT, USA; Department of Medicine, University of California; San Francisco, CA, USA; Institute for Experimental Immunology and Imaging, University of Duisburg-Essen; Essen, Germany; Leibniz-Institut für Analytische Wissenschaften ISAS -e.V.; Dortmund, Germany; Center for Experimental Medicine and Systems Biology, University of Tokyo; Tokyo, Japan; Louis Pasteur Center for Medical Research; Kyoto, Japan; Vascular Biology Program, Boston Children’s Hospital; Boston, MA, USA; Centre de Recherche du Centre Hospitalier Universitaire de Québec, Faculté de Médecine de l’Université Laval; Québec, QC, Canada; Department of Medicine, Division of Rheumatology, Inflammation, and Immunity, Brigham and Women’s Hospital, Harvard Medical School; Boston, MA, USA; Division of Stem Cells and Cancer, German Cancer Research Center (DKFZ) and DKFZ-ZMBH Alliance; Heidelberg, Germany; Heidelberg Institute for Stem Cell Technology and Experimental Medicine (HI-STEM gGmbH); Heidelberg, Germany; Faculty of Biosciences, Heidelberg University; Heidelberg, Germany; Bloodworks Northwest Research Institute; Seattle, WA, USA; Division of Hematology and Oncology, University of Washington; Seattle, WA, USA; Oklahoma Medical Research Foundation; Oklahoma City, OK, USA; Institute of Pharmaceutical Science & Centre for Lung Health, King’s College London; London, UK

## Abstract

Neutrophils are phenotypically heterogenous cells that mediate host defense and tissue homeostasis. Here, we identify emperipolesis – the evolutionarily conserved process by which neutrophils pass through megakaryocytes – as a phenotypically transformative route of egress from bone marrow. By intravital microscopy and 3-D histology, we show that the rapid form emperipolesis is markedly enhanced under inflammatory conditions. Neutrophils exit from megakaryocytes directly to the blood, acquiring exosomes enriched in proteins related to metabolism, migration, and immune function. This transfer induces a distinct neutrophil phenotype characterized by enhanced glycolysis, oxidative phosphorylation, cytokine release, and longevity. Correspondingly, emperipolesis-educated neutrophils display accelerated migration *in vitro* and *in vivo*. Disrupting emperipolesis does not alter circulating neutrophil abundance but impairs neutrophil infiltration into inflamed tissues, including *Pseudomonas aeruginosa*-infected lung. These findings establish emperipolesis as a mechanism by which megakaryocytes amplify neutrophil-mediated immunity.

## INTRODUCTION

Neutrophils are innate immune cells that are essential for pathogen defense and tissue repair ^1^. Once considered uniform, neutrophils are now recognized as functionally diverse and responsive to tissue-specific environmental cues ^2–7^. Interestingly, in bone marrow, some neutrophils invade megakaryocytes (MKs) in a distinctive cell-in-cell interaction termed emperipolesis (EP) ^8–12^. In EP, binding of neutrophil β2 integrins to ICAM-1 on MKs triggers actin-dependent entry into a vesicle termed the emperisome ^8,10,13^. Some neutrophils pass into the MK cytoplasm, where they accelerate pro-platelet formation ^8,10,14,15^. Neutrophils remain viable during EP and may exit rapidly or persist within MKs for an hour or longer, dichotomous behaviors termed fast and slow EP ^8,10^. These features distinguish EP from other cell-in-cell processes such as phagocytosis, efferocytosis, endocytosis, and nexocytosis, which culminate in the death of the engulfed cell ^9,16^.

This prolonged contact with MKs could have functional consequences for neutrophils, potentially contributing to its conservation over >90 million years of mammalian evolution ^8,9,17^. EP has been characterized primarily in hematologic disorders with markedly elevated EP rates, such as hereditary macrothrombocytopenia and myeloproliferative neoplasms, where frequencies can exceed 50% of MKs ^15,18–25^. In these settings, increased EP may reflect impaired efferocytotic clearance of senescent neutrophils and other condition-specific mechanisms ^15,24–26^ and can involve neutrophil degranulation with transfer of granule contents to MKs and platelets, contributing to platelet dysfunction, MK death, and marrow fibrosis ^15,21,22,26^. However, EP is also induced under stress conditions characterized by inflammation and increased neutrophil demand ^8,9,12,14,27–29^. In these contexts, EP is not associated with neutrophil degranulation or MK death, and its impact on neutrophils is unknown ^8,10,26,29^.

Here, we show that EP is common in healthy mice and increases strikingly in throughput across diverse inflammatory settings. Egressing neutrophils passed directly into circulation, identifying EP as a route for neutrophil exit from bone marrow. During EP, neutrophils acquired MK-derived exosomes and distinct transcriptional, metabolic, and inflammatory-response profiles, migrating more efficiently *in vitro* and *in vivo*. Using a mouse model of sustained circulating platelet abundance but marrow MK depletion, we found that elimination of EP altered the functional characteristics of circulating neutrophils and impaired neutrophilic inflammation in *Pseudomonas aeruginosa* pneumonia. Together, these findings establish EP as a phenotypically transformative novel pathway of neutrophil egress from marrow that enhances the potency of the circulating neutrophil pool.

## RESULTS

### Emperipolesis is enhanced across inflammatory pathologies

We and others previously observed increased EP in specific inflammatory conditions ^8,14,27^, leading us to suspect that enhanced EP is a general response to inflammation. To test this supposition, we quantified EP in mouse models of multiple inflammatory conditions using bone marrow histology. In the cecal ligation and puncture (CLP) model of bacterial polymicrobial sepsis ^30^, the frequency of MKs undergoing EP was approximately 2-fold higher in septic mice than in sham-operated controls (Fig. 1A). Similarly, in the K/BxN serum-transfer model of arthritis ^31,32^ and the bleomycin-induced model of systemic sclerosis (SSc) ^33,34^, inflamed mice exhibited elevated EP compared to vehicle-injected mice (Fig. 1B, Fig. S1A). Interleukin-1 receptor antagonist (*Il1rn*)-deficient mice that develop multi-organ inflammation due to unopposed IL-1*β* signaling ^35,36^ also displayed increased EP compared to wild type (WT) animals (Fig. S1D). Inflamed mice often had multiple (≥3) neutrophils per MK, a state rare among controls (Fig. S1B,C,E,F).

**Figure 1.**
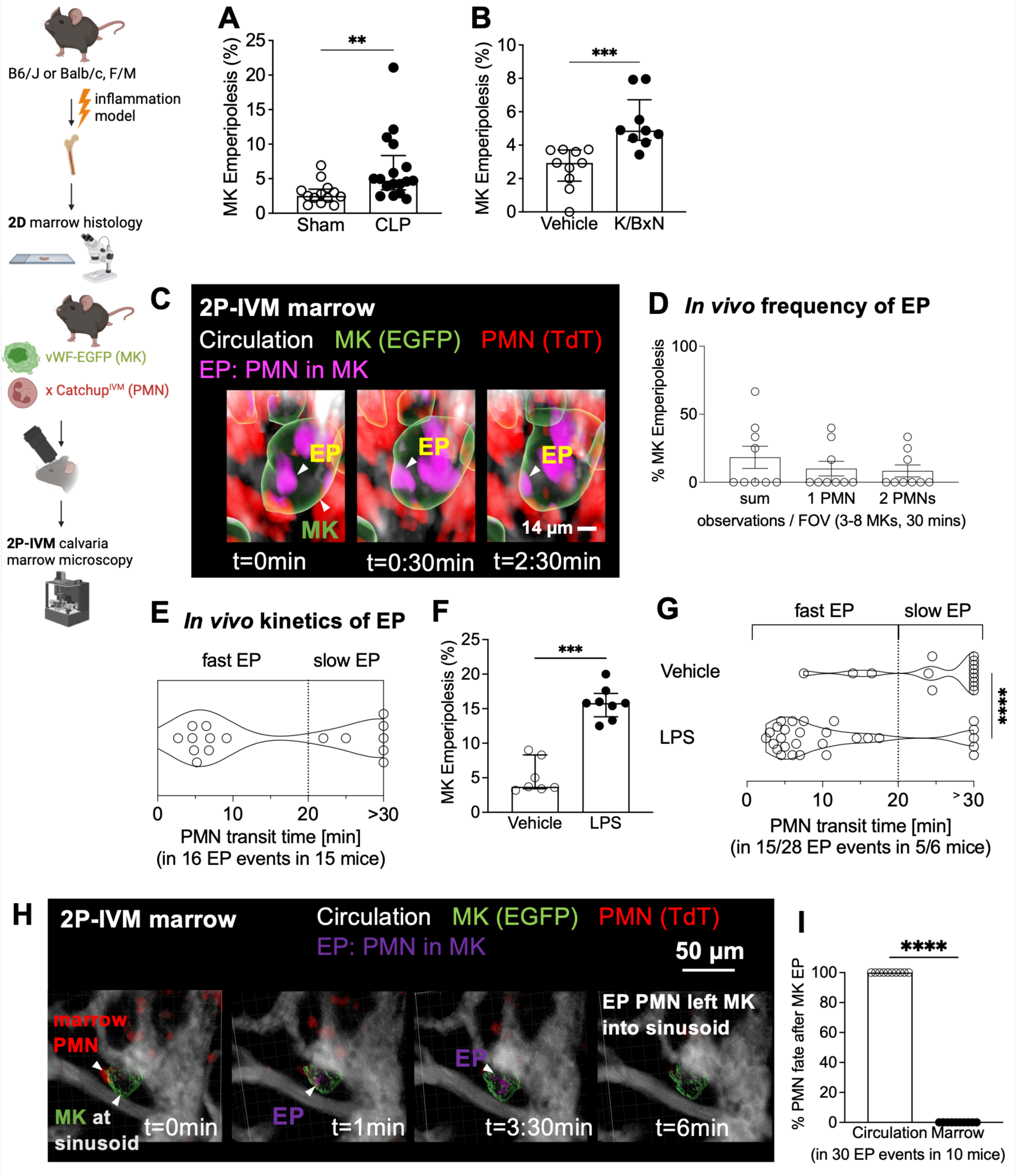
Enhanced emperipolesis is conserved across inflammatory pathologies and provides a trans-megakaryocyte route for neutrophil egress from the bone marrow. (A,B) Bone marrow megakaryocyte (MK) emperipolesis (EP) frequency in mouse inflammation models. (A) Cecal ligation and puncture (CLP) sepsis, day 4 vs sham (n=17/13). (B) K/BxN serum-transfer arthritis, days 7-8 vs vehicle (n=10/9). (C-I) Two-photon intravital microscopy (2P-IVM) of calvarial marrow in Catchup^IVM^ x vWF-EGFP mice; neutrophils tdTomato (red; magenta when internalized), MKs EGFP (green), vasculature Evans Blue (gray). (C-E) EP in healthy mice. (C) 3D reconstruction of neutrophil transit through an MK (arrowheads). (D) EP frequency per 30 min and FOV (3-8 MKs/FOV, n=9 mice): MKs internalizing any, 1, or 2 neutrophils. (E) EP transit times (16 EP events, n=7 mice). (F,G) Mice were treated with LPS (2 mg/kg i.p., n=8) vs vehicle (PBS; n=7). (F) EP frequency. (G) Transit times (28 LPS vs 15 vehicle EP events). (H,I) Neutrophils egress via MKs into circulation. (H) 3D reconstruction of exit (arrowheads). (I) Neutrophil fraction exiting MKs towards circulation vs marrow (30 EP events, 10 mice). (A-B,D,F,I) Bars, median values, error bars, IQR. Statistics: (A,B,F,G,I) Mann-Whitney U tests. Significance: **P ≤ 0.01, ***P ≤ 0.001, ****P ≤ 0.0001.

Because MKs reach 50-100μm in diameter, the true point prevalence of EP is likely to be underestimated by 5μm paraffin sections. We therefore generated whole-mount preparations that preserve the 3D architecture of the bone marrow. Indeed, the frequency of MKs engulfing neutrophils was higher than in 2D sections, in both control mice and arthritic or LPS-treated mice (Fig. S1G-I). Again, multiple neutrophils per MK were common in inflamed mice but rare in controls (Fig. S1G, blow-up).

To quantify the frequency of EP in living animals, we employed two-photon intravital microscopy (2P-IVM) in Catchup^IVM^ x vWF-EGFP neutrophil/MK reporter mice (Tomato+ neutrophils, EGFP+ MKs) ^37,38^. Over 30 minutes of time-lapse imaging, 10% of MKs engulfed neutrophils (typically single, occasionally multiple) (Fig. 1C,D, Movie S1), whereas the majority of MKs did not engage in EP, even over longer periods of observation (up to 2 hours). Neutrophils exhibited bimodal transit times, including fast events (<20 minutes, median passage time: 5.7 minutes) and slower or stationary events (≥20 minutes) (Fig. 1E), confirming *in vivo* the existence of fast and slow EP previously identified *in vitro* ^10,26^. Individual MKs supported both fast and slow EP, showing that these modes are particular to an individual neutrophil-MK interaction rather than to a specific MK state (Fig. S1J, Movie S2). After LPS, the frequency of EP tripled (Fig. 1F). Median transit time was shorter in inflamed mice (Fig. S1K), reflecting a selective increase in fast EP with LPS treatment (Fig. 1G). Together, these observations show that EP is common in healthy animals and induced strikingly in inflammation, both in frequency and throughput.

### Emperipolesis as a trans-megakaryocyte route for neutrophil egress from the bone marrow

To understand the function of EP, we evaluated where EP-engaged MKs localize within the marrow niche. By histology, we observed EP exclusively in large (>25 µm ⌀), morphologically mature MKs attached to sinusoids, the vessels that enable cellular trafficking from the marrow to the circulation (Fig. S2A). This finding echoes our previous *in vitro* finding that immature MKs rarely engage in EP ^10^. Interestingly, neutrophils undergoing EP could be observed at the interface between MKs and sinusoids, sometimes in direct contact with the circulation (Fig. S2B,C, Movie S3). These findings suggested that neutrophils might pass from the marrow into the blood via MKs, a previously undescribed route of egress. Indeed, such a course was readily observed, occurring in all neutrophils that were observed transiting through MKs (Fig. 1H,I, Movie S4).

### Neutrophils acquire megakaryocyte exosomes during emperipolesis

The observation that some neutrophils transit through MKs *en route* to the blood, rather than via a simpler direct route, suggested that EP might alter neutrophil phenotype. Taking advantage of an established *in vitro* model for EP ^8,10^, we noted that essentially all neutrophils passing through MKs acquired intracellular puncta that stained strongly for the abundant MK membrane protein CD41 (Fig. 2A). These puncta were approximately 200 nm in diameter, comparable to exosomes ^39^. Flow cytometry confirmed that CD41+ particles were transferred to neutrophils upon direct coculture with MKs but barely if separated by a transwell membrane, indicating that particles were not acquired from the supernatant (Fig. S3A,B). Consistent with small extracellular vesicle identity, the MK alpha granule protein PF4 was not transferred under either condition (Fig. S3C). Mitochondria did not transfer from MKs to neutrophils, as determined by co-culture of WT neutrophils with MKs from PF4-Cre x PhAM^fl/WT^ mice harboring green mitochondria specifically in the MK lineage.

**Figure 2.**
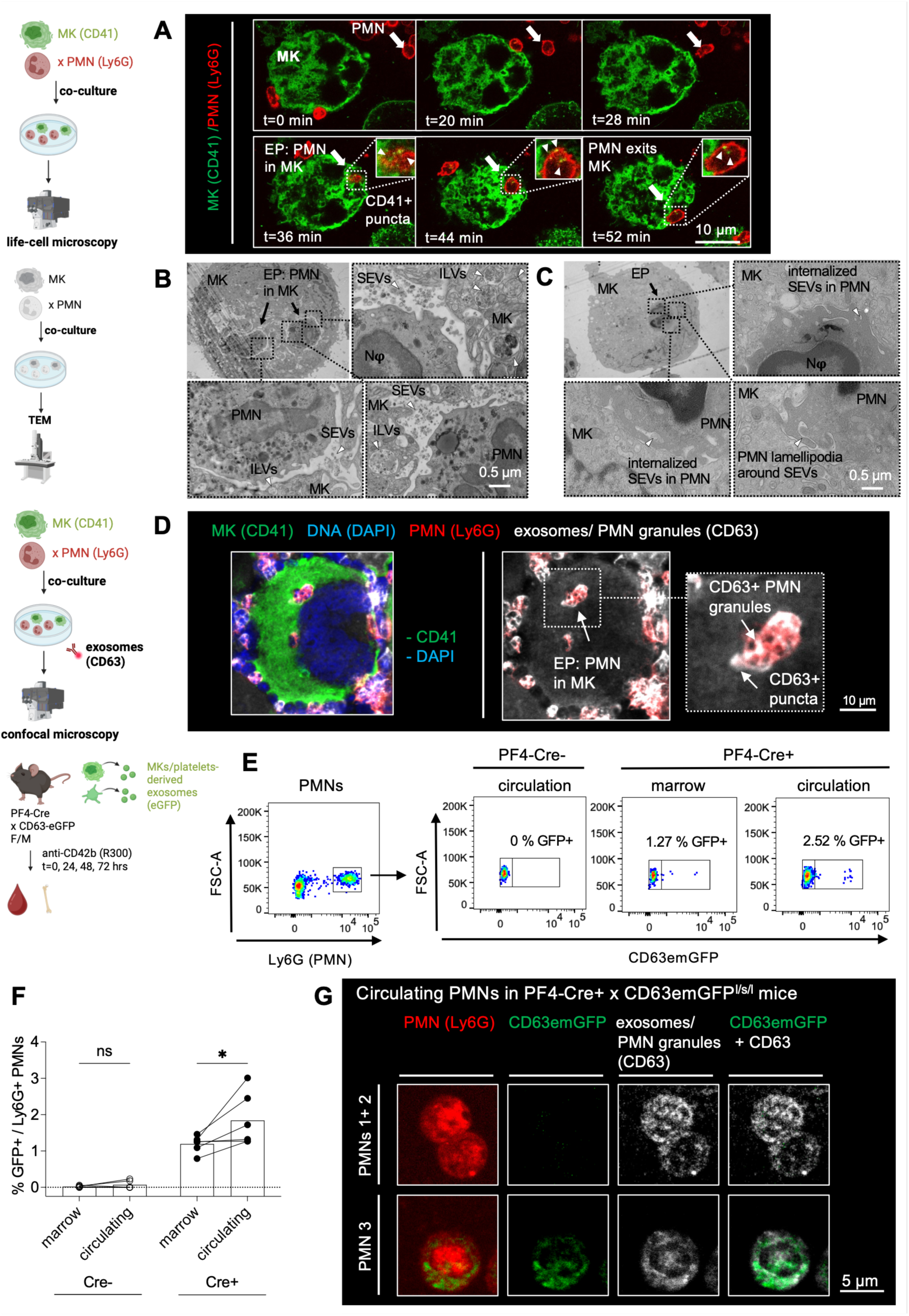
Transiting neutrophils acquire MK exomes during emperipolesis. (A) Time-lapse confocal imaging of MK–neutrophil co-cultures; MKs anti-CD41 (green), neutrophils anti-Ly6G (red). Arrow, EP neutrophil; arrowheads, CD41⁺ puncta internalized by the neutrophil. Representative of n=3. (B,C) Transmission electron microscopy of MK-neutrophil co-cultures. Representative of n=3. (B) MK multivesicular bodies with intraluminal vesicles (ILVs) adjacent to emperisomes; neutrophils surrounded by small extracellular vehicles (SEVs). (C) Neutrophil within an emperisome extends lamellipodia enclosing and internalizing SEVs. (D) Confocal microscopy of MK-neutrophil co-cultures. MKs anti-CD41 (green), neutrophils anti-Ly6G (red), DNA DAPI (blue), exosomes/granules anti-CD63 (gray). Left, composite; right and inset, EP event, CD63 and Ly6G only. Arrow, CD63⁺ neutrophil granules; arrowhead, CD63⁺ puncta surrounding EP neutrophils. Representative of n=3. (E-G) PF4-Cre+ x CD63emGFP^l/s/l^ mice and Cre- littermate controls (n=6/6) were treated with anti-CD42b (clone R300, 2 mg/kg daily x 72 h) to deplete platelets. (E) Flow-cytometry gating for GFP⁺ fraction within Ly6G⁺ neutrophils. (F) GFP⁺ fraction in matched marrow vs blood. (G) Confocal microscopy of circulating neutrophils from a Cre⁺ mouse: two GFP⁻ and one GFP⁺ neutrophil; anti-Ly6G (red) and anti-CD63 (gray); right, GFP-CD63 composite showing overlap. (F) Bars, median. Statistics: (F) Wilcoxon matched-pairs test. Significance: *P ≤ 0.05.

To explore the possibility that extracellular vesicles are transferred during EP, we performed transmission electron microscopy (TEM) on fixed MK-neutrophil co-cultures. MK multivesicular bodies containing intraluminal vesicles (ILVs) – the intracellular storage form of exosome precursors ^40^ – aggregated near internalized neutrophils (Fig. 2B). Some of these vesicles appeared to be released as exosomes into the emperisome, the intra-MK vacuole that houses the neutrophils just after MK entry ^8^ (Fig. 2B). Neutrophils within emperisomes exhibited lamellipodia-like structures, indicative of endocytosis, as well as internalized small extracellular vesicles (Fig. 2C), consistent with the acquisition of MK-derived exosomes during EP.

We further investigated exosome trafficking during EP in MK-neutrophil co-cultures by confocal microscopy, using CD63 as a marker of exosomes ^39–41^. We observed CD63+ punctate clusters surrounding internalized neutrophils, consistent with the TEM observations (Fig. 2D). CD63 and other exosome markers, such as CD81, CD9, and Alix, are also present in neutrophil granules, platelet dense granules, and other forms of small extracellular vesicles such as ectosomes ^41,42^, limiting specificity (see Fig. 2D). Yet in the presence of 20µM GW4869, a neutral sphingomyelinase inhibitor that halts exosome generation in cultured MKs (Fig. S3D) ^43,44^, these clusters were not observed, while CD63+ granules within neutrophils were still present, implying MK origin of the clusters rather than neutrophil degranulation.

To investigate exosome transfer *in vivo*, we generated PF4-Cre+ x CD63-emGFP^l/s/l^ mice expressing CD63-emGFP fusion protein in the MK-lineage ^45^. We confirmed that these mice serve as reporters for MK- and platelet-derived exosomes, with the majority of mature MKs and all circulating platelets expressing CD63-emGFP (Fig. S3E-G). To selectively study MK exosome transfer, we depleted circulating platelets with anti-CD42b, an antibody that does not reduce MK density (Fig. S3H,I) ^46–48^. Depletion was maintained for 72 hours, a period exceeding the lifespan of circulating neutrophils by approximately 7-fold ^49^. In Cre+ platelet-depleted mice, bone marrow neutrophils contained some GFP+ material (Fig. 2E,F). The proportion of neutrophils containing GFP+ material was higher in circulating than marrow neutrophils (Fig. 2E,G). Confocal microscopy revealed GFP+ material in some circulating neutrophils in Cre+ mice (Fig. 2G) but not in Cre- mice. The GFP+ material overlapped with the distribution of CD63+ material, did not morphologically resemble phagocytosed platelet debris (Fig. 2g, right bottom panel), and is unlikely to derive from phagocytosis because GFP fluorescence is quenched soon after entry into neutrophil phagolysosomes ^50^. These results confirm the presence of MK-derived exosomes in a subset of neutrophils, potentially acquired as the neutrophils enter the circulation via a trans-MK route.

### Megakaryocytes transfer proteins related to metabolism, migration and immunity to neutrophils

Exosomes are well-established regulators of cellular functions, and the transfer of proteins contained in exosomes are known mediators of these effects ^39,40^. To understand the implications of exosome transfer in EP, we employed proteomics and transcriptomics.

We used Stable Isotope Labelling by Amino Acids in Cell Culture (SILAC) ^51^ to selectively label MK proteins and identify the MK proteins transferred to unlabeled neutrophils. MKs were grown in media containing heavy isotope-labeled amino acids, achieving a labeling efficiency of approximately 70% (Fig. S4A). The labeled MKs were then co-cultured with unlabeled neutrophils to allow for emperipolesis, or, as a control, with neutrophils separated by 0.45µm transwell inserts to prevent direct contact while still allowing the exchange of soluble factors and exosomes. Following culture, sorted control neutrophils showed a labeling rate of 1.3%, whereas co-cultured neutrophils exhibited a rate of 8.2% (Fig. S4B). Of the 155 transferred proteins, two were unique to the control group, 31 were shared, and 122 were specific to the co-culture condition, indicating contact-dependent protein transfer from MKs to neutrophils (Fig. S4C, Table S1). PANTHER overrepresentation analysis ^52^ of the 122 peptides transferred only to co-cultured neutrophils revealed enrichment for Gene Ontology (GO) Cellular Component terms associated with the cell membrane, the actin cytoskeleton, and extracellular vesicles and exosomes (Fig. 3A, Table S2), consistent with exosome-mediated transfer. Further pathway analysis of the transferred peptides highlighted integrin signaling, cytoskeletal regulation, inflammatory chemokine and cytokine signaling, and metabolic pathways including the pentose phosphate pathway, glycolysis, and ATP synthesis (Fig. 3B, Table S3).

**Figure 3.**
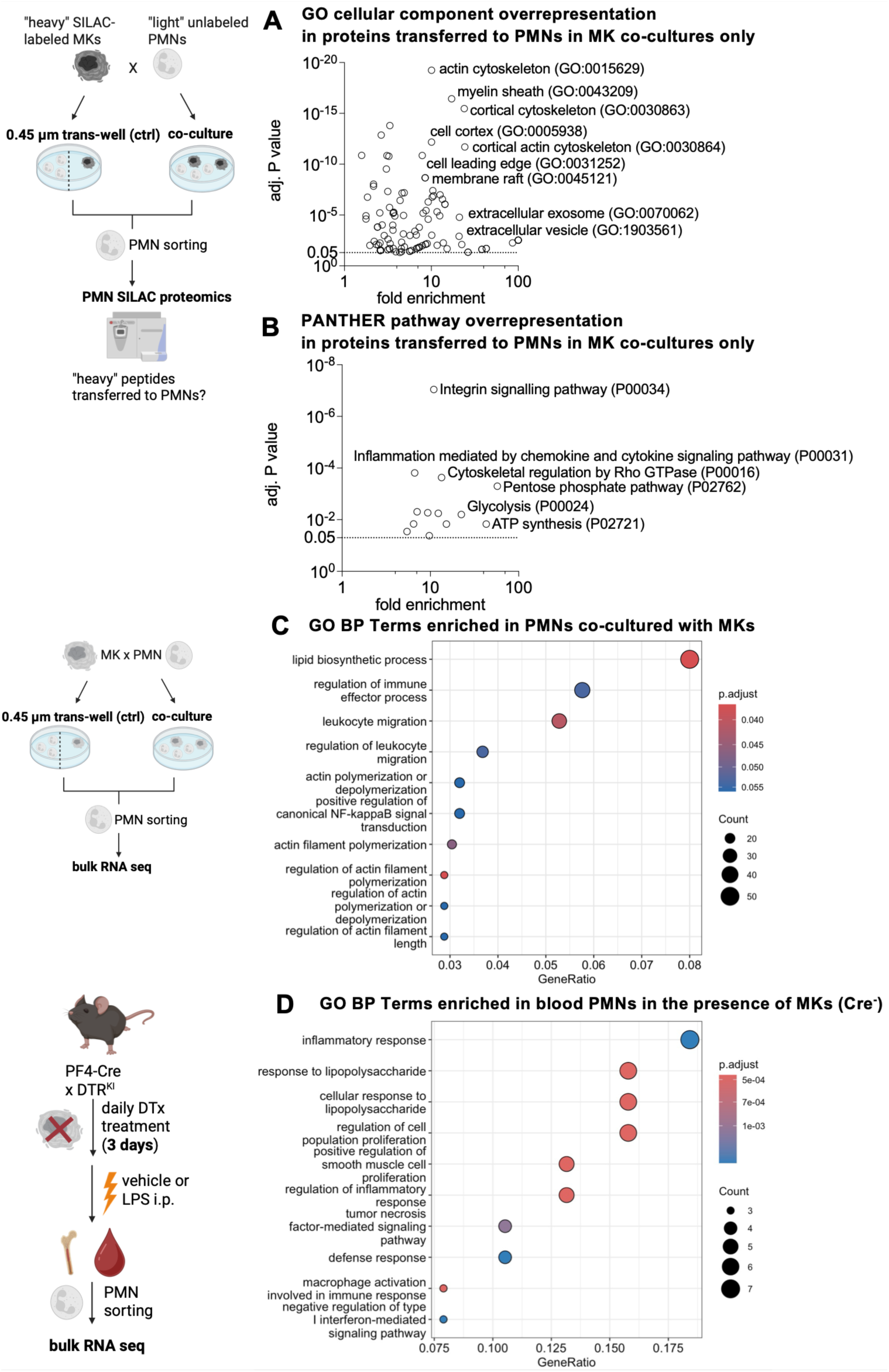
Megakaryocytes transfer proteins related to metabolism, migration, and immunity to neutrophils via exosomes and shape the neutrophil transcriptome. (A,B) SILAC proteomics to identify contact-dependent protein transfer from MKs to neutrophils. MKs were labeled with ^13^C^15^N-lysine/arginine; unlabeled neutrophils were co-cultured with MKs (EP) or separated by a 0.45 µm transwell (control) for 3 hours. Labeled proteins detected in FACS-sorted neutrophils were tested for overrepresentation of Gene Ontology (GO) Cellular Component (A) and PANTHER pathways (B), using the PANTHER Overrepresentation test tool. N=3 paired samples. (C) Bulk transcriptomic analysis of FACS-sorted neutrophils after co-cultures with MKs (EP) vs 0.45 µm transwell control (3 hours). Bubble chart of the top-10 enriched GO Biological Process terms in co-cultured neutrophils. N=8 paired samples. (D) Blood neutrophil transcriptome in the presence of MKs. PF4-Cre⁺ × iDTR^fl/fl^ vs Cre⁻ littermates received diphtheria toxin to deplete MKs (250 ng i.p. daily x 3 days), followed by vehicle or LPS treatment. FACS-sorted circulating neutrophils were analyzed for transcriptional changes associated with the presence and absence of MKs across vehicle/LPS treatment. Bubble chart of top-10 enriched GO Biological Process terms in MK-sufficient mice. N=3-5/group. Statistics: (A,B) Fisher’s exact test, FDR-adjusted (α<0.05), (C,D) hypergeometric test, FDR-adjusted (α<0.1).

To assess whether MKs alter neutrophils more globally, we compared the transcriptome and proteome of neutrophils co-cultured directly with MKs with those separated from MKs by 0.45µm transwell inserts. In line with the SILAC data, the transcriptome of co-cultured neutrophils exhibited enrichment in transcripts functionally associated with leukocyte migration and actin cytoskeletal regulation, including *Dock5* and *Prex1* (chemotaxis machinery), immune response, including *Pglyrp1* (peptidoglycan recognition protein), and metabolic regulation, including *Pfkfb4d* and *Gpx1*, which are relevant to glycolytic control and redox hemostasis, respectively (Fig. 3C, Fig. S4D, Tables S4-S7) ^53–57^.

Similarly, the proteome of co-cultured neutrophils exhibited enrichment in metabolic pathways, reflected in increased levels of key enzymes of the citrate cycle, including Idhb3, as well as an increase in MK/platelet-derived cell adhesion molecules, such as CD41, CD61 and thrombospondin 1 (Fig. S4E,F, Tables S8-10). Proteins associated with Fc gamma receptor-mediated phagocytosis and regulated cell death pathways were reduced (Table S10).

These findings reflect changes in the post-EP neutrophil proteome and transcriptome *in vitro*, suggesting that EP may change neutrophils with respect to metabolism, migration, immunological effector functions, and survival.

### Megakaryocytes shape the neutrophil transcriptome *in vivo*

We next developed a system to test the effect of MKs on neutrophils in living animals. EP can be blocked by paralysis of the actin cytoskeleton of either MKs or neutrophils ^8^, but this strategy is not viable *in vivo*. Therefore, we adapted the established PF4-Cre^+^ x iDTR^+/+^ MK depletion system ^58^. In Cre^+^ mice, treatment with diphtheria toxin (DTx) eliminates MKs within 1-2 days; platelets disappear more slowly as a function of their 5-day half-life but are not directly targeted by DTx since their survival is not dependent on protein synthesis. Since platelets strongly influence blood neutrophil biology, including the ability of neutrophils to infiltrate into tissues ^59,60^, this system yields a “window” wherein MKs are absent but platelets remain abundant. Specifically, after 3 days of DTx treatment, mice exhibit complete MK ablation, while platelet counts and the expression of CD62P, the major platelet ligand for interaction with neutrophils ^59^, remain close to normal (Fig. S5A-C). During this period, changes in neutrophil behavior are reasonably attributed to the impact of MKs, and to a lesser degree to other lineages expressing PF4, such as a small fraction of monocytes and macrophages and distal epithelial colon ^61^.

Employing this system, we assessed the impact of MK depletion on the neutrophil transcriptome across steady-state and inflammatory conditions. Blood and marrow neutrophils were sorted from MK-depleted animals and MK-sufficient mice treated with vehicle or LPS, and bulk transcriptomic changes associated with the presence of MKs across treatments were analyzed. Any effects caused by EP should become evident only in circulating neutrophils, since essentially all marrow neutrophils undergoing EP leave directly into the vasculature, while both marrow and blood neutrophils may still be influenced by other MK-dependent functions, such as maintenance of the marrow niche integrity and immune response ^62^.

In blood neutrophils from MK-sufficient mice, we observed increased expression of multiple transcripts, including *Pfn1*, which is required for actin cytoskeletal organization and efficient neutrophil migration ^63^ (Fig. S5D, Tables S11,S12). GO Biological Process (BP) term enrichment analysis revealed minimal enrichment in blood neutrophils from MK-depleted mice (Table S14) but numerous enriched terms in the MK-sufficient animals, including inflammatory and antibacterial response, cytoskeletal organization and chemotaxis, and cytokine production (Fig. 3D, Table S13). In contrast, the transcriptome of marrow neutrophils remained largely unchanged by MK depletion (Fig. S5E, Tables S15,16). Marrow neutrophils showed minimal GO BP term enrichment in the presence of MKs and no enrichment in their absence, and none of the transcriptional changes observed in marrow neutrophils overlapped with those in blood neutrophils (Table S17).

The *in vivo* transcriptional differences in blood neutrophils mirror the transcriptomic and proteomic shifts observed in neutrophils after MK co-culture *in vitro*, with both analyses suggesting that MKs shape neutrophil responses to inflammation, particularly programs linked to metabolism and migration.

### Megakaryocytes enhance neutrophil metabolism, including via exosomes

We next examined whether proteomic and transcriptomic changes translated to altered neutrophil function, initially focusing on metabolism, a key determinant of neutrophil maturation and functional diversification ^64–66^. Extracellular metabolic flux analysis revealed that neutrophils co-cultured with MKs exhibited higher glycolytic rates compared with neutrophils separated from MKs by 0.45µm transwell inserts (Fig. 4A, Fig. S6A). Basal respiratory capacity remained unchanged but maximal respiration was increased, albeit less than glycolysis (Fig. 4B, Fig. S6B). Following LPS stimulation, as expected ^67^, control neutrophils displayed increased glycolysis and reduced oxidative phosphorylation (Fig. S6C,D). In contrast, MK-co-cultured neutrophils maintained high oxidative phosphorylation alongside enhanced glycolysis (Fig. S6C,D). SILAC proteomics had revealed the transfer of glycolytic enzymes from MKs to neutrophils via exosomes, so we tested whether the metabolic shifts are exosome dependent. Pre-treating MKs with GW4869, which inhibits exosome production (Fig. S3D), prevented the enhancement of glycolysis, but not oxidative phosphorylation, indicating partial exosome dependency of these metabolic changes (Fig. 4C, Fig. S6E).

**Figure 4.**
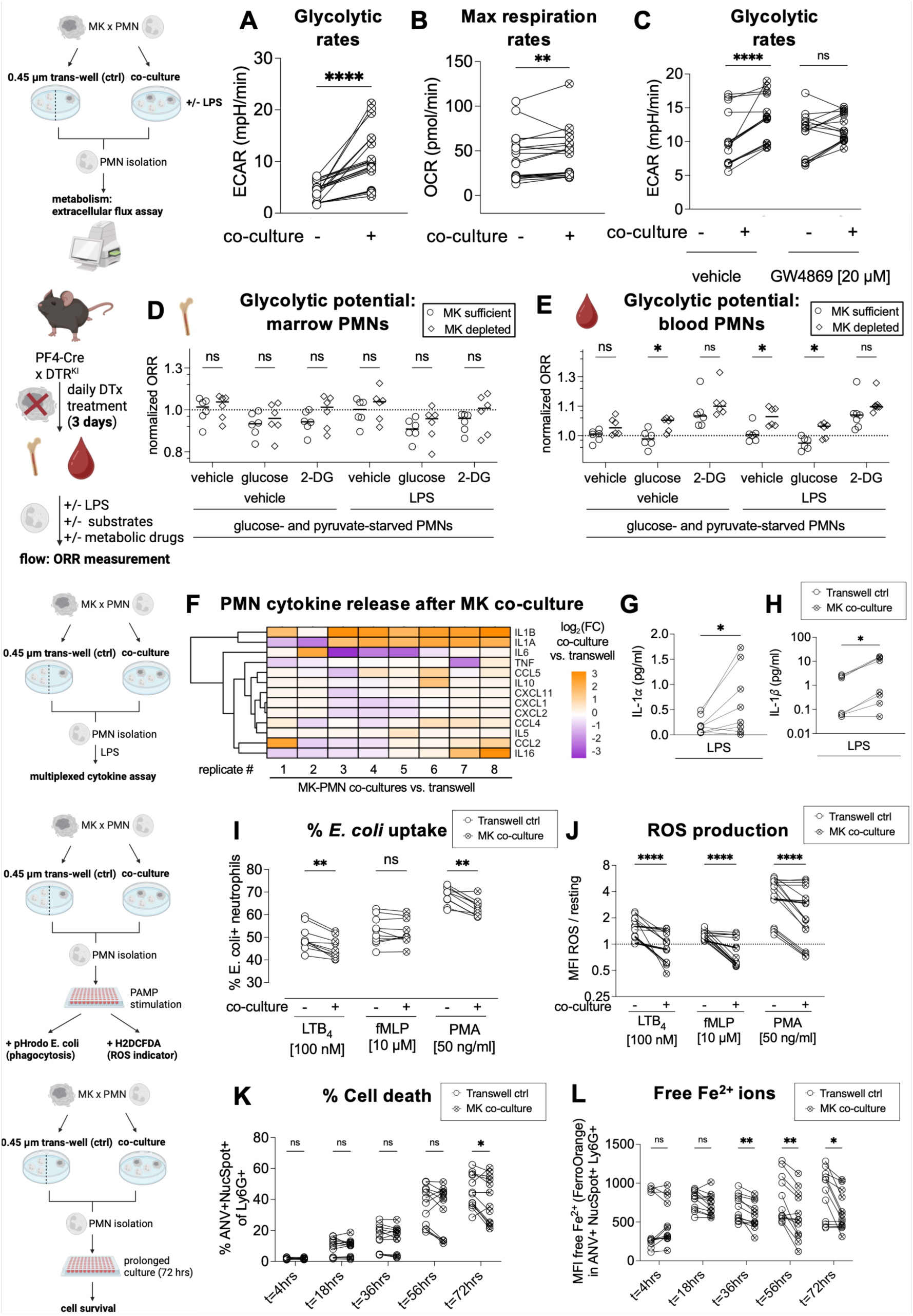
Megakaryocytes enhance neutrophil metabolism, cytokine production and survival but reduce phagocytosis and ROS production. (A-C) Metabolic profiling of neutrophils after MK co-culture (3 hours, EP) vs transwell control; ± exosome inhibitor GW4869 (20 µM). N=21(A,B)/15(C). (A) Glycolytic rate (ECAR, extracellular acidification rate) after glucose. (B) Maximal mitochondrial respiration (OCR, oxygen consumption rate) after FCCP. (D,E) Optical redox ratios (ORRs) in neutrophils from MK-depleted PF4-Cre⁺ × iDTR^fl/fl^ vs Cre⁻ littermates after DTx (250 ng i.p. daily × 3 days). N=6 paired marrow (D) and blood (E) per genotype. Cells were glucose/pyruvate-starved, ± LPS (10 µg/ml); ORRs by flow cytometry at baseline and after glucose and 2-DG, normalized to median vehicle ORR. (F-H) Neutrophil cytokine release after MK co-culture (3 hours, EP) vs transwell control, then +LPS (10 µg/ml). N=8 paired samples. (F) Heatmap of log₂ fold-change analyte concentration vs paired controls (Ward D2 clustering). (G) IL-1α, (H) IL-1β concentrations. (I) Neutrophil phagocytosis of fluorescent E. coli particles by flow cytometry after MK co-culture (3 hours, EP) vs transwell control, ± LTB4 (100 nM), fMLP (10 µM), or PMA (50 nM). N=10 paired samples. (J) Neutrophil ROS after MK co-culture (3 hours, EP) vs transwell control, ± LTB4 (100 nM), fMLP (10 µM), or PMA (50 nM); MFI CM-H2DCFDA by flow cytometry normalized to unstimulated. N=10 paired samples. (K,L) Neutrophil survival and labile Fe²⁺ after MK co-culture (3 hours, EP) vs transwell control, followed by 72 hours culture. (K) % Annexin V⁺ NucSpot⁺ Ly6G⁺ cells. (L) Fe²⁺ (MFI FerroOrange) in Annexin V⁺ NucSpot⁺ Ly6G⁺ neutrophils. (D,E) Lines, median. Statistics: (A-C,G-L) (multiple) Wilcoxon matched-pairs tests, FDR-adjusted (α<0.05); (D,E) multiple Mann-Whitney U tests, FDR-adjusted (α<0.05). Significance: *P ≤ 0.05; **P ≤ 0.01; ****P ≤ 0.0001. Abbreviations: CM-H2DCFDA: 5-(and-6)-chloromethyl-2’,7’-dichlorodihydrofluorescein diacetate, DCF: 2′,7′-dichlorofluorescein, 2-DG: 2-deoxy-glucose, FCCP: carbonyl cyanide-4 trifluoromethoxy phenylhydrazone.

To validate these findings *in vivo*, we examined neutrophil metabolism in MK-depleted mice, again expecting EP to affect only circulating neutrophils but not marrow neutrophils. Due to the limited yield of blood neutrophils, it was not possible to perform reliable extracellular flux analysis on an individual animal basis. Instead, we measured NADH and FAD levels by flow cytometry to calculate optical redox ratios (ORRs), which reflect the balance between oxidative phosphorylation (high ORRs) and glycolysis (low ORRs) ^68^. Marrow neutrophils from MK-depleted mice showed no differences in ORRs under glucose refeeding conditions or following treatment with the mitochondrial uncoupler FCCP, with or without LPS stimulation (Fig. 4D, Fig. S6F). In contrast, circulating neutrophils from MK-sufficient mice exhibited lower ORRs when re-fed with glucose, reversed by the glycolysis inhibitor 2-DG and augmented by LPS (Fig. 4E). Upon LPS stimulation, these cells also exhibited lower ORRs with FCCP, reversible with rotenone and antimycin A, inhibitors of the mitochondrial complexes I and III (Fig. S6G). Further, circulating but not marrow neutrophils in MK-depleted mice showed reduced glucose uptake capacity, an effect again augmented by LPS (Fig. S6H-J). Together, these data indicate that MKs are necessary for normal glycolytic capacity of circulating but not marrow-resident neutrophils *in vivo*.

### Megakaryocytes enhance neutrophil cytokine production and survival but reduce ROS production and phagocytosis

Proteomic and transcriptomic analyses suggested that, beyond metabolism, multiple neutrophil effector functions are altered in the presence of MKs. Consistent with this observation, using a multiplexed cytokine release assay, we found that following MK co-culture, LPS-stimulated neutrophils exhibited a distinct cytokine profile, characterized by increased IL-1α/β and reduced TNF secretion, with no change in the release of chemokines such as CXCL1, CXCL2, and CCL4 (Fig. 4F-H, Fig. S7A–D). Co-cultured neutrophils demonstrated reduced phagocytosis of *E. coli* particles by flow cytometric assays, although this defect was overcome by stimulation with fMLP (Fig. 4I). By microscopy, neutrophils containing internalized MK-derived CD41+ material phagocytosed *E. coli* particles less efficiently than CD41- neutrophils (Fig. S7E,F). Co-cultured neutrophils also produced less reactive oxygen species (ROS) at rest and after stimulation, as expected given the close association of phagocytosis and ROS generation (Fig. 4J) ^69^. Conversely, co-cultured neutrophils exhibited less cell death than control neutrophils (Fig. 4K), and showed reduced levels of labile Fe^2+^ ions, indicative of reduced oxidative and/or mitochondrial stress (Fig. 4L). Apoptosis as measured by the rate of caspase 3/7 activation was not reduced (Fig. S7G). Ferrostatin-1, a specific inhibitor of lipid peroxidation occurring in ferroptosis triggered by extreme Fe^2+^ elevation ^70^, also did not negate the reduced rates of cell death in co-cultured neutrophils (Fig. S7H).

### Emperipolesis enhances neutrophil migration

The observation that EP-educated neutrophils exhibit enhanced metabolism and live longer but have reduced phagocytosis and ROS production suggested that these cells are adapted to another energy-intensive function. A possibility suggested by the proteomic and transcriptomic data sets was migration. Indeed, in a trans-well migration assay, neutrophils co-cultured with MKs migrated more efficiently along an LTB_4_ chemotactic gradient compared to control neutrophils cultured alone or separated from MKs by a 0.45µm transwell insert (Fig. 5A). This effect was abolished by inhibition of MK exosome generation by GW4869 (Fig. 5B), but was not replicated by exposing neutrophils to isolated MK exosomes (Fig. S8A), consistent with dependence on exosome transfer during EP.

**Figure 5.**
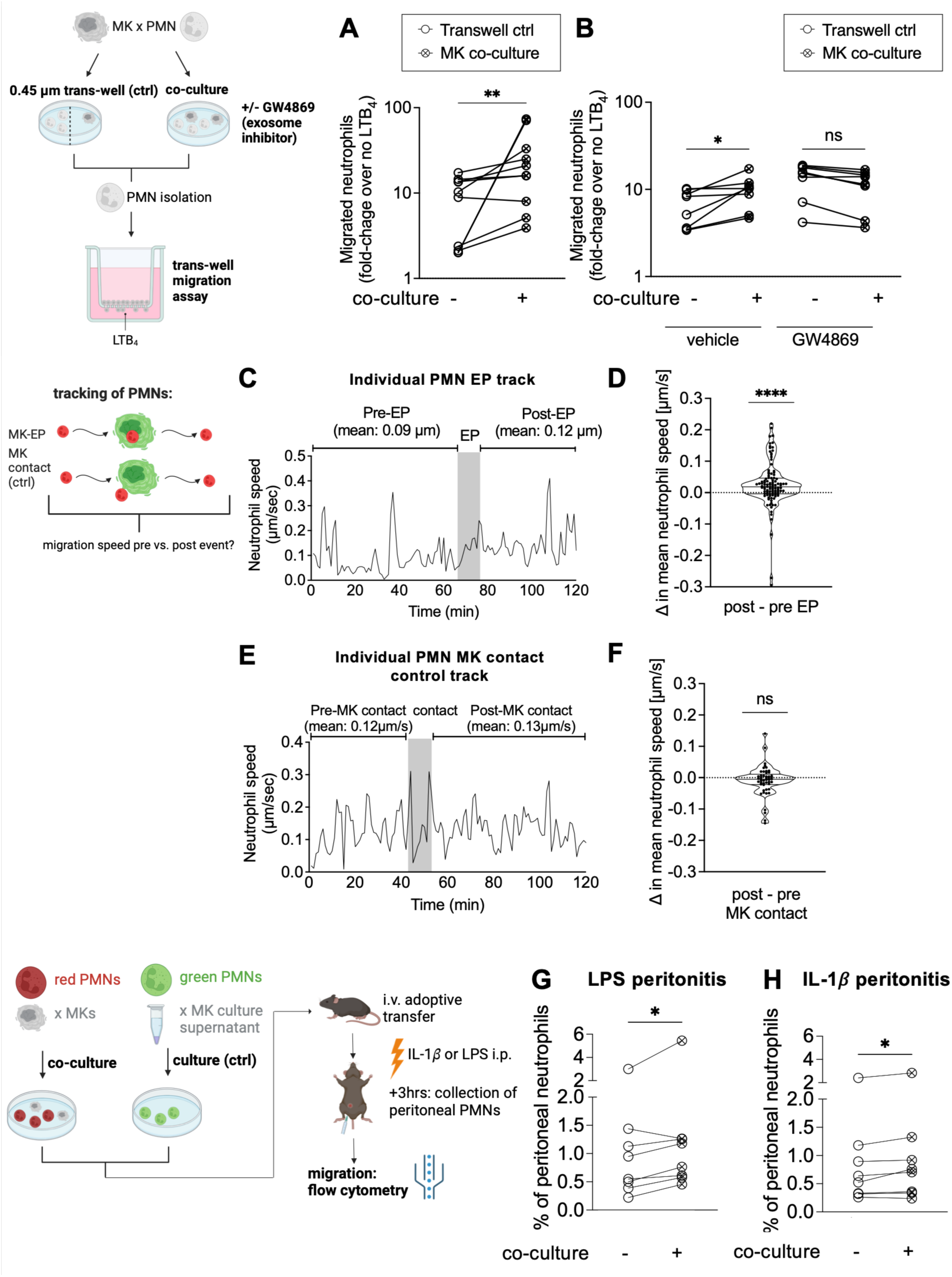
Emperipolesis enhances neutrophil migration. (A,B) Neutrophil transwell migration (3 µm pores, 2 hours) toward LTB_4_ (100 nM) after MK co-culture (3 hours, EP) vs transwell control. Migration quantified as fold-increase of Ly6G⁺ cells vs no gradient by flow cytometry. (B) ± exosome inhibitor GW4869 (20 µM) or vehicle (DMSO). N=10(A)/8(B). (C-F) Trans-MK neutrophil chemotaxis assay: MKs stained with anti-CD41-AF488 (green) were embedded in collagen and Catchup^IVM/Het^ neutrophils (TdT^+^, red) were imaged migrating through the MK-seeded matrix along an LTB_4_ (100 nM) gradient. Individual neutrophils speeds were recorded pre-/peri-/post-EP and MK-surface contact (controls). Representative EP (C) and control (E) tracks. Mean speed change post-EP (D) or post–surface contact (F). Pooled from n=3; 107 EP vs 50 control tracks. (G,H) Migration of neutrophils after MK co-culture (3 hours, EP) vs transwell control. Neutrophils were labeled and i.v. injected in mice subjected to peritonitis (G: LPS: 2 mg/kg i.p.; h: IL-1β: 1 µg/kg i.p., 1 hour). Frequency of dye⁺ Ly6G⁺ neutrophil fraction by flow cytometry. N=8 paired samples/model. (D,F) Violin plots: lines, median ± IQR. Statistics: (A,B,G,H) (multiple) Wilcoxon matched-pairs tests, FDR-adjusted (α<0.05); d,f one-sample Wilcoxon signed-rank test (μ₀=0). Significance: *P ≤ 0.05; **P ≤ 0.01; ****P ≤ 0.0001.

To track neutrophils passing through MKs during chemotaxis, we developed a three-chamber model. Marrow neutrophils in one chamber were induced to migrate toward LTB_4_ in another chamber, passing through a middle chamber containing MKs embedded in collagen matrix, where cells were imaged. As before, neutrophils passing through MKs acquired CD41+ puncta. We noted that mean neutrophil migration velocity of CD41+ neutrophils was enhanced compared to CD41- neutrophils (Fig. S8B). To test whether this phenomenon reflected a pre-existing characteristic of neutrophils destined to engage in EP, or instead the impact of EP itself, we tracked the migration velocity of individual neutrophils before and after EP, comparing neutrophils that passed through MKs with neutrophils that simply encountered the MK surface (Fig. S8C,D). Indeed, EP selectively enhanced migration speed while cell contact had no effect (Fig. 5C-F). Neutrophil velocities before EP were not different from those before MK surface contact alone (Fig. S8E). Mean migration velocity before EP correlated linearly with mean velocity after EP, indicating that EP increased neutrophil migration speed additionally and independent of initial velocity (Fig. S8F). Most EP events observed were fast EP (≤20 min). We did not observe a correlation between EP duration and neutrophil velocity gain after EP (Fig. S8G). Neutrophils undergoing EP within the same MK exhibited variable alterations in velocity, consistent with our *in vivo* observation above that the fate of neutrophils engaged in EP with the same MK may differ (Fig. S8H).

We tested the migratory capacity of EP-educated neutrophils *in vivo* using adoptive transfer. Neutrophils co-cultured with MKs and control neutrophils cultured in MK-conditioned media were adoptively transferred at a 1:1 ratio into mice subjected to LPS- or IL-1*β*-induced peritonitis. Co-cultured neutrophils showed enhanced migration into the peritoneum, implicating EP in enhanced neutrophil migration *in vivo*, although the overall effect size was modest (Fig. 5G,H).

### Emperipolesis enhances neutrophil entry into inflamed tissues

The assembled observations indicate that EP confers distinct metabolic, functional, and migratory characteristics upon some neutrophils, suggesting that MK-educated neutrophils might enhance neutrophil-mediated immune defense. We sought to test this possibility *in vivo*.

Using our PF4-Cre^+^ x iDTR^fl/fl^ system to selectively ablate MKs *in vivo*, we first tested our assumption that EP is *unlikely* to make an important contribution to circulating neutrophil abundance, given the ready alternate route for neutrophils to leave the marrow via bone marrow sinusoids not occupied by MKs. Despite MK ablation, PF4-Cre^+^ mice showed no change in circulating neutrophil count compared to those in MK-sufficient PF4-Cre^-^ littermate controls, either at baseline, following stress mobilization by G-CSF or CXCL1, or in recovery from peripheral neutrophil ablation via a depleting antibody (G-CSF increased EP frequency in marrow, while CXCL1 and peripheral neutrophil depletion did not; G-CSF: Fig. S9A,B, CXCL1: Fig. S9C,D, peripheral neutrophil depletion: Fig. S9E,F). Thus, while EP is common and increases with systemic inflammation, it does not measurably contribute to neutrophil abundance in blood in steady state or during stress mobilization.

Transcriptomic data suggested that MKs may regulate neutrophil migration in response to bacterial infection *in vivo* (see Fig. 3). To test this possibility, we induced a sublethal *Pseudomonas aeruginosa* (PsA) pulmonary infection in MK-depleted but platelet-sufficient mice ^71^. PsA infection for 4 hours triggered neutrophil pulmonary recruitment without systemic spread, as confirmed by the absence of bacteremia and undetectable spleen and liver bacterial loads. Four hours post inoculation, EP frequency in the marrow increased, alongside significant lung neutrophil influx (Fig. 6A,B). In MK-depleted mice, marrow and circulating CD11b^+^Ly6G^+^ neutrophils were unchanged, but lung neutrophil recruitment was reduced, while the number of other myeloid cells, including the macrophage-enriched CD11b^+^Ly6C^-^F4/80^+^ population, remained stable (Fig. 6B, Fig. S10A-C). In MK-sufficient mice, marrow EP frequency correlated with lung neutrophil abundance (Fig. 6C). Cytokine and chemokine levels were largely unchanged in plasma (Fig. S10D-G), but many analytes were reduced in the lungs of MK-depleted mice, notably IL-1α, IL-1β, IL-6, and TNF (IL-1β: Fig. 6D, others: Fig. S10H-K); of note, IL-1β levels in MK-sufficient mice approximated those of neutropenic mice infected with PsA (Fig. 6D). Neutrophil-recruiting chemokines, such as the CXCL1/2-inducing IL-17A and CXCL1 and CXCL2 themselves, remained unchanged in plasma and lung (plasma: Fig. S10D-G, lungs: Fig. S10H-K). These shifts in cytokine profile mirror our *in vitro* observations of reduced IL-1α/β secretion and stable chemokine release in neutrophils cultured without versus with MKs (see Fig. 4F-H). Histology confirmed reduced cellular infiltration (Fig. 6E) and revealed reduced overall inflammatory burden in MK-depleted mice, similar to levels observed in PsA-infected neutrophil-depleted mice (Fig. 6F). Reduced lung inflammation correlated closely with reduced neutrophil influx (Fig. S10L). While neutrophils are essential for protecting mice from septic shock by controlling PsA proliferation ^72,73^, neutrophils themselves can also contribute to lung injury in PsA infection, particularly at low bacterial burdens, a scenario consistent with our observations ^73,74^. Together, these findings suggest that MKs are required for the pulmonary recruitment of fully functional neutrophils in early sublethal PsA infection *in vivo*.

**Figure 6.**
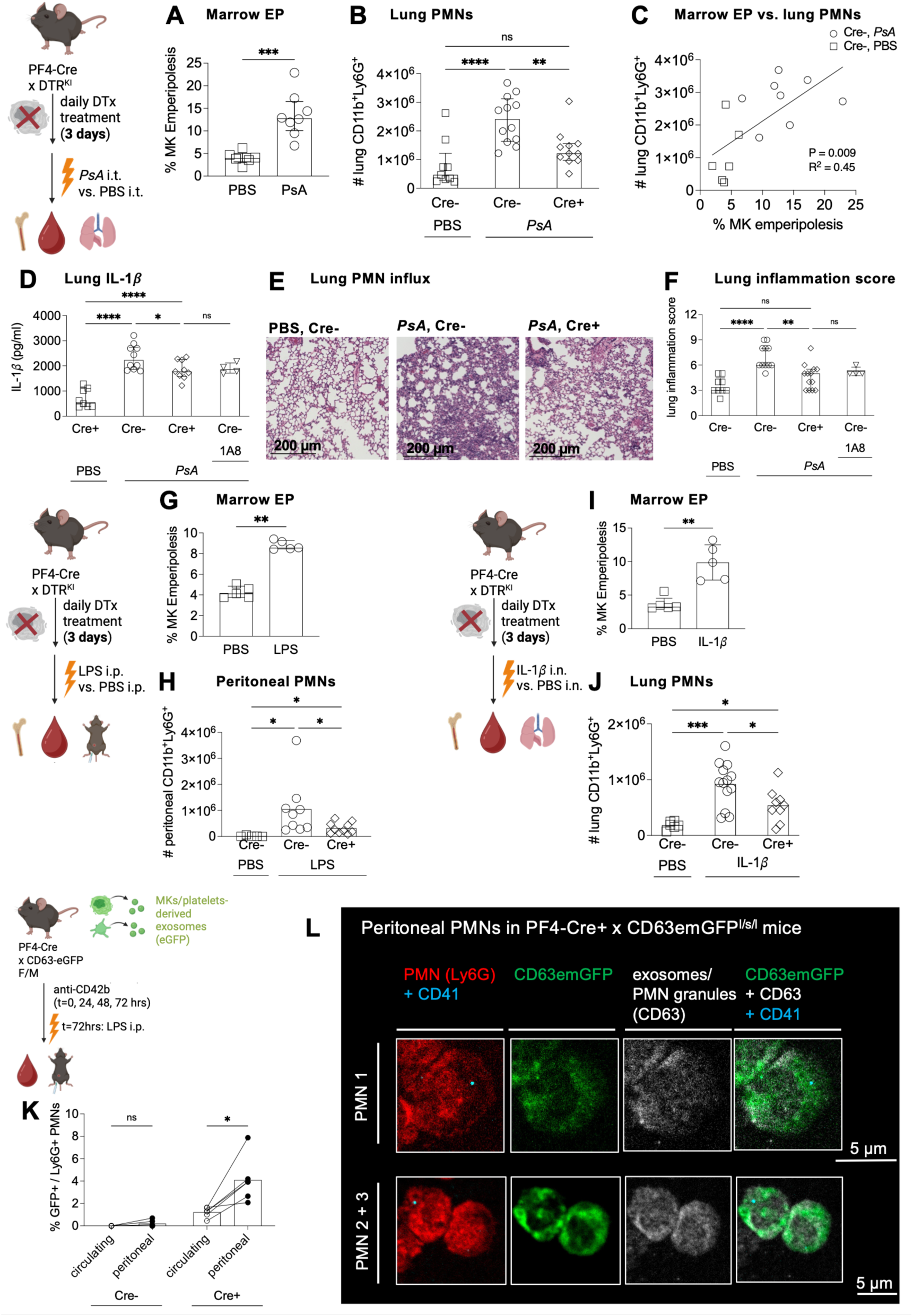
Emperipolesis enhances neutrophil entry into inflamed tissues. (A-F) Pseudomonas aeruginosa (PsA) pneumonia in MK-depleted PF4-Cre⁺ × iDTR^fl/fl^ vs Cre⁻ littermates (250 ng DTX i.p. daily × 3 days). Pneumonia was induced by i.t. inoculation with PsA (strain PA103, 3×10^5^ CFU, 4 hours) vs vehicle (PBS, control); neutrophils were depleted with 1A8 where indicated (24 h prior). (A) Bone-marrow MK EP frequency in PsA vs controls (n=6/8). (B) CD45⁺Ly6G⁺CD11b⁺ lung neutrophils (n=9/12/11). (C) Correlation between lung neutrophils and EP frequency (MK-sufficient only; n=21). (D) Lung IL-1β by ELISA of tissue homogenates (n=8/10/9/4). (E) Representative H&E lung histology showing cellular infiltration. (F) Histologic lung inflammation score (≥15 FOVs/sample; n=9/12/11/4). (G,H) LPS peritonitis (2 mg/kg i.p.; 2 hours) in MK-depleted PF4-Cre⁺ × iDTR^fl/fl^ vs Cre⁻ littermates (250 ng DTx i.p. daily × 3 days). (G) Bone-marrow MK EP in LPS vs controls (n=6/8). (H) CD45⁺Ly6G⁺CD11b⁺ neutrophils in peritoneal lavage (n=5/9/9). (I,J) IL-1β pneumonitis (1 µg/kg i.n.; harvest 4 hours) in MK-depleted PF4-Cre⁺ × iDTR^fl/fl^ vs Cre⁻ littermates (250 ng DTx i.p. daily × 3 days). (I) Bone-marrow MK EP in IL-1β vs controls (n=5/5). (J) CD45⁺Ly6G⁺CD11b⁺ lung neutrophils (n=6/13/9). (K,L) LPS peritonitis (2 mg/kg i.p.; harvest 2 hours) in platelet-depleted PF4-Cre⁺ × CD63emGFP^l/s/l^ MK exosome reporter mice (anti-CD42b/R300, 2 mg/kg daily × 72 h) vs Cre⁻ controls. (K) GFP⁺ fraction of matched circulating vs peritoneal neutrophils. (L) Confocal microscopy of GFP⁺peritoneal neutrophils from a Cre⁺ mouse, Ly6G (red), CD41 (cyan), CD63 (gray); left, Ly6G-CD41 overlay; right: GFP-CD63-CD41 overlay (representative of n=2). (A,B,D,F-K) Bars, median; error bars, IQR. c Line, linear regression. Statistics: (A,G,I) Mann-Whitney U; (B,D,F,H,J) one-way ANOVA, Holm-Šídák-adjusted (α<0.05); (C) linear correlation with t test; k multiple Wilcoxon matched-pairs tests, FDR-adjusted (α<0.05). Significance: *P ≤ 0.05; **P ≤ 0.01; ***P ≤ 0.001; ****P ≤ 0.0001.

To validate our findings, we examined additional models of neutrophil recruitment to sites of inflammation. Peritonitis induced by i.p. LPS triggered increased EP frequency in the marrow alongside neutrophil influx into the peritoneal cavity (Fig. 6G,H). MK-depleted mice exhibited normal bone marrow and blood neutrophil counts yet demonstrated reduced recruitment to the peritoneal cavity (Fig. 6H, Fig. S10M,N). Similarly, pneumonitis triggered by i.n. IL-1β induced increased EP in marrow and neutrophil infiltration into the lungs, with MK-depleted mice displaying diminished neutrophil recruitment to the lungs, while marrow and blood neutrophil counts remained unaffected (Fig. 6I,J, Fig. S10O,P).

Our *in vitro* findings suggested that EP-educated neutrophils gain enhanced migratory capacity through internalization of MK-derived exosomal material. To examine if this mechanism also operates *in vivo*, we investigated whether neutrophils migrating to inflammatory sites carry MK-derived material. In IL-1β-induced peritonitis, neutrophils recovered from the peritoneal cavity contained CD41^+^ puncta resembling those transferred in MK-neutrophil co-cultures (Fig. S11A, Movie S5, and see Fig. 2A). To confirm MK lineage origin of this material, we induced peritonitis in PF4-Cre^+^ x mT/mG MK/platelet membrane reporter mice. Up to 1% of peritoneal neutrophils displayed green-fluorescent puncta, showing a pattern consistent with MK-derived vesicles (Fig. S11B,C). These GFP^+^ neutrophils lacked MK/platelet α-granule markers PF4 and vWF, suggesting selective uptake of MK exosomal material rather than MK/platelet-derived α-granular material or platelet debris (Fig. S11D).

Finally, we tested MK exosomal transfer *in vivo* in PF4-Cre+ x CD63-emGFP^l/s/l^ MK exosome reporter mice. We depleted platelets in these mice, resulting in the majority of mGFP+ material being MK-derived exosomes. When subjected to LPS-induced peritonitis, the proportion of GFP⁺ neutrophils was higher in peritoneal neutrophils than in blood neutrophils, implying that neutrophils containing MK-derived exosomal material have higher migratory potential (Fig. 6K). Confocal microscopy confirmed that the GFP^+^ material was intracellular and colocalized with CD63^+^ and CD41^+^ intracellular material, while no CD41^+^ material was detected on the neutrophil surface (Fig. 6L). Together, our results confirm that EP confers a neutrophil phenotype that enhances both individual and collective neutrophil recruitment to sites of inflammation.

## DISCUSSION

Emperipolesis (EP), the transit of neutrophils through bone marrow megakaryocytes (MKs), is evolutionarily conserved, occurs in healthy animals, and is differentially regulated in disease ^8–12^. Here, we show that EP is broadly upregulated across diverse inflammatory contexts, that it provides a trans-MK route for neutrophil egress from the marrow, and that it enhances neutrophil metabolic fitness and migratory capacity via MK-derived exosome transfer. Abrogating EP *in vivo*, we confirm that EP is required for efficient neutrophil recruitment and cytokine production. Thus, we identify EP as an interaction between the hematopoietic and innate immune compartments that licenses neutrophilic inflammation.

By histology and intravital microscopy, we show that EP is confined to large, morphologically mature MKs attached to marrow sinusoids at the interface between marrow and circulation. Only a fraction of sinusoid-attached MKs engaged in EP, suggesting that a subset within this population is specialized for EP, consistent with the notion that MKs are transcriptionally and functionally heterogeneous ^62^. A distinct subset of small, low-ploidy, non-attached MKs with an “immune MK” transcriptional signature has been described ^62,75,76^, but we and others have not observed this subset to undergo EP ^26^. Instead, mature high-ploidy MKs, which also express a broad repertoire of immune receptors, appear strategically well positioned to sense systemic inflammatory cues and rapidly increase EP in response ^77^. MK ablation did not alter circulating neutrophil numbers, even during stress granulopoiesis, indicating that EP contributes little to bulk neutrophil output. Rather, the transcriptional and functional changes observed support a model in which EP reprograms a subset of neutrophils at the marrow-blood interface, allowing these neutrophils to fulfill a distinct role within the circulating neutrophil pool.

During EP, MKs rewire the neutrophil proteome, transcriptome and effector functions, both *in vitro* and *in vivo*. A central role in this process is played by MK-derived exosomes released into the emperisome. CD41^+^ material in neutrophils has previously been attributed to uptake of platelet-derived vesicles or platelet debris ^78–80^, but our data reveal direct transfer from MKs as an additional mechanism, with enrichment of such neutrophils in inflamed sites reflecting associated enhancement of migratory capacity. Using SILAC-based proteomics, we found that MKs transferred membrane, cytoskeletal, and vesicle-associated components, with pathways linked to integrin signaling, migration, inflammation, and metabolic regulation, aligning with the phenotypic changes observed in neutrophils after EP and consistent with the known ability of exosomes to reprogram recipient cells ^39,40^. Correspondingly, inhibiting MK exosome generation largely blunted MK-induced migratory and metabolic changes of neutrophils.

Functionally, MKs increased the metabolic capacity of neutrophils undergoing EP, particularly under inflammatory conditions, a setting in which neutrophils typically favor aerobic glycolysis at the expense of oxidative phosphorylation ^67^. EP allowed neutrophils to maintain high oxidative phosphorylation while further increasing glycolysis. Metabolism holds a central role in neutrophil biology as many effector functions are energy intensive ^64,66,81^. EP-mediated metabolic rewiring may thus license the other functional changes observed, including increased tissue infiltration and cytokine release in inflammatory environments *in vivo*. These environments are typically limited in nutrients and oxygen, so neutrophils critically depend on increased glycolysis and mitochondrial integrity for survival and execution of effector functions ^64,66,67,81,82^. MK-exposed neutrophils also produced less ROS, showed reduced phagocytosis, and survived longer with evidence of less oxidative stress, which may reflect reduced shunting of glycolytic intermediates into the pentose phosphate pathway, curbing oxidative burst and thus contributing to redox hemostasis and prolonged survival ^83^.

Intriguingly, although EP-educated neutrophils likely reflect a small proportion of circulating neutrophils (∼1-2% based on exosome content), ablation of EP decreases neutrophil infiltration into inflamed tissues by ∼50%. Further, while EP-educated neutrophils are enriched in inflammatory exudates they remain a small minority. These observations suggest that the role of EP-derived neutrophils may be to serve as “pathfinders”, guiding other neutrophils via soluble mediators and other mechanisms to coordinate the global neutrophil response. Such a role is further suggested by the observations that post-EP neutrophils acquire mobility and longevity at the expense of the effector functions phagocytosis and ROS production. If this speculation is correct, then just as tissue-level signals adapt neutrophils to serve specific local roles, EP may adapt a subset of blood neutrophils to play orchestrate neutrophil recruitment, offering a plausible explanation for the evolutionary conservation of EP as well as its marked induction under inflammatory conditions.

Our study has limitations. EP *in vitro* depends on actin remodeling and integrin engagement ^8^, but disrupting these pathways *in vivo* would broadly affect MK and neutrophil biology. To approximate neutrophil behavior without EP, we refined a mouse model in which MKs can be ablated by diphtheria toxin while largely preserving platelet counts, avoiding confounding thrombocytopenia. However, our model removes all MK functions, which are broad and include cytokine production and marrow-niche regulation ^62,77,84–90^, and still offers only a short window before platelet counts decline, prohibiting analysis of long-term outcomes. Moreover, because we cannot specifically identify EP-experienced neutrophils, our analyses necessarily involve mixed populations, diluting EP-specific signatures and preventing definitive attribution of findings to EP as opposed to other forms of interaction between neutrophils and MKs. We have not studied the functional advantages conferred by EP *in vivo* beyond migration and cytokine release, nor its role in sterile autoimmunity, tumor immunity, or chronic low-grade inflammation associated with aging and atherosclerosis. Finally, the relevance of EP-induced neutrophil reprogramming in human disease remains uncertain, although notably human EP closely resembles murine EP morphologically and also increases with inflammation^8,28^.

Despite these limitations, our findings identify EP as an inflammation-responsive interaction between MKs and neutrophils that equips a subset of neutrophils with enhanced metabolic fitness and migratory capacity, contributing to neutrophil diversity and allowing a small neutrophil subset to support rapid but controlled recruitment of neutrophils during inflammation.

## METHODS

### Mouse strains

The following validated animal strains were used: C57BL/6J (WT, strain #000664), BALB/cJ (strain #000651), mT/mG (B6.129(Cg)-Gt(ROSA)26Sortm4(ACTB-tdTomato,-EGFP)Luo/J, #007676), PF4-Cre (C57BL/6-Tg(Pf4-icre)Q3Rsko/J, #008535), Ai14 (B6.Cg-Gt(ROSA)26Sortm14(CAG-tdTomato)Hze/J, #007914), iDTR (C57BL/6-Gt(ROSA)26Sortm1(HBEGF)Awai/J, #007900), PhAM^fl/WT^ (B6;129S-Gt(ROSA)26Sortm1 (CAG-COX8A/Dendra2)Dcc/J, # 018385) and CD63-emGFP^l/s/l^ (C57BL/6J-Gt(ROSA) 26Sorem1(CAG-Cd63/EmGFP)Adly/J, #036865) mice were purchased from The Jackson Laboratory (USA). Catchup mice were a gift from Matthias Gunzer (University Hospital Duisburg-Essen, Essen, Germany) ^37^. vWF-GFP mice were a gift from Claus Nerlov (European Molecular Biology Laboratory, Heidelberg, Germany) ^38^. *Il1rn*-/- mice were a gift from Yoichiro Iwakura ^35^. For neutrophil reporter mice, we generated Catchup^IVM^ mice by crossing homozygous Catchup mice with homozygous Ai14 mice. For MK/platelet/neutrophil reporter mice, we generated Catchup^IVM^ x vWF-GFP mice by crossing homozygous Catchup^IVM^ mice with homozygous vWF-GFP mice. For MK/platelet reporter mice, we generated PF4-Cre x mT/mG mice by crossing hemizygous PF4-Cre mice with homozygous mT/mG mice. For MK/platelet mitochondria reporter mice, we generated PF4-Cre x PhAM^fl/WT^ mice by crossing hemizygous PF4-Cre mice with heterozygous PhAM^fl/WT^ mice. For MK/platelet-derived exosome reporter mice, we generated PF4-Cre x CD63-emGFP^l/s/l^ mice by crossing hemizygous PF4-Cre mice with homozygous CD63-emGFP^l/s/l^ mice. For MK depletion, we generated PF4-Cre x iDTR^fl/fl^ mice by crossing hemizygous PF4-Cre mice with homozygous iDTR^fl/fl^ mice ^58^.

### Animal experimentation

All procedures involving animals were approved by the Institutional Animal Care and Use Committee (IACUC) at Boston Children’s Hospital (protocol numbers #00002316, #00002319). All animal experiments were done in accordance with the NIH Guide for the Care and Use of Laboratory Animals.^91^ The initial approval process for animal experimentation ensures that all experiments were done in accordance with the 3Rs principles and that humane endpoints were defined and observed for all procedures. Euthanasia was performed by cervical dislocation of mice deeply anesthetized with isoflurane (3.5 % isoflurane, 1l/min O_2_). All mice were bred in local facilities at Boston Children’s Hospital. Mice were kept under a 12-hour light cycle with softwood bedding, nesting material, and ad libitum access to food and water under specific pathogen-free conditions. Sample sizes were not determined prior to experimentation because the expected effect size was unknown, but at least 6 animals (3F/3M) were included per experimental group in at least 3 replicates to ensure statistical robustness. Healthy male and female mice aged 8-12 weeks were included in experiments; no specific other inclusion or exclusion criteria were used. Mice and biological samples isolated from mice were randomly allocated to experimental groups, and whenever applicable, littermate mice served as controls. Whenever possible, investigators were blinded to outcome assessments by allocating mouse genotypes to outcome measures only after performing experiments and data analysis

### Antibodies

All antibodies listed are anti-mouse, unless otherwise stated. Dilutions are specified elsewhere. Anti-mouse CD11b (clone M1/70, #101212), CD144 (BV13, 138016), CD31 (MEC13.3, #102524), CD41 (MWReg30, #133904, #133912), CD42d (1C2, #148501), CD45 (S18009F, #157214), CD62L (MEL-14, #104420), CD61 (2C9.G2, #104308), CD62P (RMP-1, #148304), CD9 (MZ3, #124806), and Ly6G (1A8, #127612, #127608, #127614, #127606) were purchased from Biolegend (USA); anti-mouse PF4 (goat polyclonal, #AF595) from R&D Systems (USA); anti-mouse CD63 (EPR21151, #ab217345) and anti-EGFP (EPR14104, #ab183734) from Abcam (UK); anti-mouse vWF (rabbit polyclonal, #PA5-80223) from Thermo Fisher Scientific (USA); anti-rabbit IgG HRP (donkey polyclonal, #NBP1-75290) from Bio-Techne Corporation (USA), and anti-mouse CD42b (R300, #R300) and rat IgG isotype control (C301, #C301) from emfret Analytics GmbH & Co.KG (Germany). For all antibodies used in this work, validation statements are available via the providers’ websites.

### Reagents

*Activators/inhibitors:* G-CSF, CXCL1, and TNF were from PeproTech (USA); IL-1β was from R&D Systems (USA); LTB_4_, bleomycin, N-formyl-methionyl-leucyl-phenylalanine (fMLP), Phorbol 12-myristate 13-acetate (PMA), and GW4869 from Cayman Chemical (USA); LPS (O111:B4, E. coli) and diphtheria toxin (Corynebacterium diphtheriae, unnicked) from Merck (Germany). *Cell labeling/imaging reagents*: CellTracker^TM^ Green CMFDA, CellTracker^TM^ Red CMTPX, CellTracker^TM^ DeepRed, Draq5, DAPI, and Hoechst 33342 were from Thermo Fisher Scientific (USA); fluorophore-conjugated dextrans were from Biotium (USA) or Thermo Fisher Scientific (USA); and Evans Blue was from Merck (Germany).

### Immunoblotting

Cell lysates were prepared in 1x RIPA buffer (Thermo Fisher Scientific, USA) in the presence of 1x protease inhibitor (Merck, Germany) on ice. Cellular debris was spun down (10.000 g, 10 minutes, 4 °C), and supernatants were stored at -20 °C until analysis. Total protein content was determined using a BCA assay (Thermo Fisher Scientific, USA). Ten micrograms of the lysates were loaded onto 4-20 % gradient sodium dodecyl sulfate polyacrylamide (SDS-PAGE) gel electrophoresis gels (Bio-Rad Laboratories, USA). The separated proteins were transferred onto a 0.2 µm polyvinylidene difluoride (PVDF) membrane (GenScript Biotech, China). The membranes were blocked in a blocking buffer containing 5 % w/v dry milk powder (Merck, Germany) in 1x Tris-buffered saline (Boston BioProducts, USA) with 0.1% v/v Tween-20 (Merck, Germany) (TBS-T) for 2 hours at RT. The blocked membranes were then incubated with the primary antibodies at the appropriate dilution in blocking buffer overnight at 4 °C. The membranes were washed in 1xTBS-T, incubated with the secondary antibody at the appropriate dilution in blocking buffer for 1 hour at RT, washed again, and incubated with ECL chemiluminescent substrate (Bio-Rad Laboratories, USA). Chemiluminescence was visualized using the Invitrogen™ iBright™ FL1500 Imaging System (Thermo Fisher Scientific, USA).

### Flow cytometry and fluorescent cell sorting (FACS)

Whole blood samples, bone marrow samples, peritoneal lavage fluid samples, and lung samples were incubated with ACK buffer on ice to lyse erythrocytes. Samples were washed (500 g, 5 minutes, 4 °C), resuspended in flow cytometry buffer (0.5 % FBS, 2 mM EDTA, 1x PBS), and stained with fluorophore-conjugated antibodies (clones as listed above) at optimized dilutions (1:100 – 1:400) for 20 minutes at 4 °C in the dark. After washing, for flow cytometric analysis, samples were run on an LSR-Fortessa^TM^ instrument (Becton, Dickinson and Company, USA). For fluorescent cell sorting, samples were run on a MA900 Multi-Application Cell Sorter (Sony Biotechnology, USA). Unstained, single-stained, isotype control samples, and fluorescence minus one (FMO) control samples were used for compensation and gating. Voltage settings and gating strategies were kept constant throughout the study. Data analysis was performed using FlowJo software (version 10.0.8, Becton, Dickinson and Company, USA). Expression levels were quantified as median fluorescence intensity (MFI) and normalized to FMO or isotype control samples.

### Harvest and culture of megakaryocytes and neutrophils

Bone marrow was flushed from femora, tibiae, and iliac bones of 8-week-old WT, mT/mG, PF4-Cre x mT/mG, Catchup^IVM/Het^, and PF4-Cre x PhAM^fl/WT^ female and male mice, pooled, and passed through a 40 µm cell strainer to prepare a single-cell suspension. Erythrocytes were removed after lysis with ACK buffer, and remaining cells were resuspended in RPMI 1640 medium supplemented with 10% FBS, 10 mM HEPES, 1x sodium pyruvate, 1x non-essential amino acids, 1x GlutaMAX^TM^, and 1x penicillin/streptomycin (all from Gibco, Thermo Fisher Scientific, USA). For MK generation, bone marrow cells were cultured in complete RPMI medium supplemented with 2% conditioned medium from the TPO-producing fibroblast cell line GP122 ^92^ (hereafter referred to as TPO medium) for 5 days. MKs were then isolated using a two-step albumin gradient (3% and 1.5% bovine serum albumin in 1x PBS), as described ^93,94^. When stated, neutrophils were enriched by magnetic selection using neutrophil isolation kits (EasySep^TM^ Mouse Neutrophil Enrichment Kit, Stemcell Technologies, USA; Anti-Ly-6G MicroBeads mouse; Miltenyi Biotec, USA).

### Extracellular flux analysis of neutrophil metabolic function

Neutrophils were enriched from WT murine bone marrow using a negative isolation kit and co-cultured with live MKs (1 MK:10 neutrophils, emperipolesis) or separated from MKs by a 0.45µm transwell insert (controls) for 3 hours at 37 °C. When stated, co-cultures were treated with 10 µg/ml LPS or with 20 µM of the exosome inhibitor GW4869. Neutrophils were then re-isolated using a positive isolation kit, and their glycolytic and mitochondrial functions were assessed by real-time measurement of extracellular acidification rate (ECAR) and oxygen consumption rate (OCR), respectively, using the Seahorse XFe96 Extracellular Flux Analyzer (Agilent, USA). Briefly, neutrophils were resuspended in metabolic assay media (1x Dulbecco’s Modified Eagle’s Medium, 20 mM HEPES, pH 7.40) supplemented with 1 mM L-glutamine (ECAR) or with 1 mM L-glutamine, 10 mM D-glucose, and 1 mM sodium pyruvate (OCR) (all from Merck, Germany). Neutrophils were then seeded at 4×10^5^ cells per well in a 96-well assay plate (Agilent, USA), centrifuged (500 g, room temperature, no brake), and allowed to settle for ≥ 1 hours at 37 °C without CO_2_. ECAR was quantified after sequential treatment with: (1) 10 mM D-glucose, (2) 10 µM oligomycin A, and (3) 50 mM 2-deoxy-glucose, and OCR was quantified after: (1) 10 µM oligomycin A, (2) 5 µM carbonyl cyanide-4 trifluoromethoxy phenylhydrazone (FCCP), (3) 5 µM rotenone and 5 µM antimycin A (all from Merck, Germany). ECAR and OCR were measured three times before and after each treatment, with each experiment performed in three technical replicates. Glycolysis was calculated as: mean ECAR after D-glucose treatment – mean basal ECAR. Maximal respiration was calculated as: mean OCR after FCCP treatment – mean basal OCR.

### Neutrophil multiplexed cytokine release assay

Neutrophils were enriched from WT murine bone marrow using a negative isolation kit and co-cultured with live MKs (1 MK:10 neutrophils, emperipolesis) or separated from MKs by a 0.45µm transwell insert (controls) for 3 hours at 37 °C. Neutrophils were then re-isolated using a positive isolation kit, and stimulated with 10 µg/ml LPS for 20 minutes at 37 °C. Following stimulation, the neutrophils were gently pelleted by brief centrifugation (3 min, 250 g, 4 °C) and the cell culture supernatants were harvested and filtered using a 0.22 µm syringe-driven filter. The cell culture supernatant samples were immediately frozen to -80 °C and thawed on ice prior to analysis. Cytokine levels in the cell culture supernatant samples were measured using a commercially available, proximity extension assay-based multiplexed cytokine assay (Olink Proteomics, USA). We profiled 43 analytes using the Olink® Target 48 Mouse Cytokine Panel, following the manufacturer’s instructions. Of this broad panel, only analytes with detectable expression and known expression by neutrophils (13/43 analytes) were included in the final analysis. We calculated the fold-change in cytokine expression in the experimental samples relative to the expression in the paired control samples, followed by log2-normalization. We performed unsupervised hierarchical clustering of the log2-normalized relative cytokine expression data via the Ward D2 method and generated a heatmap using the pheatmap R-package (version: 1.0.12).

### Neutrophil ROS production assays via flow cytometry

Neutrophils were enriched from WT murine bone marrow using a negative isolation kit and co-cultured with live MKs (1 MK:10 neutrophils, emperipolesis) or separated from MKs by a 0.45µm transwell insert (controls) for 3 hours at 37 °C. The co-cultures were then loaded with 10 µM of the ROS indicator 5-(and-6)-chloromethyl-2’,7’-dichlorodihydrofluorescein diacetate (CM-H2DCFDA, Thermo Fischer Scientific, USA) for 15 mins at 37 °C, and when indicated, co-stimulated with 100 nM LTB_4_, 10 µM fMLP, or 50 nM PMA. ROS production was stopped by placing the incubation tubes on ice. Neutrophils were stained with a fluorophore-coupled *α*-Ly6G antibody (1:200) on ice for 20 mins, and the amount of ROS production was quantified as the MFI (DCF, 2,7-dichlorofluorescein) among Ly6G+ cells, normalized to the MFI (DCF) in unstimulated control neutrophils. Gating controls included samples without the ROS indicator.

### Neutrophil phagocytosis assay via flow cytometry

Neutrophils were enriched from murine WT bone marrow using a negative isolation kit and co-cultured with live MKs (1 MK:10 neutrophils, emperipolesis) or separated from MKs by a 0.45µm transwell insert (controls) for 3 hours at 37 °C. The co-cultures where then co-incubated with 10 µg/ml pHrodo^TM^ Deep Red E. coli Bioparticles^TM^ Conjugate (Thermo Scientific, USA) for 30 mins at 37 °C, and when indicated, co-stimulated with 100 nM LTB_4_, 10 µM fMLP, or 50 nM PMA. Phagocytosis was stopped by placing the incubation tubes on ice. Neutrophils were stained with a fluorophore-coupled *α*-Ly6G antibody (1:200) for 20 mins on ice, and the amount of phagocytosis was quantified as the % E. coli particles+ among Ly6G+ cells, and as the MFI (E. coli bioparticles) among Ly6G+ cells, normalized to the MFI (E. coli bioparticles) in unstimulated control neutrophils. Negative biological controls included samples co-incubated with E. coli particles on ice, and samples without E. coli particles.

### Neutrophil phagocytosis assay via microscopy

Bone marrow cells were isolated from Catchup^IVM/Het^ bone marrow (harboring TdTomato+, TdT+, neutrophils) and co-cultured with MKs grown from WT mice (1 MK:10 neutrophils, emperipolesis) for 3 hours at 37 °C in the presence of anti-mouse CD41-AF488 (1:100). The co-cultures were then co-incubated with 25 µg/ml pHrodo^TM^ Deep Red E. coli Bioparticles^TM^ Conjugate (Thermo Scientific, USA) for 60 mins at 37 °C, and when indicated, co-stimulated with vehicle control, 10 µg/ml LPS, or 10 µM fMLP. Cultures were imaged with a Nikon Ti2 inverted microscope with a Yokogawa W1 spinning disk, using a 40x objective. Per condition in each replicate, at least 3 randomly selected field of views (FOVs) were imaged, and analyzed using Fiji (version 2.0.0)^95^. Briefly, neutrophils were identified as TdT+ cells, and then classified by co-staining with anti-mouse CD41 and the uptake of Deep Red E. coli bioparticles. Negative controls included conditions without MKs, without anti-CD41, or without E. coli bioparticles.

### Neutrophil cell death rate assays

WT murine bone marrow cells were co-cultured with live MKs (1 MK:20 bone marrow cells, emperipolesis) or separated from MKs by a 0.45µm transwell insert (controls) for 4 hours at 37 °C. Neutrophils were then enriched from the cultures using a negative isolation kit, and cultured in a 96-well plate at a density of 4×10^5^ cells/well for up to 72 hours in complete RPMI. Where indicated, neutrophils were treated with the ferroptosis inhibitor Ferrostatin-1 at a concentration of 10 µM (Merck, Germany). At the specified time points, neutrophils were harvested, washed in 1x HBSS (350g, 5 mins, 4 °C), and stained with the following reagents for 20 minutes at 4 °C in the dark in a buffer containing 1x HBSS, 5% FBS, and 2 mM Ca^2+^: anti-mouse Ly6G-APC/Cy7 (1:400), Annexin V-Pacific Blue (Biolegend, USA; 1:200), NucSpot® Far-Red (Biotium, USA, 1x), FerroOrange (Merck, Germany, 1 µmol/L), and CellEvent™ Caspase-3/7 Detection Reagent Green (Thermo Fisher Scientific, USA, 1:100). Cell death rates were assessed by flow cytometry, where necrotic neutrophils were measured as the % Annexin V+NucSpot+ of Ly6G+ cells, apoptotic neutrophils were measured as the % Annexin V+ CellEvent™+ cells of Ly6G+ cells, free Fe^2+^ ions were measured as MFI(FerroOrange) among Annexin V+NucSpot+LyG6+ cells, and ferroptosis as the % Annexin V+NucSpot+ cells of Ly6G+ cells in the presence of 10 µM Ferrostatin-1.

### Neutrophil trans-well migration assay

WT murine bone marrow cells were cultured overnight alone (control), in 0.45 µm-filtered MK culture supernatant (control), with live MKs (1 MK:10 neutrophils, emperipolesis), with MK-derived exosomes (isolated by ultracentrifugation as described)^96^, or with the exosome generation inhibitor 20 µM GW4869, as indicated. For migration assays, the lower chambers of a 24-well plate well were filled with 600 µl of TPO medium containing 100 nM LTB_4_ or vehicle control. Trans-well inserts with 3 µm pores (Corning, USA) were added, and 200 µl of bone marrow cell suspensions were added to the inserts. After incubation for 2 hours at 37 °C, cells were collected from the supernatant of the upper and lower chambers, and Ly6G+ neutrophils were quantified by flow cytometry using 15 µm ∅∅ counting beads (Polysciences, USA).

### Neutrophil trans-megakaryocyte chemotaxis assay

Neutrophil trans-MK migration was imaged using µ-slide chemotaxis chambers (Ibidi GmbH, Germany). Briefly, MKs in TPO medium were mixed with 1.5% bovine collagen I (Thermo Fisher Scientific, USA) and stained with AF488-coupled anti-CD41 (1:100). Six µl of this MK matrix was instilled into the middle chamber of the µ-slide chemotaxis chamber and incubated at 37 °C for 60 minutes to allow matrix gelation. The chamber on the right was filled with 75 μl of Catchup^IVM/Het^ murine bone marrow cell suspension in TPO medium, harboring TdT+ neutrophils, and that on the left was filled with 75 μl of TPO medium. To establish a chemotaxis gradient, 37.5 µl of 100 nM LTB_4_ medium was added to the left chamber prior to imaging, which was performed on a Nikon Ti2 inverted microscope with a Yokogawa W1 spinning disk, using a 10x objective. Time-lapse images were acquired every 1-2 minutes for 2 hours, with 2 FOVs, ∼30 Z-stacks, and 2.5 µm per stack. Images were analyzed using Imaris software (version 10.1, Bitplane, UK). Briefly, to analyze emperipolesis events, MKs were surface-rendered and masked to visualize internalized neutrophils and migrating neutrophils were automatically tracked to measure migration speed before and after emperipolesis events or, as control events, before and after MK surface contact without undergoing emperipolesis. In alternate experiments, the migration speed of CD41+ vs. CD41- neutrophils was tracked.

### Neutrophil rolling on endothelial cells

Murine B.End5 endothelial cells ^97^ (ECACC Cat# 96091930, RRID:CVCL_2252) were grown to confluence in glass-bottomed FluoroDish cell culture dishes (World Precision Instruments, USA) and stimulated with TNF (20 ng/ml) for 2 hours. Neutrophils were enriched from WT bone marrow using a negative isolation kit, stained with fluorophore-coupled anti-Ly6G (1:200) and anti-CD41 antibodies (1:100), and added to the endothelial cells for live cell imaging. In other experiments, B.End5 cells were grown in a flow chamber (µ-slide I Luer, Ibidi GmbH, Germany) and activated with TNF as described above. Neutrophils enriched from WT murine bone marrow were aspirated into a peristaltic pump tubing system, connected to the flow chamber. A flow rate of 300 to 500 µl/min was applied. Cells were observed by spinning disc confocal microscopy or widefield microscopy (Nikon Ti2 inverted microscope) and images were analyzed using Imaris software (version 10.1, Bitplane, UK).

### Neutrophil adoptive transfer

Bone marrow cells were collected from WT mice and divided equally per animal. One half was stained with 10 µM CellTracker^TM^ DeepRed or Red and co-cultured with MKs (1 MK:20 marrow cells, emperipolesis), while the other half was stained with 10 µM CellTracker^TM^ Green CMFDA and cultured separated from MKs by a 0.45µm transwell insert (controls) for 3 hours. WT mice were treated i.p. with 2 mg/kg LPS or 1 µg/kg IL-1*β*. Thirty minutes after induction of peritonitis, the cultured bone marrow cells were remixed and injected retro-orbitally into the pre-treated mice. After 1 hour, retro-orbital bleeding and peritoneal lavage were performed. For peritoneal lavage, the peritoneal cavity was rinsed twice with 10 ml of ice-cold buffer (1x PBS containing 2 % BSA, 2 mM EDTA). The count of Ly6G+ CellTracker^TM^ DeepRed or Red and Green CMFDA+ neutrophils in the circulation and the peritoneal cavity was analyzed by flow cytometry using 15 µm ∅∅ counting beads (Polysciences, USA).

### Platelet depletion

PF4-Cre^+^ x CD63-emGFP^l/s/l^ experimental mice and PF4-Cre^-^ x CD63-emGFP^l/s/l^ littermate controls received daily i.v. injections of 2 mg/kg of anti-mouse CD42b (clone R300) or rat IgG isotype control (clone C301) in 150 µL 1x PBS (emfret Analytics GmbH&Co.KG, Germany) via the RO route for 3 consecutive days, to selectively deplete circulating platelets while maintaining bone marrow MKs ^46–48^. The efficiency of platelet depletion was monitored by quantifying the count of circulating CD41+ platelets by flow cytometry using 1 µm ∅∅ counting beads (Polysciences, USA). MK sufficiency was confirmed by femur histology.

### Megakaryocyte depletion

PF4-Cre^+^ x iDTR^fl/fl^ experimental mice and PF4-Cre^-^ x iDTR^fl/fl^ littermate controls received daily i.p. injections of 250 ng diphtheria toxin in 100 µl ddH_2_O for 3 consecutive days to selectively ablate bone marrow MKs while maintaining normal platelet counts or for 6 days to ablate bone marrow MKs and platelets ^47,58^. MK depletion was confirmed by femur histology.

### Measurement of optical redox ratios in neutrophils in MK-depleted mice

After MK ablation, bone marrow and blood cells were harvested, and single-cell suspensions were stained with a live/dead stain and fluorophore-coupled antibodies against CD45, CD11b, and Ly6G in 1xFACS buffer (as outlined above), washed once, and resuspended in metabolic assay media (1x Dulbecco’s Modified Eagle’s Medium, 20 mM HEPES, pH 7.40) supplemented with either 1 mM L-glutamine (to study the glycolytic potential) or with 1 mM L-glutamine, 10 mM D-glucose, and 1 mM sodium pyruvate (to study the respiratory potential) (all from Merck, Germany). Where indicated, cells were stimulated with 10 µg/mL LPS for 10 minutes at 37 °C. To determine the glycolytic potential of neutrophils, the cells were then treated sequentially for 5 minutes each at 37 °C with: (1) vehicle, (2) 10 mM D-glucose, and (3) 50 mM 2-deoxy-glucose. To determine the respiratory potential of neutrophils, the cells were treated with: (1) vehicle, (2) 5 µM carbonyl cyanide-4 trifluoromethoxy phenylhydrazone (FCCP), (3) 5 µM rotenone and 5 µM antimycin A (all from Merck, Germany). Optical redox ratios (ORRs) were then determined via flow cytometry as per published protocols ^68^. Briefly, the levels of NADH and FAD among live CD45+CD11b+Ly6G+ neutrophils were assessed by their autofluorescence in the BV421 and AmCyan channels, respectively. Raw MFIs were used to calculate ORRs as follows: ORR = MFI(FAD) / [MFI(NADH) + MFI(FAD)]. ORR values were normalized to the median ORR in the vehicle-treated control (MK-sufficient) group.

### Measurement of glucose uptake by neutrophils in MK-depleted mice

After MK ablation, bone marrow and blood cells were harvested, and single-cell suspensions were stained with a live/dead stain and fluorophore-coupled antibodies against CD45, CD11b, and Ly6G in 1xFACS buffer (as outlined above), washed once, and resuspended in metabolic assay media (1x Dulbecco’s Modified Eagle’s Medium, 20 mM HEPES, pH 7.40) supplemented with either 0 mM glucose or with 10 mM glucose as a control. The cells were stimulated with 10 µg/mL LPS or a vehicle control for 15 minutes at 37 °C. Glucose uptake was then determined using a glucose uptake capacity assay kit (Dojindo Molecular Technologies, USA), as per the manufacturer’s instructions. Briefly, the cells were incubated with 1x Glucose Uptake Probe-Green for 15 minutes at 37 °C, followed by washing the cells twice in 1x ice-cold washing solutions. Until analysis, the cells were kept in 1x ice-cold washing solution on ice in the dark and processed within 1 hour. The uptake of the Glucose Uptake Probe-Green in neutrophils was determined via flow cytometry by assessing the MFI(Glucose Uptake Probe-Green) in the FITC-channel among live CD45+CD11b+Ly6G+ neutrophils.

### Neutrophil bone marrow egress in megakaryocyte-depleted mice

After MK ablation, mice were injected with either 300 ng CXCL1 i.v. or 2.5 µg G-CSF i.v. to induce neutrophil egress from the bone marrow. Blood samples were collected retro-orbitally before treatment and at specified time points after treatments. Bone marrow cells were harvested after the treatment as described above, and the number of Ly6G+ neutrophils in the marrow and circulation was quantified by flow cytometry with 15 µm ∅∅ counting beads (Polysciences, USA). Femora were collected for histology to confirm MK depletion and quantify emperipolesis as described below.

### Pseudomonas aeruginosa pneumonia in megakaryocyte-depleted mice

*Pseudomonas aeruginosa* pneumonia was induced as described previously in detail elsewhere ^98^. Briefly, *Pseudomonas aeruginosa* (PsA, strain PA103, obtained from ATCC) were cultured from a frozen stock on a TSB/agar plate and grown 16–24 h at 37 °C, followed by selection of a single colony of PsA from the TSB/agar plate and inoculation into 3 mL of sterile tryptic soy broth (TSB) in a 15 mL Falcon tube. Next, PsA were grown 16–20 h in a heated shaker at 250 RPM, 37 °C, then washed 3x and quantified using spectrophotometry (after prior titration) immediately prior to inoculation. Following MK ablation as described, mice were inoculated with PsA at a non-lethal dose (3×10^5^ CFU, diluted to desired concentration with sterile PBS 1x) via the intratracheal route in a volume of 75 µL of 1x PBS, or with 1x PBS only as a vehicle control. Where indicated, mice were pre-treated 24 hours prior to PsA inoculation with 100 µg i.p. of a neutrophil-depleting antibody (clone 1A8, Bio X Cell Inc., USA). Tissues were harvested 4 hours after inoculation. To confirm the absence of bacteremia and systemic spread of PsA at this timepoint, blood, spleen and liver homogenate samples were analyzed. Whole livers (without the gall bladder) and spleens were removed post mortem and homogenized in PBS with 0.05% Triton X-100 to release any potential intracellular bacteria. Colony-forming units (CFU) in the blood, liver or spleen were determined by plating 3 serial dilutions of the blood or tissue suspension on TSB/agar and incubated for 18 hours at 37 °C. Resulting quantities were normalized to organ weight prior to homogenization. For cell count quantification via flow cytometry, blood was drawn via cardiac punctures, marrow was collected via flushing of the femur cavity, and lung cells were harvested by digesting lungs for 30 minutes at 37° C in 1640 RPMI containing 2 mg/mL Collagenase II (Merck, Germany) and 0.1 mg/ml DNAse I (Roche, Switzerland), followed by passing of the digested lung through a 40 µm cell strainer to prepare a single-cell suspension. The number of CD45+CD11b+Ly6G+ neutrophils and CD45+CD11b+Ly6C-F4/80+ macrophage-enriched myeloid cells was quantified by flow cytometry with 15 µm ∅∅ counting beads (Polysciences, USA).

### Histological lung assessment of Pseudomonas aeruginosa infected mice

For lung histology, lung samples were fixed in 4% PFA/1x PBS for 1 day, dehydrated in 70 % ethanol for 1 day, and embedded in paraffin. Five µm lung sections were stained with H&E. To quantify lung inflammation, whole sections were imaged with a slide scanner (EVOS M7000, Thermo Fisher Scientific, USA) at 200x magnification, and entire sections were scored employing an adapted published lung inflammation sum score, consisting of the following three components: alveolar congestion, neutrophil infiltration, and hyaline membrane formation [0-4 each, sum score ranging from 0-12, with 0 meaning no damage and 12 maximal damage, median values were recorded] ^99^.

### Multiplexed cytokine assay of plasma and lung samples of Pseudomonas aeruginosa infected mice

To quantify cytokine levels, plasma samples were collected by spinning blood samples (1300g, 7 min, RT) and lung homogenates were prepared from lung samples using an mechanical homogenizer while keeping the samples in 1x tissue extraction buffer (T-PER™, Tissue Protein Extraction Reagent, Thermo Fisher Scientific, USA). Samples were frozen to - 80 °C immediately after collection and thawed on ice prior to analysis. Cytokine levels were measured using a commercially available, proximity extension assay-based multiplexed cytokine assay (Olink Proteomics, USA). We profiled 43 analytes using the Olink® Target 48 Mouse Cytokine Panel, following the manufacturer’s instructions. Of this panel, only analytes with detectable expression were included in the final analysis. We performed unsupervised hierarchical clustering of the z-score -normalized cytokine expression data via the Ward D2 method and generated a heatmap using the pheatmap R-package (version: 1.0.12).

### IL-1*β*-induced pneumonitis in megakaryocyte-depleted mice

After MK ablation, mice underwent intranasal installation of 1 µg/kg IL-1*β*, and tissues were harvested after 2 hours. For cell quantification, blood was drawn via cardiac punctures, marrow was collected via flushing of the femur cavity, and lung cells were harvested by digesting lungs for 30 minutes at 37° C in 1640 RPMI containing 2 mg/mL Collagenase II (Merck, Germany) and 0.1 mg/ml DNAse I (Roche, Switzerland), followed by passing of the digested lung through a 40 µm cell strainer to prepare a single-cell suspension. The number of CD45+CD11b+Ly6G+ neutrophils cells was quantified by flow cytometry with 15 µm ∅∅ counting beads (Polysciences, USA).

### LPS-induced peritonitis in megakaryocyte-depleted mice

After MK ablation, mice underwent intraperitoneal injection of 2 mg/kg LPS, and tissues were harvested after 2 hours. For cell quantification, blood was drawn via cardiac punctures, marrow was collected via flushing of the femur cavity, and peritoneal extravasate was collected by flushing the peritoneal cavity twice with 2x 10 ml of ice-cold 1xFACS buffer, followed by passing of lavage fluid through a 40 µm cell strainer to prepare a single-cell suspension. The number of CD45+CD11b+Ly6G+ neutrophils cells was quantified by flow cytometry with 15 µm ∅∅ counting beads (Polysciences, USA).

### Neutrophil migration in IL-1*β*-induced peritonitis

WT or PF4-Cre^+^ x mT/mG MK/platelet membrane reporter mice and PF4-Cre^-^ x mT/mG littermate controls underwent intraperitoneal injection of 1 µg/kg IL-1*β*. After 2 hours, omentum tissue samples were collected in 4% PFA/1x PBS for histology and peritoneal extravasate was collected as described for confocal micrscopy and flow cytometry.

### Neutrophil migration in LPS-induced peritonitis

PF4-Cre^+^ x CD63emGFP MK/platelet exosome reporter mice and PF4-Cre^-^ x CD63emGFP littermate controls underwent peripheral platelet depletion with daily i.v. injections of 2 mg/kg of anti-mouse CD42b (clone R300) for 72 hours as described. Platelet depletion was confirmed by flow cytometry with 1 µm ∅∅ counting beads (Polysciences, USA). Platelet-depleted mice then were subjected to intraperitoneal injection of 2 mg/kg LPS. After 2 hours, blood samples were collected via retro-orbital bleeding, and peritoneal extravasate was collected as described for confocal microscopy and flow cytometry.

### Histological quantification of emperipolesis in mouse models of inflammatory diseases. K/BxN arthritis

150 µl of pooled serum collected from 9-11 week-old K/BxN mice were transferred i.p. to WT recipient mice, on day 0 and day 2, as described elsewhere.^84^ Femora were harvested 7 to 8 days after induction. *Cecal ligation and puncture (CLP) model of polymicrobial sepsis:* A small incision was made in the lower abdominal area to expose the cecum. The cecum was ligated with a 3.0 silk suture and then carefully punctured a single time with an 18-gauge needle, allowing the release of fecal contents into the peritoneal cavity and causing polymicrobial sepsis, as described elsewhere ^100^. Control animals underwent sham surgery without cecal puncture. Femora were harvested 4 days after surgery. *Systemic sclerosis:* 100 µg of bleomycin in 1x PBS was injected intradermally into the dorsal skin of female WT mice daily, 5 times a week, for 4 weeks, as described ^33,34^. Femora were harvested on day 28 of treatment. *LPS-induced peritonitis*: WT mice were injected i.p. with 2 mg/kg LPS in 1x PBS, and femora were harvested after 2 hours. *Pseudomonas aeruginosa pneumonia*: WT were inoculated with *Pseudomonas aeruginosa* (*PsA,* strain PA103) at a non-lethal dose (3×10^5^ CFU) via the intratracheal route in a volume of 75 µL of 1x PBS, or with 1x PBS only as a vehicle control, and femora were harvested after 4 hours. *IL-1β-induced pneumonitis*: WT mice underwent intranasal installation of 1 µg/kg IL-1*β*, and femora were harvested after 2 hours. *All animal models*: Harvested femora were fixed in 4% PFA/1x PBS for 2 days, dehydrated in 70 % ethanol for 2 days, decalcified with Decalcifying Solution-Lite (Merck, Germany) for 2 days and embedded in paraffin. 5 µm bone marrow sections were stained with H&E. For quantification of emperipolesis, whole sections were imaged with a slide scanner (EVOS M7000, Thermo Fisher Scientific, USA) at 200x magnification, at least 20 randomly selected FOVs containing at least 300 individual MKs were analyzed for the % of MKs containing at least 1 neutrophil, and median values were recorded.

### Whole mount bone marrow histology

Whole mount bone marrow preparation, staining, and imaging were performed as described elsewhere ^88^. Briefly, mice were injected i.v. with 5 µg AF647-coupled anti-CD31/CD144 and perfused with 4% PFA/1x PBS after 15 minutes. Femora and tibiae were harvested, postfixed in 4% PFA/1x PBS for 30 minutes at room temperature, cryopreserved in sucrose, frozen in O.C.T. compound (Thermo Fisher Scientific, USA) at -20°C, and shaved on a cryostat to expose the marrow cavities. Bones were then incubated in 1x PBS with 10% FBS, 0.5% Triton X-100, AF594-coupled anti-Ly6G (1:100), AF488-coupled anti-CD41 (1:100), and Hoechst 33342 for 2 days. Images were acquired on a Zeiss LSM 800 confocal microscope, and reconstructed in 3D using Imaris software (version 10.1, Bitplane, UK).

### Confocal microscopy

4%-formalin or ice-cold methanol fixed cytospun cells or tissues were permeabilized and blocked in a buffer of 1xPBS containing 0.1% Tween® 20 (Merck, Germany), 5% FBS, 1% w/v BSA (Merck, Germany), and 0.3 M glycine (Merck, Germany) for 1 hour at room temperature, then incubated with primary antibodies (1:100) in a buffer of 1xPBS containing 0.1% Tween® 20, 1% w/v BSA, and 0.3 M glycine overnight (4°C, dark). After 5x washing with 1xPBS, secondary antibodies (1:200) in a buffer of 1xPBS containing 0.1% Tween® 20, 1% w/v BSA, and 0.3 M glycine were added for 1 hour at room temperature. Draq5, Hoechst 33342 or DAPI was added during the last 15 minutes, as indicated. After 5x washing in 1xPBS, cells and tissues were mounted with FluorSave™ Reagent (Merck, Germany). Imaging was performed with a Nikon C1 Plus or Zeiss LSM 800 confocal microscope.

### Electron microscopy

Cells or decalcified bones were fixed in 0.1 M sodium cacodylate buffer (pH 7.4) with 2.5% glutaraldehyde and 2.5% paraformaldehyde overnight at 4°C. Samples were postfixed in 1% osmium tetroxide and 1.5% potassium ferrocyanide for 1 hour at room temperature, washed in maleate buffer, and incubated in 1% uranyl acetate for 30 minutes. After dehydration in ethanol, samples were infiltrated in a 1:1 mixture of propylene oxide and Epon resin (TAAB Laboratories Equipment Ltd, UK), then embedded in Epon resin and polymerized at 60°C for 2 days. Sections (50 nm) were imaged using a JEOL 1200EX electron microscope.

### Intravital two-photon microscopy of calvarial marrow

Mice were anesthetized with 2% isoflurane, and immobilized in a head holder adapter (SGM4, Narishige). The skin over the calvarium was incised from the parietal to the frontal bone, and the periosteum was removed with scissors. A custom-made skull stabilizer was attached to the exposed calvaria and a drop of 1x PBS was placed between the objective and the skull. A 60 µm thick area of the skull marrow (20-30 Z-stacks, 2-3µm per stack) was imaged at 30 second intervals for 30 minutes. For quantification of emperipolesis, bone autofluorescence was eliminated by generating an SHG-based surface and masking it, and MKs were surface-rendered and masked to visualize internalized neutrophils, using Imaris software (version 10.1, Bitplane, UK).

### Stable isotope labeling by amino acids in cell culture (SILAC) proteomics to identify proteins transferred from megakaryocytes to neutrophils

The peptides of murine marrow-derived megakaryocytes were labeled with ^13^C_6_ ^15^N_2_ L-lysine and ^13^C_6_ ^15^N_4_ L-arginine by culturing mouse marrow cells in RPMI 1640 medium, supplemented with 10% dialyzed FBS, 50 mg/L ^13^C_6_ ^15^N_2_ L-lysine, 50 mg/L ^13^C_6_ ^15^N_4_ L-arginine, 200 mg/L L-proline (all Thermo Fisher Scientific), and 2% conditioned medium from the TPO-producing fibroblast cell line GP122 ^92^. Following validation of labeling efficiency via mass spectrometry, labeled MKs were then co-cultured with unlabeled murine marrow neutrophils for 3 hours, or co-cultured with unlabeled neutrophils separated by 0.45 µm trans-well inserts as a control. Cultured murine neutrophils were FACS-sorted as life, single, CD45+CD11b+Ly6G+ cells (>0.5 x 10^6^ / replicate). Sorted neutrophils were lysed by resuspending washed cells in 1x RIPA buffer (Thermo Fisher Scientific, USA) in the presence of 1x protease inhibitor (Merck, Germany). Per replicate, 25 µg of protein content in the neutrophil lysates were separated via SDS-PAGE and ratios of labeled versus unlabeled peptide ratios were identified by mass spectrometry. The identified labeled peptides were analyzed using the publicly accessible Protein Analysis THrough Evolutionary Relationships (PANTHER) resource (https://www.pantherdb.org/, version: 19.0) ^52^, using the PANTHER overrepresentation test tool (released 2024-08-07), assessing the GO cellular component complete database (version: 21836) from the GO ontology database (version: DOI: 10.5281/zenodo.15066566, released 2025-03-16), the PANTHER Pathway Ontology (version: 19.0, released 2024-06-20), and as a reference list, the genome of *Mus musculus* provided in the PANTHER database (version: 19.0).

### Neutrophil transcriptome analysis

For transcriptome analysis of neutrophils following co-culture with MKs, murine marrow neutrophils were cultured with MKs as described, washed twice in a buffer containing 1x PBS, 0.5 % w/v BSA and 2 mM EDTA (350g, 5 mins, 4 °C), and FACS-sorted as life, single, CD45+CD11b+Ly6G+ cells (>1×10^6^ / replicate). For transcriptome analysis of MK-depleted mice, PF4-Cre^+^ x iDTR^fl/fl^ experimental mice and PF4-Cre^-^ x iDTR^fl/fl^ littermate controls were treated with diphtheria toxin for three days as described and treated with vehicle or LPS (2mg/kg) i.p. to induce systemic inflammation. After 2 hours, neutrophils were isolated from the marrow as described or from EDTA-anticoagulated blood samples obtained via cardiac punctures. Red blood cells in marrow and blood samples were lysed with 1x ACK buffer on ice and cells were washed in a buffer containing 1x PBS, 0.5 % w/v BSA and 2 mM EDTA (350g, 5 mins, 4 °C). Marrow and blood neutrophils were FACS-sorted as life, single, CD45+CD11b+Ly6G+ cells (>0.2×10^6^ / replicate). From all samples, RNA was isolated using a Direct-zol RNA Miniprep Kit (Zymo Research Corp, USA) in a single batch, following the manufacturer’s instruction, and RNA was eluted in 25 µl of water. RNA quality (RIN values) was assessed using a 2100 Bioanalyzer instrument (Agilent, USA) in house. 10 ng samples of RNA (RIN>7) were sent for low-input bulk RNA seq-library preparation at the Broad Institute Clinical Labs, using the Smart-seq2 technique^101^, followed by 2×38bp paired-end sequencing on an Illumina® next-generation sequencing platform. Initial quality control of the raw sequencing data for read number, read length, and GC content was done using the MultiQC tool (v1.26) ^102^. Quality trimming was performed using trimmomatic (v0.39) ^103^ with quality thresholds set to 20. Pseudo-mapping and count quantification was performed using Salmon (v1.10.2) ^104^. To this end, a transcriptome index was created based on GRCm39 reference transcriptome (Ensemble, release 113) ^105^. Transcript abundances were aggregated to gene-level count matrices using tximeta (v3.20) ^106^. Differential gene expression analysis was performed using DSeq2 (v3.20) (Wald test, FDR: Padj<0.10) ^107^. Gene Ontology (GO) term enrichment and KEGG term enrichment was done using clusterProfiler (v4.14.3) (FDR: Padj<0.10) ^108,109^. To this end, GO term annotation was based on org.Mm.eg.db (v3.20.0) and KEGG annotations were retrieved online via clusterProfiler.

### Neutrophil proteome analysis

For proteome analysis, cultured murine neutrophils were FACS-sorted as life, single, CD45+CD11b+Ly6G+ cells (>0.5 x 10^6^ / replicate). Sorted neutrophils were lysed by resuspending washed cells in 1x RIPA buffer (Thermo Fisher Scientific, USA) in the presence of 1x protease inhibitor (Merck, Germany). Neutrophil proteins were precipitated using a TCA protein precipitation kit (ProteoExtract® Protein Precipitation Kit, Merck, Germany), following the manufacturer’s instruction. The neutrophils protein content was then identified via mass spectrometry. Results were analyzed for differentially abundant proteins using proDA (version 1.20.0) ^110^ and for enriched pathway clusters based on mouse KEGG pathway terms using PathfindR (version 2.4.1) ^111^.

### Statistical analysis

Unless otherwise stated, statistical comparisons were done as follows. All statistical tests were two-tailed. The Mann-Whitney U test was used to assess differences in medians between two groups. For paired samples between two groups, the Wilcoxon matched-pairs test was used. For comparisons involving more than two groups, the Kruskal-Wallis test was used, followed by a post-hoc Dunn’s test for multiple comparisons. When analyzing multiple groups with paired samples, the Friedman test was applied, followed by false discovery rate (FDR) correction (α<0.05) to adjust for multiple comparisons. To compare proportions, the Yates’ continuity-corrected chi-square test was used. Unless otherwise noted, data points represent individual observation units, such as individual animals or experimental conditions, and central values and errors are presented as medians and interquartile ranges. Statistical analyses were performed using GraphPad Prism software (version 10, Dotmatics, USA) or R (version 4.4.2, R Foundation for Statistical Computing, Austria). *P*-values are reported as follows: \**P* ≤ .05; \*\**P* ≤ .01; \*\*\**P* ≤ .001, **** *P* ≤ .0001. *P*-values > 0.5 were considered not significant.

### Data and material availability

Raw data and material are available upon reasonable request. Inquires may be directed to the corresponding author.

## ACKNOWLEDGEMENTS

We thank Teresa Bowman of the Specialized Histopathology Core at Dana-Farber / Harvard Cancer Center for histopathology services. We thank Dr. Ross Tomaino of the Taplin Mass Spectrometry Facility at the Harvard Medical School for mass spectrometry sample processing. We gratefully acknowledge the MicRoN (Microscopy Resources on the North Quad) core at Harvard Medical School for imaging support. Metabolism studies were supported by the Enders Cell Function and Imaging Core at Boston Children’s Hospital (National Institute of Health P30 Harvard Digestive Disease Center grant DK034854).

## AUTHOR CONTRIBUTIONS

Conceptualization: J.K.K., P.C., P.A.N.; Methodology: D.G., O.B., L.G., A.We., S.J.C., M.G., Y.I., D.P.H.v.K., J.E.I., M.R.L., W.B.; Investigation: J.K.K., P.C., G.H.B., R.D., F.Y.H., R.M.H., J.E.S., D.M., S.B., A.Wa.; Writing - Original Draft: J.K.K., P.C., P.A.N.; Writing - Review & Editing: J.K.K., P.C., G.H.B., R.D., O.B., L.G., S.J.C., M.G., D.P.H.v.K., M.R.L., E.B., P.A.N.; Funding acquisition: P.A.N.

## EXTENDED DATA FIGURES AND LEGENDS

**Figure S1.**
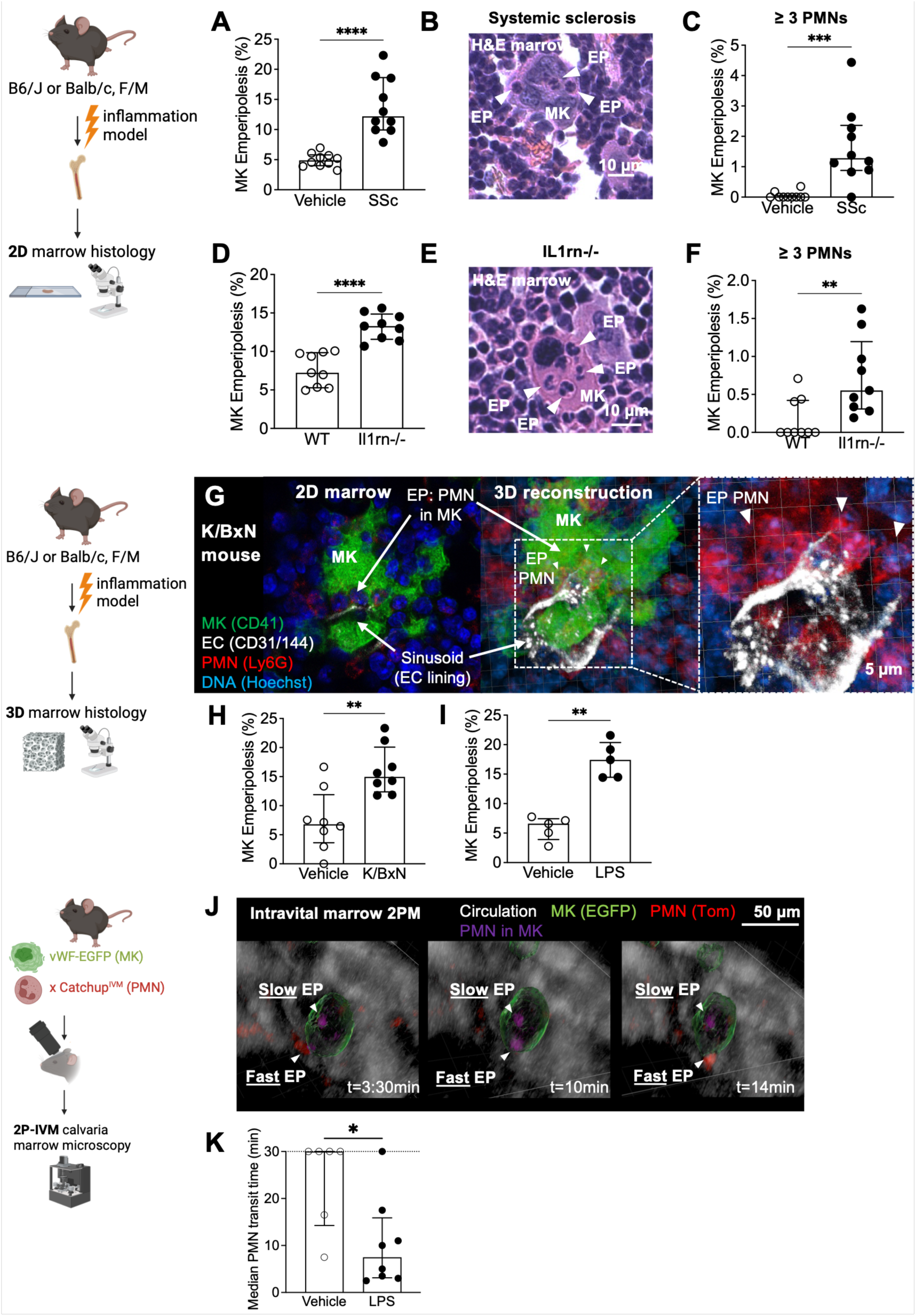
Enhanced emperipolesis is conserved across inflammatory pathologies. (A-G) Marrow MK EP frequency involving ≥1 (A,D) or ≥3 neutrophils (C,F) in mouse inflammation models. (A-C) Bleomycin-induced systemic sclerosis (SSc) vs vehicle (28 days; n=10/10). (D-F) Il1rn^-/-^ inflamed mice vs BALB/cJ wild-type controls (10 weeks; n=9/9). (B,E) H&E-stained marrow sections in SSc (B) and Il1rn^-/-^ mice (E), arrowheads: internalized neutrophils. (G-I) Whole-mount confocal microscopy of marrow from K/BxN arthritic mice (G,H, 7-8 days vs vehicle, n=8/8) and LPS-treated mice (I, 2 mg/kg i.p., 2 hours vs vehicle, n=5/5). (G) Left, 2D Z-stack, right and inset, 3D reconstruction; MKs anti-CD41 (green), neutrophils anti-Ly6G (red), endothelium (EC) anti-CD31/CD144 (gray), DNA Hoechst 33342 (blue). Arrowheads: EP events. (H,I) EP frequency quantified in 3D-reconstructed whole-mounts. (J,K) Two-photon intravital microscopy of calvarial marrow in Catchup^IVM^ × vWF-EGFP mice; neutrophils tdTomato (red; magenta when internalized), MKs EGFP (green), vasculature Evans Blue (gray). (J) Arrowheads: Simultaneous fast and slow EP within one MK. (K) Neutrophil transit time after LPS (2 mg/kg, i.p., 2 hours; n=8, 35 events) vs vehicle (PBS; n=6, 16 events). (A,C,D,F,I,K) Bars, median; error bars, IQR. Statistics: (A,C,D,F,H,I,K) Mann-Whitney U tests. Significance: *P ≤ 0.5, **P ≤ 0.01, ***P ≤ 0.001, ****P ≤ 0.0001.

**Figure S2.**
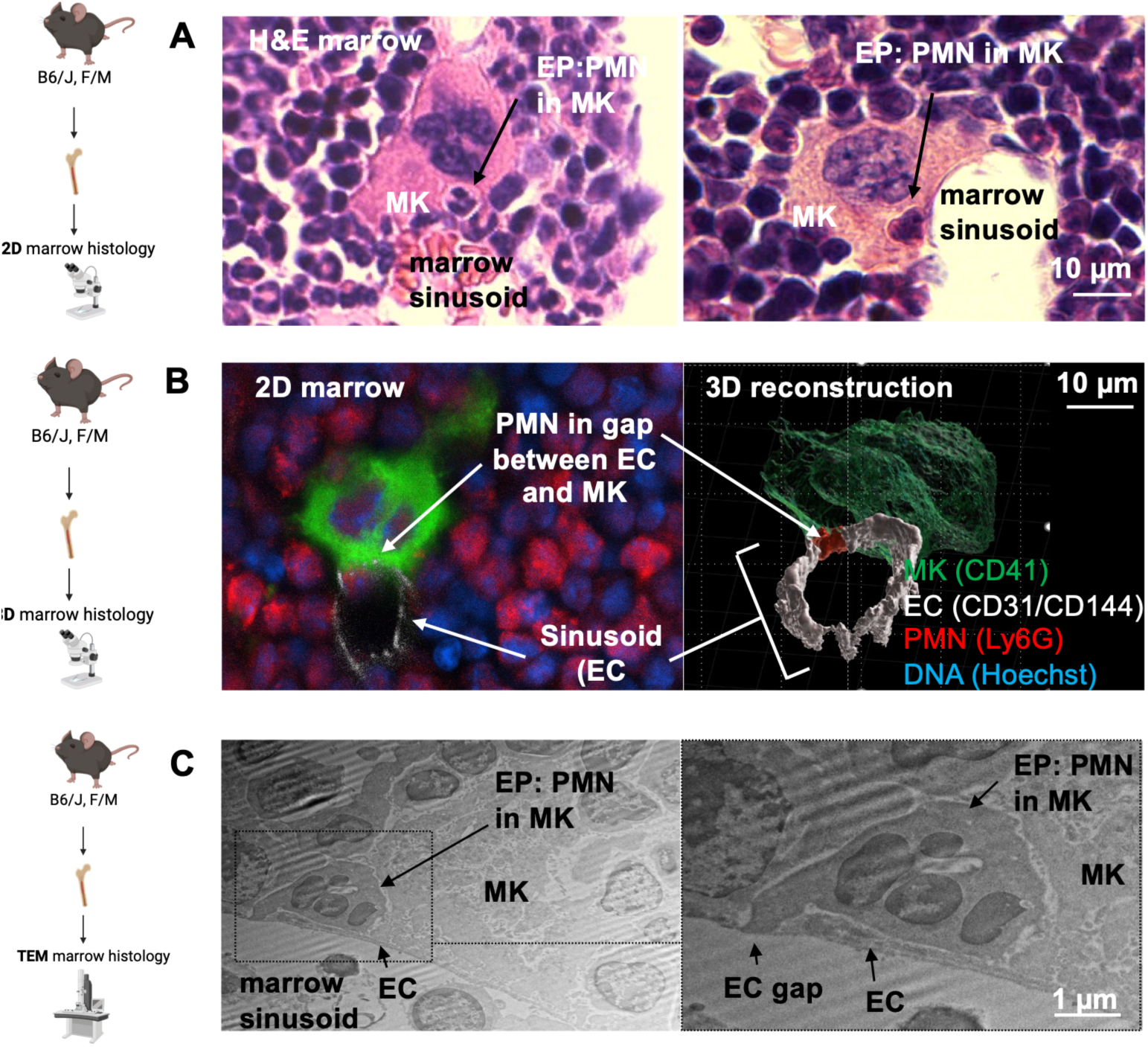
Emperipolesis provides a trans-megakaryocyte route for neutrophil egress from the bone marrow. (A) Light microscopy of H&E-stained sections, representative of 30 specimens. Arrowheads indicate neutrophils at the MK-sinusoid interface. (B) Whole-mount immunofluorescence confocal imaging; left, single 2D Z-stack; right, 3D reconstruction. MKs anti-CD41 (green), neutrophils anti-Ly6G (red), endothelium (EC) anti-CD31/CD144 (gray), DNA Hoechst 33342 (blue); representative of 10 specimens. (C) Transmission electron microscopy (TEM) of OsO_4_-stained sections; representative of 2 specimens. (B,C) Arrowheads indicate neutrophils extending into the sinusoid through an endothelial gap.

**Figure S3.**
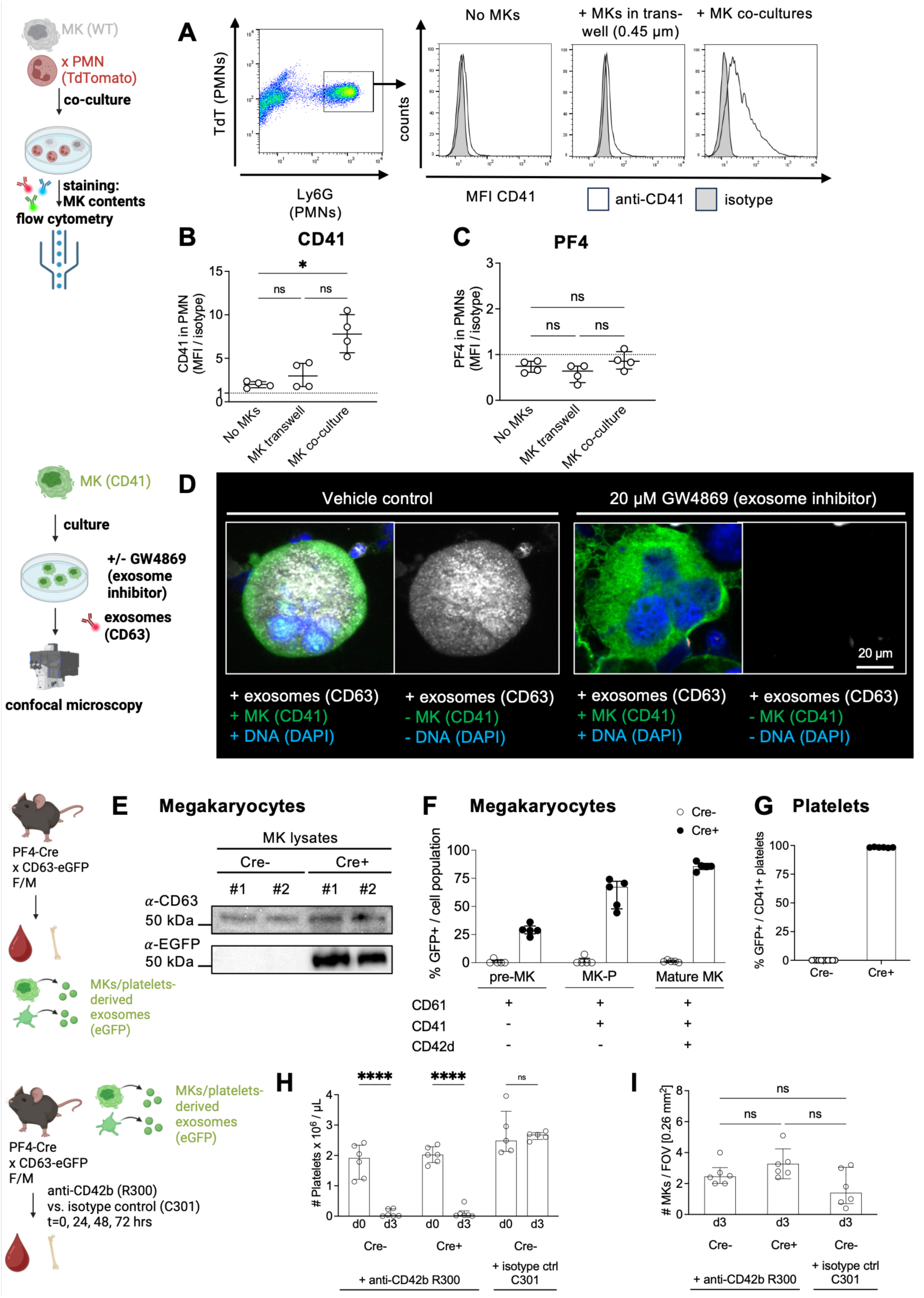
Transiting neutrophils acquire MK exosomes during emperipolesis. (A-C) Flow cytometry of tdTomato⁺ (TdT⁺) neutrophils co-cultured alone, with MKs across a 0.45µm transwell, or with MKs (EP). (A) CD41 in neutrophils (clear) vs isotype (gray). (B,C) CD41 (B) and PF4 (C) MFI in neutrophils, normalized to isotype (n=4). (D) Confocal microscopy of MKs treated with the exosome inhibitor GW4869 (20 µM, 24 hours) vs vehicle. MKs anti-CD41 (green), DNA DAPI (blue), exosomes anti-CD63 (gray). Left, composite; right, CD63 only. (E-G) MKs (E,F) and platelets (G) from PF4-Cre+xCD63emGFP^l/s/l^ mice. (E) Immunoblots of MK lysates, anti-CD63 (upper row), anti-GFP (bottom row), showing a CD63emGFP fusion protein (expected size: ∼50 kDa) in Cre^+^ but not Cre^-^ donors. (F,G) GFP^+^ fraction in CD61^+^CD41^-^CD42d^-^ pre-MKs, CD61^+^CD41^+^CD42d^-^ MK progenitors (MK-P), and CD61^+^CD41^+^CD42d^+^ mature MKs (F) and CD41^+^ platelets (G) (n=5-6/group). (H,I) PF4-Cre^+^xCD63emGFP^l/s/l^ mice and Cre^-^ littermates (n=6/6) were platelet depleted (anti-CD42b, clone R300, 2 mg/kg daily x 72 h vs or isotype, C301). (H) Platelet counts. (I) Marrow MK density. (B,C,F-I) Bars/lines: median; error bars, IQR. Statistics: (B,C,I) multiple Mann–Whitney U tests with FDR adjustment (α < 0.05); (H) multiple Wilcoxon matched-pairs tests with FDR adjustment (α<0.05). Significance: *P ≤ 0.05; ****P ≤ 0.0001.

**Figure S4.**
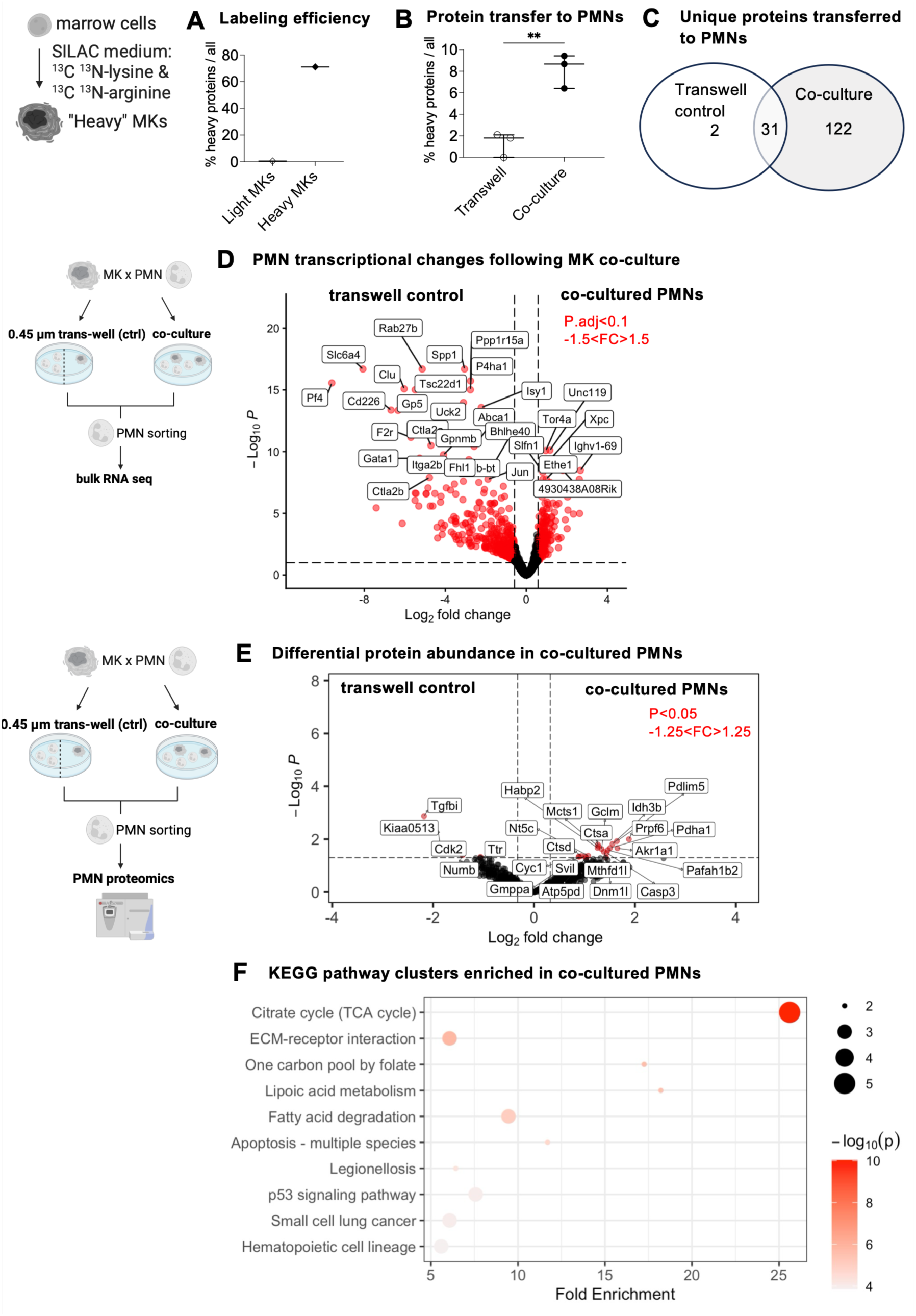
Megakaryocytes transfer proteins related to metabolism, migration, and immunity to neutrophils via exosomes and shape the neutrophil transcriptome and proteome. (A-C) SILAC proteomics to identify contact-dependent protein transfer from MKs to neutrophils. MKs were labeled with ^13^C^15^N-lysine/arginine; unlabeled neutrophils were co-cultured with MKs (EP) or separated by a transwell (control) for 3 hours (n=3 paired samples). (A) MK labeling efficiency (n=1). (B) Abundance of labeled proteins among all proteins in transwell control vs co-culture (n=3). (C) Overlap of unique labeled proteins detected in transwell control vs co-culture (pooled n=3). (D) Bulk transcriptomic analysis of FACS-sorted neutrophils after co-cultures with MKs (EP) vs transwell (control) (3 hours; n=8 paired samples). Volcano plot of differentially abundant transcripts; red, FDR-Padj<0.1 and fold-change>|1.5|. Transwell control neutrophils showed increased abundance of MK-lineage-specific transcripts (e.g., *Pf4* and *Gata1*). (E,F) Proteomics of FACS-sorted neutrophils after co-cultures with MKs (EP) vs transwell (control) (3 hours; n=3 paired samples). (E) Volcano plot of differentially abundant proteins; red, FDR-Padj<0.05 and fold-change>|1.25|. (F) Bubble chart of representative members from the top-10 enriched murine KEGG pathway clusters. (A,B): Line: mean. Statistics: (B) paired t-test. (D,E) Wald test, FDR-adjusted (α<0.1/0.05). (F) hypergeometric test, FDR-adjusted (α<0.1). Significance: \*\**P*≤ 0.01.

**Figure S5.**
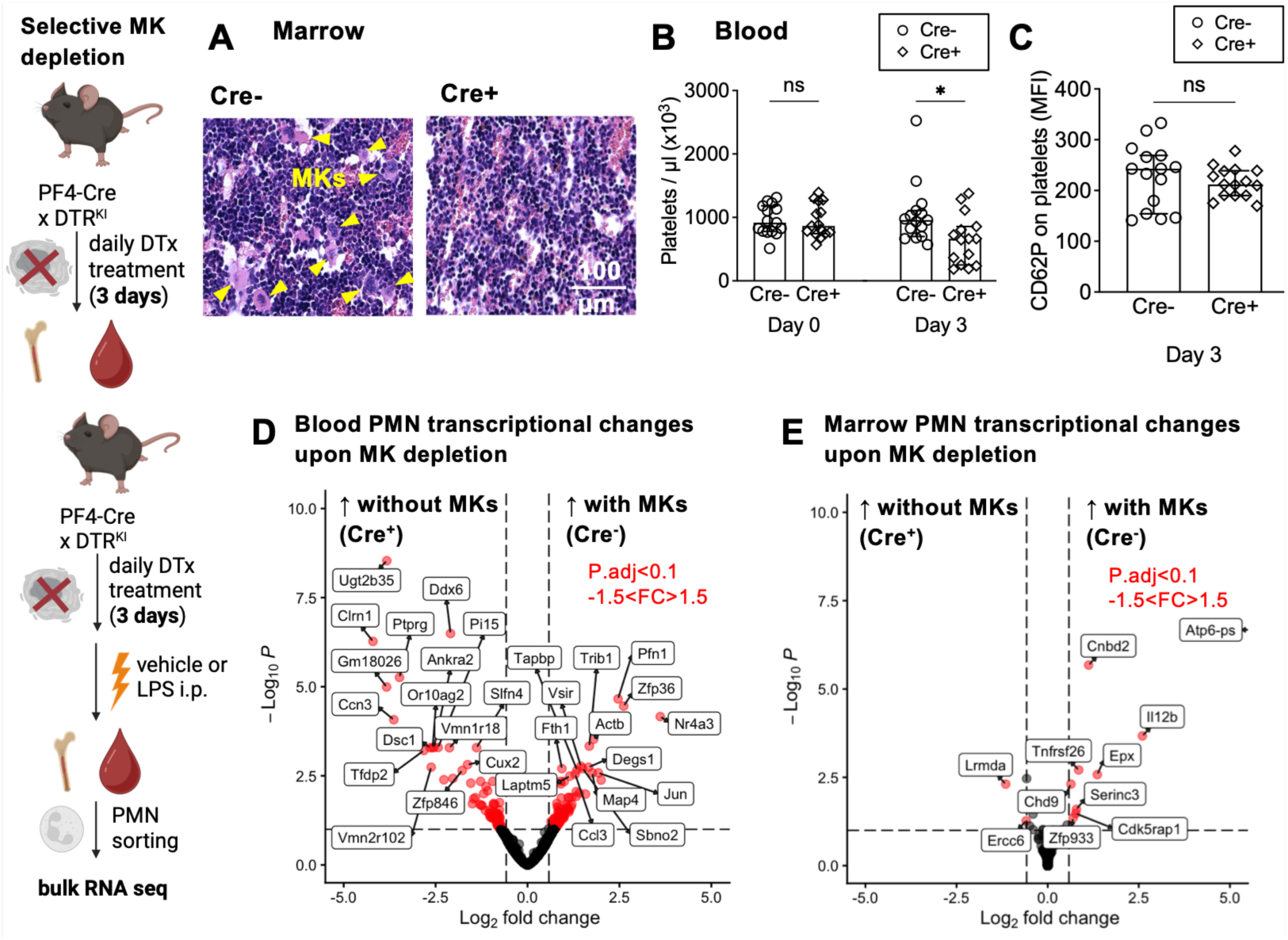
Megakaryocytes shape the neutrophil transcriptome. (A-C) Selective MK depletion in PF4-Cre^+^ x iDTR^fl/fl^ mice vs PF4-Cre^-^ x iDTR^fl/fl^ littermate controls (n=15/15). Mice received diphtheria toxin (DTx; 250 ng i.p. daily x 3 days) to deplete MKs. (A) Representative H&E-stained marrow; MKs present only in Cre^-^ mice (arrowheads). (B,C) Platelet counts (B) and platelet CD62P surface expression (C). (D,E) Blood and marrow neutrophil transcriptome in the presence of MKs. PF4-Cre⁺ × iDTR^fl/fl^ vs Cre⁻ littermates received diphtheria toxin to deplete MKs (250 ng i.p. daily x 3 days), followed by vehicle or LPS treatment (n=3-5/group). FACS-sorted circulating and marrow neutrophils were analyzed for transcriptional changes associated with the presence and absence of MKs across vehicle/LPS treatment. Volcano plot of differentially abundant transcripts in blood (D) and marrow (E) neutrophils; red, FDR-Padj<0.1 and fold-change>|1.5|. (B,D) Bars, median; error bars, IQR. Statistics: (B,C) (multiple) Mann-Whitney U tests, FDR-adjusted (α<0.05); (D,E) Wald test, FDR-adjusted (α<0.1). Significance: *P ≤ 0.05.

**Figure S6.**
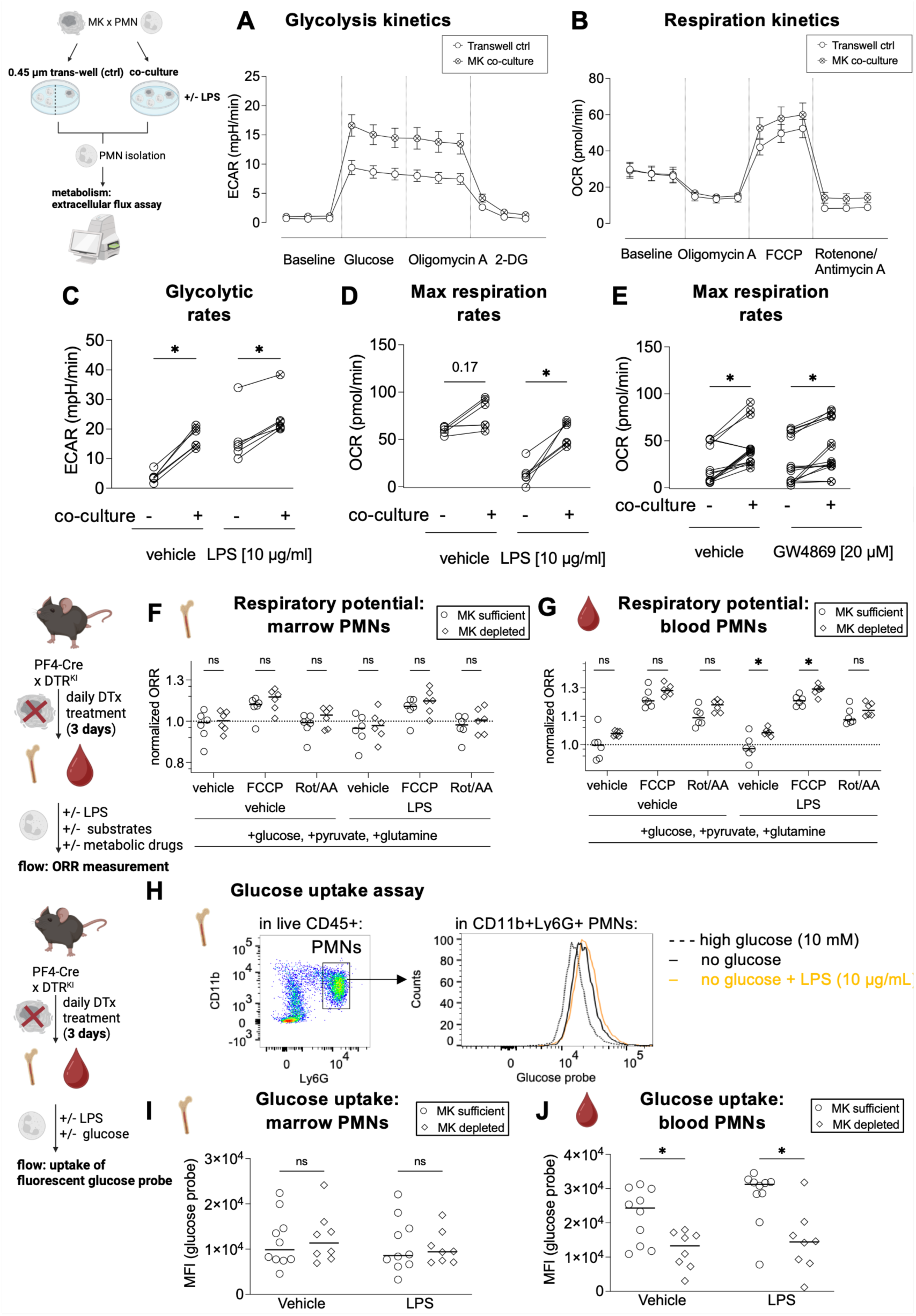
Megakaryocytes enhance neutrophil metabolism. (A-E) Metabolic profiling of neutrophils after MK co-culture (EP, 3 hours) vs transwell (control); ± exosome inhibitor GW4869 (20 µM) or LPS (10 µg/ml). n=21 (A,B), n=6 (C,D), n=15 (E). (A) Glycolytic kinetics (ECAR, extracellular acidification rate), (B) respiratory kinetics (OCR, oxygen consumption rate), (C) glycolytic rate after glucose, (D,E) mitochondrial respiration after FCCP. (F,G) Optical redox ratios (ORRs) in neutrophils from MK-depleted PF4-Cre⁺ × iDTR^fl/fl^ vs Cre⁻ littermates (n=6/group) (250 ng DTx i.p. daily × 3 days); marrow (F) and blood (G). Cells were in the presence of glucose/pyruvate/glutamine, ± LPS (10 µg/ml). ORRs measured via flow cytometry at baseline and after FCCP and Rot/A, normalized to median vehicle. (H-J) Glucose uptake in neutrophils from MK-depleted PF4-Cre^+^ x iDTR^fl/fl^ vs. vs Cre⁻ littermates (n=10/8). (H) Specificity: glucose probe uptake in neutrophils reduced by 10 mM glucose and enhanced by LPS (10 µg/ml). (I,J) Glucose probe uptake in marrow (I) and blood (J) neutrophils after vehicle or LPS. (A,B) Mean ± SEM; (F,G,I,J) Lines: median. Statistics: (C-E) (multiple) Wilcoxon matched-pairs tests, FDR-adjusted (α<0.05); (F,G,I,J) multiple Mann-Whitney U tests, FDR-adjusted (α<0.05). Significance: *P ≤ 0.05. Abbrevations: FCCP: carbonyl cyanide-4 trifluoromethoxy phenylhydrazone, Rot/AA: Rotenone/Antimycin A.

**Figure S7.**
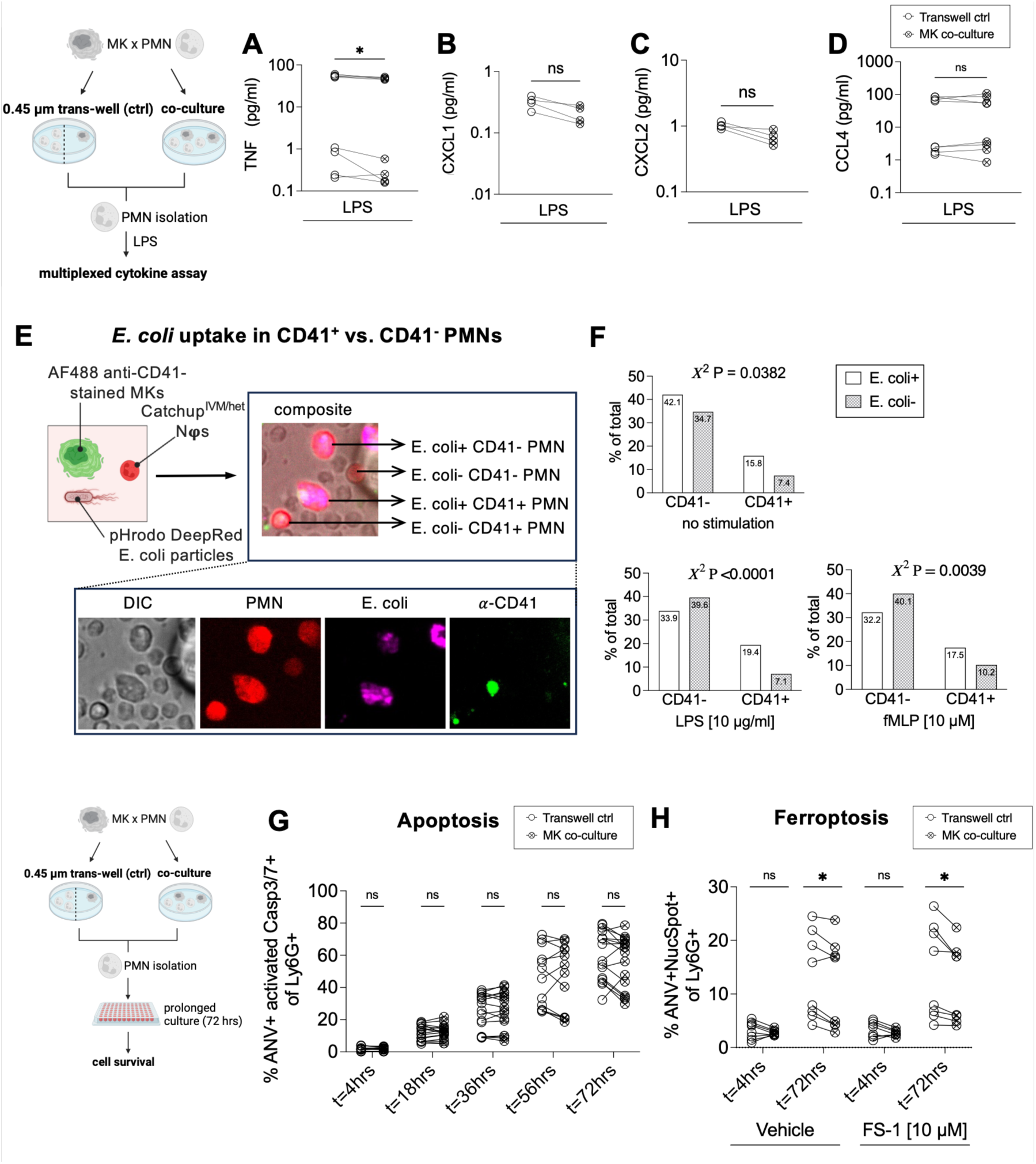
Megakaryocytes enhance neutrophil cytokine production and survival but reduce phagocytosis. (A-C) Neutrophil cytokine release after 3 h co-culture (EP) vs 0.45 µm transwell control, followed by LPS (10 µg/ml) (n=8). (A) TNF, (B) CXCL1, (C) CXCL2, (D) CCL4 concentrations. (E,F) Phagocytosis in CD41⁺ vs CD41⁻ neutrophils after co-culture with MKs. Catchup^IVM/Het^ marrow cells (mTdTomato⁺ neutrophils) were co-cultured 3 h with MKs stained with anti-CD41–AF488, then incubated with fluorescent *E. coli* (Deep Red), ± vehicle, LPS (10 µg/ml), or fMLP (10 µM). (E) Experimental schema and representative images. (F) Quantification by microscopy (n=3; ≥3 FOVs/condition). (G,H) Neutrophil survival after MK co-culture. MKs and marrow cells were co-cultured for 4 hours; control marrow cells were cultured separated by 0.45 µm transwell for 4 h. Neutrophils were then cultured for another 72 h (n=8-12). (G) Apoptosis: % Annexin V⁺ Caspase3/7⁺ among Ly6G⁺. (H) Overall death: % Annexin V⁺ NucSpot⁺ among Ly6G⁺, ± ferroptosis inhibitor Ferrostatin-1 (FS-1, 10 µM). (F) Bars, mean. Statistics: (A-D,G,H) (multiple) Wilcoxon matched-pairs tests, FDR-adjusted (α<0.05); (E) Yates’ continuity-corrected χ² test. Significance: *P ≤ 0.05.

**Figure S8.**
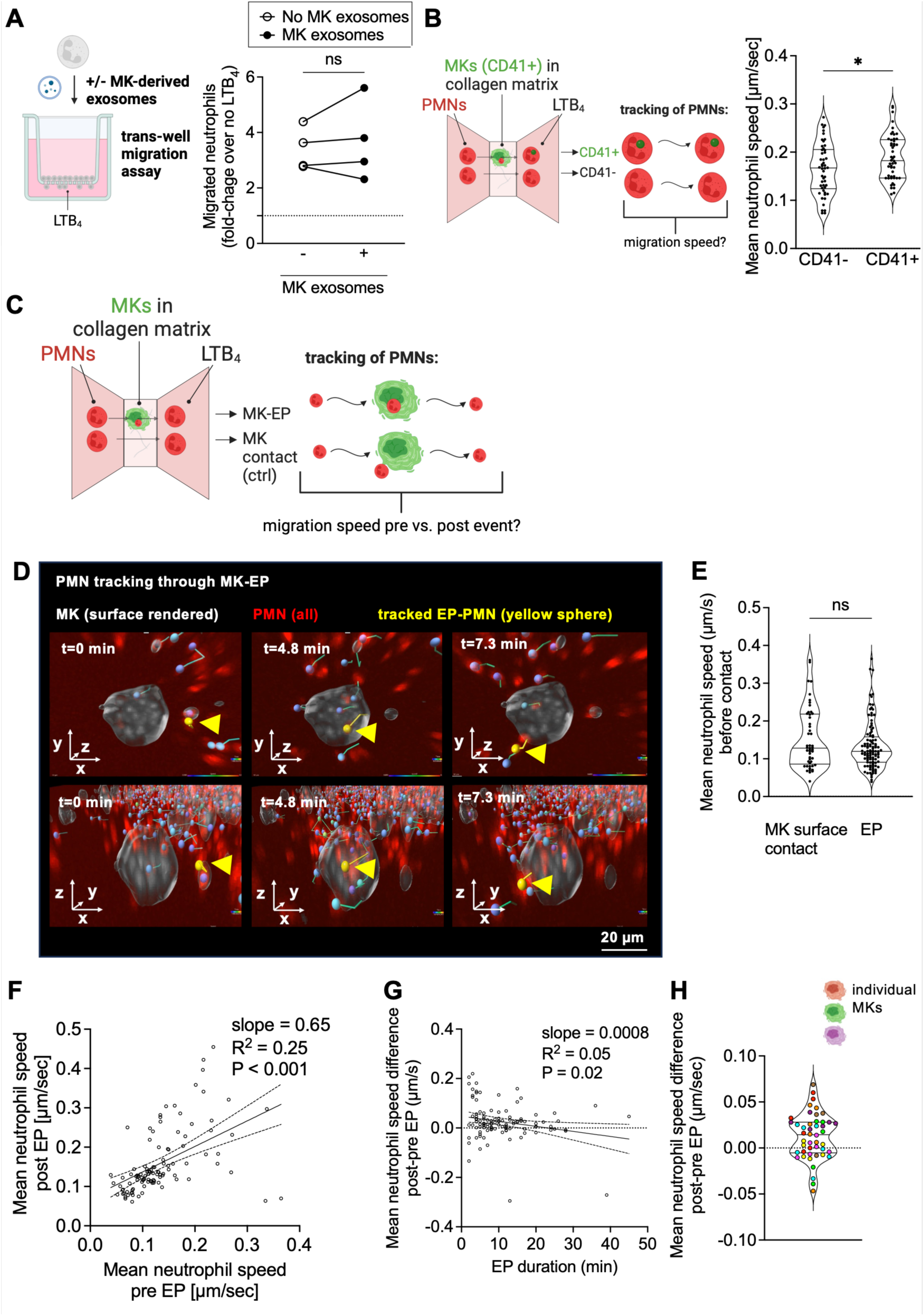
Emperipolesis enhances neutrophil migration. (A) Neutrophil transwell migration (2 hours, 3 µm pores) after culture ± MK-derived exosomes in a LTB_4_ (100 nM) gradient, migration quantified as fold-increase in Ly6G⁺ cells vs no gradient. N=4. (B-H) Trans-MK neutrophil chemotaxis assay. MKs stained with anti-CD41-AF488 (green) were incorporated into collagen. Catchup^IVM/Het^ neutrophils (TdT^+^, red) were imaged migrating through the MK-seeded matrix along a LTB_4_ gradient (100 nM). (B) Speed of CD41⁻ vs CD41⁺ neutrophils (pooled from n=3; 46 tracks/group). (C-H) Speeds of neutrophils pre-/peri-/post-EP and MK-surface contact (controls) (pooled from n=3; 50 control vs 107 EP tracks). (C) Experimental workflow. (D) A neutrophil (yellow sphere, arrowhead) undergoing EP (MK: gray surface, transparent). Upper row: top-down, lower row: side view, confirming PMN passes through the MK. (E) Speeds before MK-surface contact (control) vs before EP. (F) Correlation of pre- vs post-EP mean speeds. (G) Correlation of EP duration with speed changes. (H) Speed changes after EP; each point is a neutrophil undergoing EP, color-coded by MK identity. (B,E,H) Violin plots with median ± IQR lines. (F,G) Linear fit ± 95% CI. Statistics: (A) Wilcoxon matched-pairs test; (B,E) Mann-Whitney U tests; (F,G) linear correlation with t test. Significance: *P ≤ 0.05.

**Figure S9.**
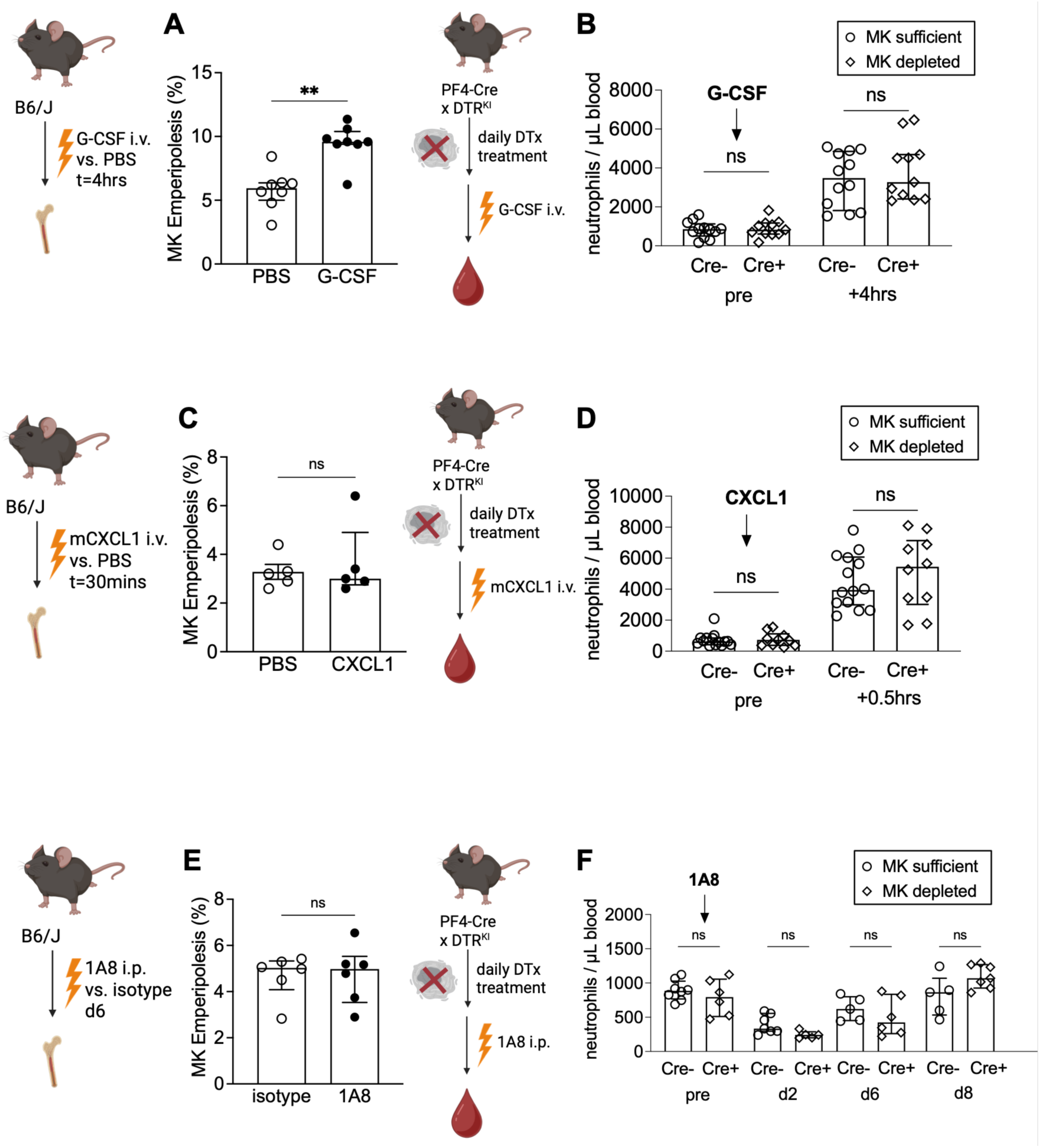
EP does not facilitate neutrophil egress from the bone marrow during steady-state or stress mobilization. (A,C,E) EP quantified in WT mouse H&E-stained marrow sections after (A) G-CSF vs vehicle (2.5 µg i.v.; n=8; 4 h), (C) CXCL1 vs vehicle (300 ng i.v.; n=5; 30 min), or (E) 1A8 vs isotype (2A3) (100 µg i.p.; n=6, 6 days). (B,D,F) Marrow egress in MK-depleted PF4-Cre⁺ × iDTR^fl/fl^ vs Cre⁻ littermates after DTx (250 ng i.p. daily × 6 days). Mice received (B) G-CSF (2.5 µg i.v.; n = 10-12), (D) CXCL1 (300 ng i.v.; n = 10-14), or (F) 1A8 (anti-Ly6G, neutrophil depleting antibody; 100 µg i.p.; n = 5-8). Blood was sampled before and at indicated times; circulating CD45⁺Ly6G⁺CD11b⁺ neutrophils were quantified by flow cytometry. (A-F) Bars, median; error bars, IQR. Statistics: (multiple) Mann-Whitney U tests, FDR-adjusted (α<0.05). Significance: **P ≤ 0.01.

**Figure S10.**
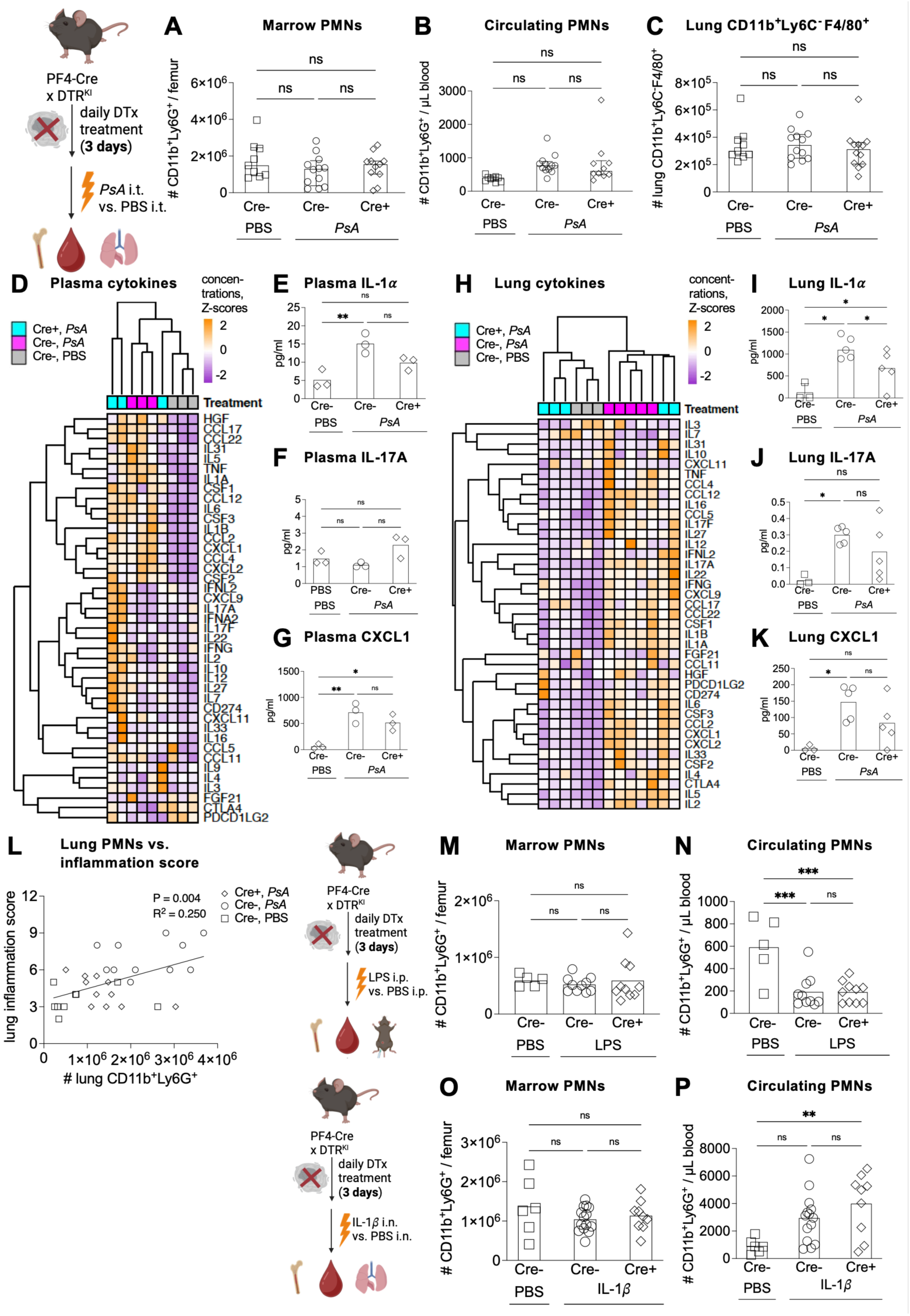
Emperipolesis enhances neutrophil entry into inflamed tissues. (A-L) *Pseudomonas aeruginosa* (PsA) pneumonia in MK-depleted PF4-Cre⁺×iDTR^fl/fl^ vs Cre⁻ littermates (250 ng DTx i.p. daily × 3 days) was induced by i.t. inoculation of PsA (strain PA103, 3×10^5^ CFU, 4 hours) vs vehicle (PBS, control); where indicated, neutrophils were depleted with 1A8 (24 hours prior). (A,B) CD45⁺Ly6G⁺CD11b⁺ neutrophils in femur (A) and blood (B). (C) CD45⁺CD11b⁺F4/80⁺Ly6C^-^ lung myeloid cells. (A-C) n=9/12/11. (D-K) Cytokine/chemokine concentrations in plasma (D-G; n=3/3/3) and lung homogenates (H-K; n=3/5/5). (D,H) Heatmaps of Z-score-normalized concentrations (Ward D2 hierarchical clustering). (E,I) IL-1*α*, (F,J) IL-17A, (G,K) CXCL1. (L) Correlation of lung CD11b^+^Ly6G^+^ neutrophils with histologic lung inflammation score (n=9/12/11). (M,N) LPS peritonitis (2 mg/kg i.p.; 2 hours) in MK-depleted PF4-Cre⁺×iDTR^fl/fl^ vs Cre⁻ littermates (DTX as above). CD45⁺Ly6G⁺CD11b⁺ neutrophils in marrow (M) and blood (N) (n=5/9/9). (O,P) IL-1β pneumonitis (1 µg/kg i.n.; 4 hours) in MK-depleted PF4-Cre⁺×iDTR^fl/fl^ vs Cre⁻ littermates (DTX as above). CD45⁺Ly6G⁺CD11b⁺ neutrophils in marrow (O) and blood (P) (n=6/13/9). (A-C,E-G,I-K,M-P) Bars, median; error bars, IQR. (L) Line, linear correlation. Statistics: (A-C,E-G,I-K,M-P) one-way ANOVA with Holm-Šídák adjustment (α<0.05); (L) linear correlation with t test. Significance: *P ≤ 0.05; **P ≤ 0.01, ***P ≤ 0.001.

**Figure S11.**
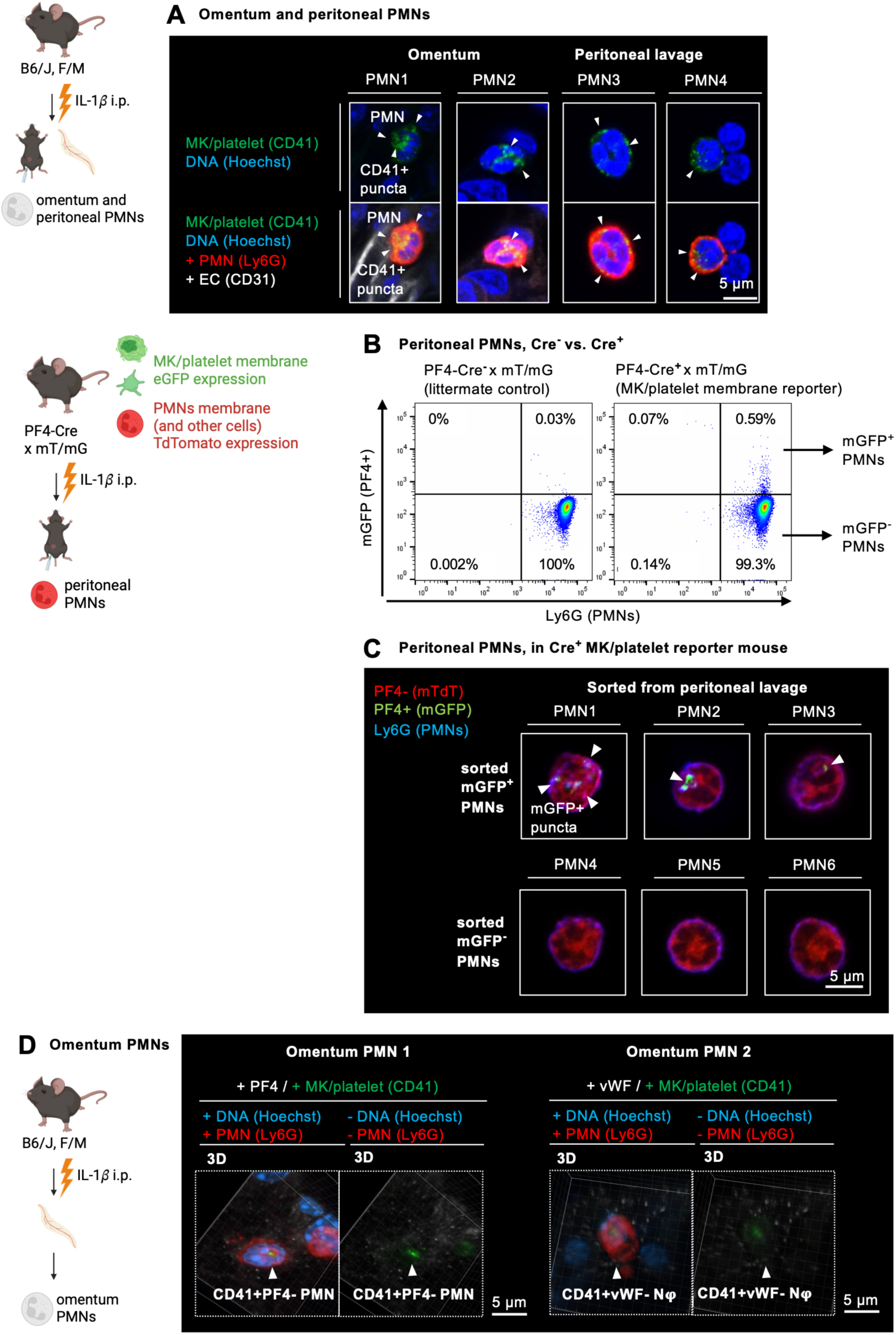
Emperipolesis enhances neutrophil entry into inflamed tissues. (A) Peritonitis was induced in WT mice (IL-1β, 1 µg/kg i.p.); omentum (left) and peritoneal cells (right) were harvested at 2 h for confocal imaging. Anti-CD41 (MK/platelet material, green), anti-Ly6G (neutrophils, red), anti-CD31 (endothelium, gray), DAPI (DNA, blue). Top, CD41 and DAPI; bottom, composite. Arrowheads, internalized CD41⁺ puncta. Representative of n=3. (B,C) Peritonitis was induced in PF4-Cre⁺ × mT/mG membrane reporter (PF4⁺ membranes mGFP⁺, green, ubiquitous membranes mTdTomato⁺, red) with IL-1β as above; followed by peritoneal lavage at 2 h. (B) Flow cytometry showing GFP⁺ MK/platelet membrane material in Ly6G⁺ neutrophils from Cre⁺ but not Cre⁻ littermates. (C) Sorted mGFP⁺ vs mGFP⁻ neutrophils stained with anti-Ly6G (neutrophils, blue); arrowheads, internalized CD41⁺ material. Representative of n=3. (D) Peritonitis was induced in WT with IL-1β as above; omentum harvested at 2 h was stained with anti-CD41 (MK/platelet material, green), anti-Ly6G (neutrophils, red), DAPI (DNA, blue) and α-granule proteins PF4 or vWF (gray, where indicated). Left, composite; right, CD41 with PF4/vWF only. Arrowheads, neutrophils internalizing CD41⁺ material; punctate CD41⁺PF4⁺ and CD41⁺vWF⁺ deposits identify platelets (validate staining). Representative of n=3.

## EXTENDED DATA MOVIES

**Video S1. MK neutrophil emperipolesis (EP) occurring in mouse marrow *in vivo***. Two-photon intravital microscopy of calvarial marrow in Catchup^IVM^ x vWF-EGFP mice; neutrophils tdTomato (red; pseudo-colored magenta when internalized), MKs EGFP (green, surface-rendered and masked), vasculature Evans Blue (gray). (Supplementary to Figure 1C).

**Video S2. A MK harboring fast and slow neutrophil EP simultaneously *in vivo*.** Two-photon intravital microscopy (2P-IVM) of calvarial marrow in Catchup^IVM^ × vWF-EGFP mice; neutrophils tdTomato (red; pseudo-colored magenta when internalized), MKs EGFP (green), vasculature Evans Blue (gray). (Supplementary to Figure S1J).

**Video S3. A neutrophil at the interface between MKs and marrow sinusoids in direct contact with the circulation.** 3D reconstruction of whole-mount immunofluorescence confocal imaging of mouse bone marrow histology. MKs anti-CD41 (green), neutrophils anti-Ly6G (red), endothelium (EC) anti-CD31/CD144 (gray), DNA Hoechst 33342 (blue). MKs, neutrophils and EC were surface-rendered and masked. (Supplementary to Figure S2B).

**Video S4. A neutrophil egressing from the bone marrow into the circulation via a trans-MK route *in vivo*.** Two-photon intravital microscopy of calvarial marrow in Catchup^IVM^ x vWF-EGFP mice; neutrophils tdTomato (red; pseudo-colored magenta when internalized), MKs EGFP (green), vasculature Evans Blue (gray). The neutrophil undergoing EP rapidly disappears from the field of view because it is swept away by the circulation. (Supplementary to Figure 1H).

**Video S5. Neutrophils migrating in inflamed omentum containing internalized CD41^+^ punctate material.** Peritonitis was induced in WT mice (IL-1β, 1 µg/kg i.p.); omentum was harvested at 2 h for confocal imaging. Anti-CD41 (MK/platelet material, green), anti-Ly6G (neutrophils, red), anti-CD31 (endothelium, gray), DAPI (DNA, blue). Series of 3D reconstructions of 3 neutrophils harboring internalized green CD41^+^ punctate material. (Supplementary to Figure S12A).

## EXTENDED DATA TABLES

**Supplementary Tables 1-3: SILAC proteomics to identify proteins transferred from megakaryocytes to neutrophils in a contact-dependent manner.** The peptides of murine marrow-derived megakaryocytes (MKs) were labeled with ^13^C ^15^N-lysine and ^13^C ^15^N-arginine. Following validation of labeling efficiency via mass spectrometry, labeled MKs were co-cultured with unlabeled murine marrow neutrophils for 3 hours (emperipolesis), or co-cultured with unlabeled neutrophils separated by 0.45 µm trans-well inserts (control). Neutrophils were FACS-sorted as single, live, CD45+CD11b+Ly6G+ cells and lysed in RIPA buffer. Proteins in the neutrophil lysates were separated via SDS-PAGE and the presence of labeled vs. unlabeled proteins were identified by mass spectrometry (S1). The identified labeled proteins were analyzed using the publicly accessible Protein Analysis THrough Evolutionary Relationships (PANTHER) resource (https://www.pantherdb.org/)^52^, assessing Gene Ontology Cellular Component Ontology (S2) and PANTHER Pathway ontology (S3). N=3.

**Supplementary Table 1.**
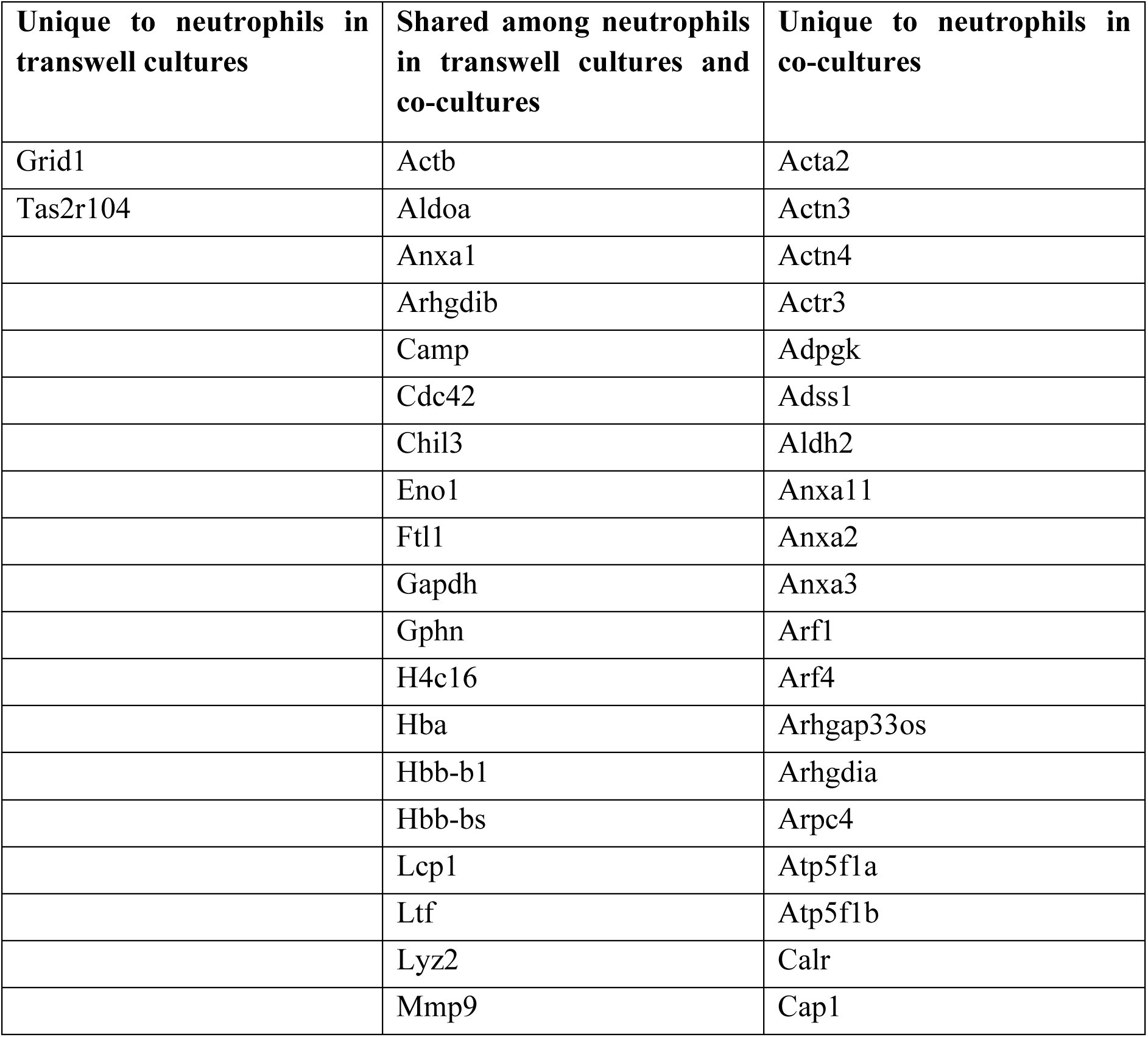
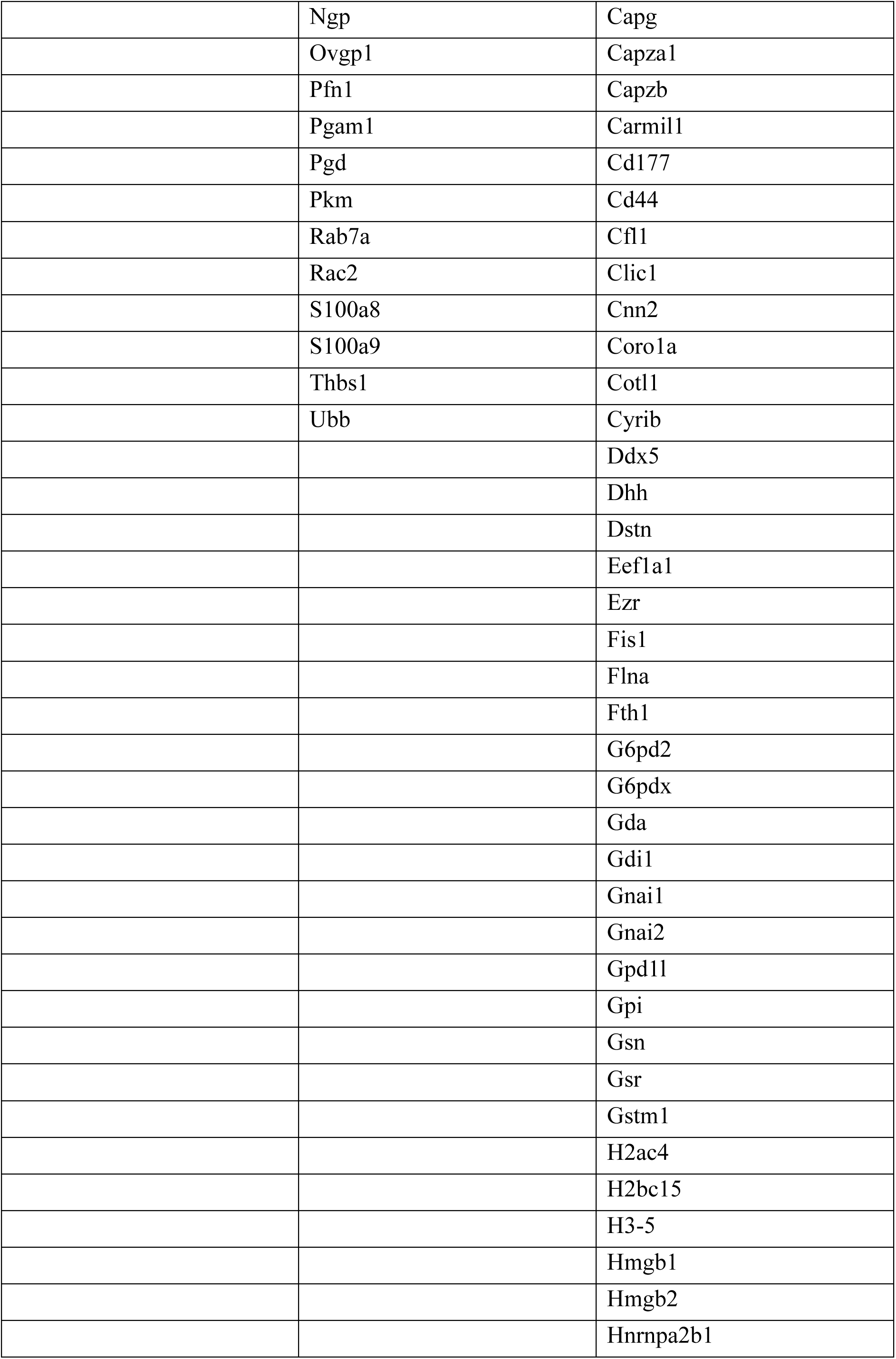
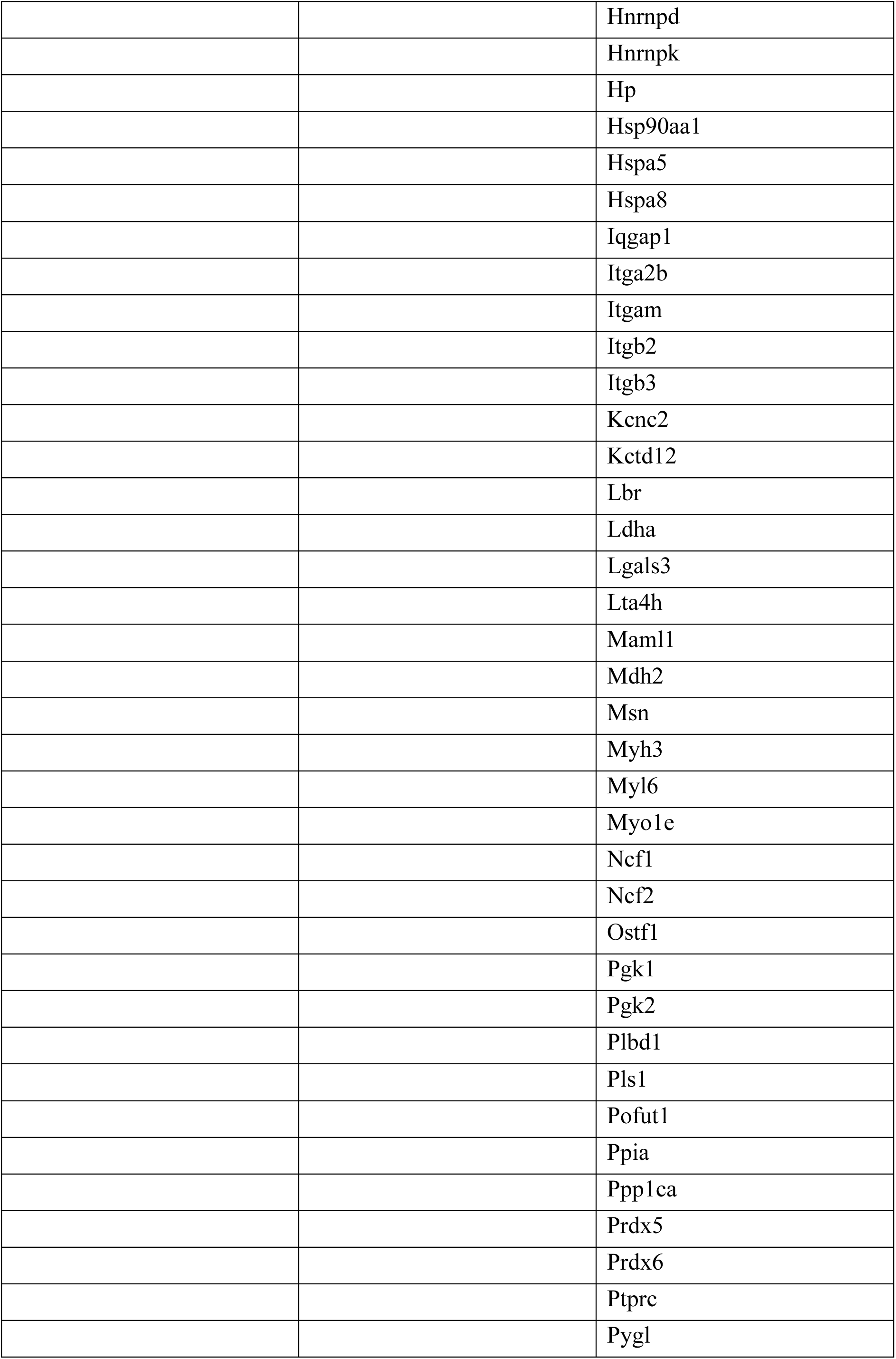
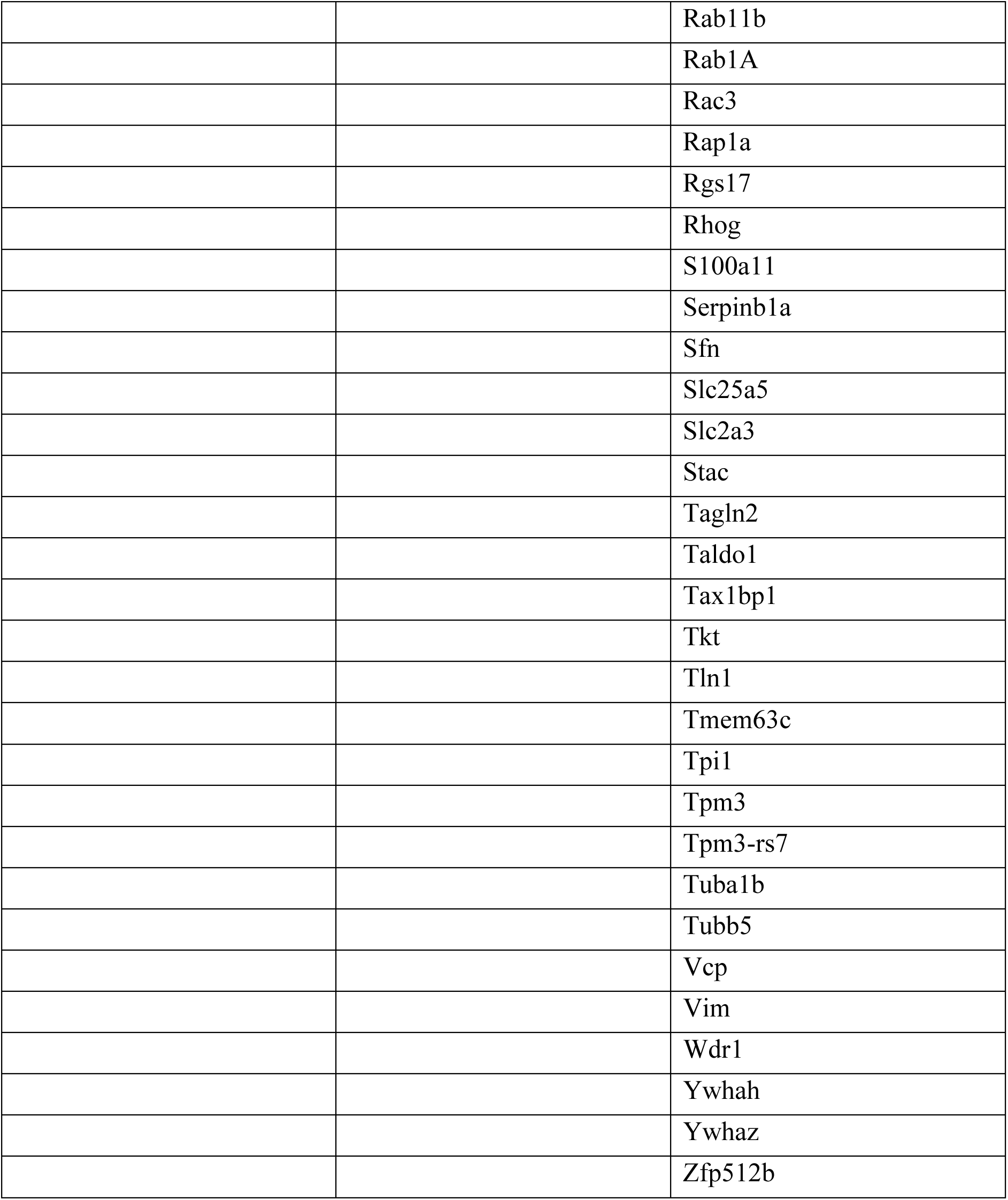
Unique labeled proteins (as Gene symbols) identified in neutrophils sorted from transwell culture control conditions only, co-culture conditions only, and both conditions.

**Supplementary Table 2.** Overrepresentation analysis for GO Cellular Component Ontology annotation terms of labeled proteins only identified in neutrophils sorted from co-culture conditions. Analysis was conducted using the publicly accessible Protein Analysis THrough Evolutionary Relationships (PANTHER) overrepresentation test tool (released 2024-08-07)^52^, assessing the GO cellular component complete database (version: 21836) from the GO ontology database (version: DOI: 10.5281/zenodo.15066566, released 2025-03-16) and as a reference list, the genome of *Mus musculus* provided in the PANTHER database (version: 19.0). P-values: Fisher’s exact test, FDR-adjusted (α<0.05). Results with fold-enrichment ≥1.5 only.

| GO annotation term, Cellular Component Ontology | Fold-enrichment | Adj. P value |
| --- | --- | --- |
| basal ectoplasmic specialization (GO:0061832) | > 100 | 3.03E-03 |
| integrin alphaIIb-beta3 complex (GO:0070442) | > 100 | 2.98E-03 |
| alphav-beta3 integrin-HMGB1 complex (GO:0035868) | > 100 | 2.94E-03 |
| integrin alphaM-beta2 complex (GO:0034688) | > 100 | 2.89E-03 |
| apical ectoplasmic specialization (GO:0061831) | 85.97 | 5.53E-03 |
| macropinosome (GO:0044354) | 42.98 | 1.99E-02 |
| pinosome (GO:0044352) | 42.98 | 1.97E-02 |
| F-actin capping protein complex (GO:0008290) | 38.21 | 2.39E-02 |
| Arp2/3 protein complex (GO:0005885) | 38.21 | 2.37E-02 |
| WASH complex (GO:0071203) | 26.45 | 4.54E-02 |
| NADPH oxidase complex (GO:0043020) | 26.45 | 4.50E-02 |
| cortical actin cytoskeleton (GO:0030864) | 24.3 | 2.00E-12 |
| cortical cytoskeleton (GO:0030863) | 23.96 | 3.37E-16 |
| lamellipodium membrane (GO:0031258) | 22.43 | 7.88E-03 |
| podosome (GO:0002102) | 21.05 | 1.69E-05 |
| integrin complex (GO:0008305) | 20.84 | 1.24E-03 |
| myelin sheath (GO:0043209) | 17.11 | 3.71E-17 |
| microvillus membrane (GO:0031528) | 15.17 | 2.07E-02 |
| melanosome (GO:0042470) | 14.33 | 8.85E-07 |
| pigment granule (GO:0048770) | 14.33 | 8.57E-07 |
| immunological synapse (GO:0001772) | 13.75 | 5.57E-03 |
| actin filament (GO:0005884) | 13.43 | 2.80E-07 |
| brush border (GO:0005903) | 12.61 | 1.04E-07 |
| actin filament bundle<br>(GO:0032432) | 12.5 | 1.18E-05 |
| protein complex involved in<br>cell adhesion (GO:0098636) | 11.66 | 9.42E-03 |
| phagocytic vesicle<br>(GO:0045335) | 11.63 | 4.83E-06 |
| ruffle (GO:0001726) | 10.58 | 1.28E-07 |
| cell-substrate junction<br>(GO:0030055) | 10.4 | 4.01E-08 |
| focal adhesion (GO:0005925) | 10.26 | 1.74E-07 |
| ruffle membrane (GO:0032587) | 10.11 | 9.73E-04 |
| actin cytoskeleton<br>(GO:0015629) | 10.06 | 5.72E-20 |
| cell cortex (GO:0005938) | 10.06 | 6.69E-13 |
| cluster of actin-based cell<br>projections (GO:0098862) | 9.21 | 5.12E-07 |
| cytoplasmic side of plasma<br>membrane (GO:0009898) | 9.21 | 2.58E-05 |
| extracellular exosome<br>(GO:0070062) | 8.96 | 6.54E-03 |
| lamellipodium (GO:0030027) | 8.68 | 3.61E-06 |
| contractile actin filament<br>bundle (GO:0097517) | 8.6 | 7.70E-03 |
| stress fiber (GO:0001725) | 8.6 | 7.60E-03 |
| membrane raft (GO:0045121) | 8.52 | 2.15E-09 |
| membrane microdomain<br>(GO:0098857) | 8.47 | 2.21E-09 |
| endocytic vesicle<br>(GO:0030139) | 8.05 | 6.57E-06 |
| actomyosin (GO:0042641) | 7.89 | 1.05E-02 |
| cell leading edge (GO:0031252) | 7.86 | 1.41E-11 |
| filopodium (GO:0030175) | 7.29 | 1.47E-02 |
| extracellular vesicle<br>(GO:1903561) | 7.29 | 1.46E-02 |
| cytoplasmic side of membrane<br>(GO:0098562) | 7.26 | 1.63E-04 |
| microvillus (GO:0005902) | 7.1 | 1.61E-02 |
| extracellular organelle<br>(GO:0043230) | 6.77 | 1.96E-02 |
| extracellular membrane-<br>bounded organelle<br>(GO:0065010) | 6.77 | 1.94E-02 |
| leading edge membrane<br>(GO:0031256) | 6.71 | 9.16E-04 |
| midbody (GO:0030496) | 6.47 | 3.91E-04 |
| plasma membrane raft<br>(GO:0044853) | 5.93 | 3.06E-02 |
| actin-based cell projection<br>(GO:0098858) | 5.67 | 1.03E-03 |
| microbody (GO:0042579) | 5.66 | 3.70E-02 |
| peroxisome (GO:0005777) | 5.66 | 3.66E-02 |
| site of polarized growth<br>(GO:0030427) | 4.91 | 1.37E-02 |
| anchoring junction<br>(GO:0070161) | 4.87 | 6.44E-08 |
| cell-cell junction (GO:0005911) | 4.79 | 1.11E-05 |
| cell projection membrane<br>(GO:0031253) | 4.76 | 7.24E-04 |
| glutamatergic synapse<br>(GO:0098978) | 4.61 | 6.93E-08 |
| collagen-containing<br>extracellular matrix<br>(GO:0062023) | 4.55 | 4.81E-03 |
| perikaryon (GO:0043204) | 4.54 | 3.96E-02 |
| contractile muscle fiber<br>(GO:0043292) | 4.52 | 1.99E-02 |
| perinuclear region of cytoplasm<br>(GO:0048471) | 4.37 | 3.68E-07 |
| growth cone (GO:0030426) | 4.35 | 4.73E-02 |
| basolateral plasma membrane<br>(GO:0016323) | 3.9 | 4.08E-02 |
| cell body (GO:0044297) | 3.8 | 1.24E-05 |
| supramolecular fiber<br>(GO:0099512) | 3.7 | 5.38E-06 |
| supramolecular polymer<br>(GO:0099081) | 3.68 | 5.90E-06 |
| distal axon (GO:0150034) | 3.64 | 1.93E-02 |
| postsynapse (GO:0098794) | 3.63 | 6.79E-06 |
| polymeric cytoskeletal fiber<br>(GO:0099513) | 3.62 | 3.10E-04 |
| side of membrane<br>(GO:0098552) | 3.6 | 9.97E-05 |
| plasma membrane protein<br>complex (GO:0098797) | 3.36 | 6.53E-03 |
| cell projection (GO:0042995) | 3.33 | 1.57E-14 |
| supramolecular complex<br>(GO:0099080) | 3.3 | 1.81E-06 |
| extracellular matrix<br>(GO:0031012) | 3.3 | 2.01E-02 |
| external encapsulating structure<br>(GO:0030312) | 3.29 | 2.03E-02 |
| cytoskeleton (GO:0005856) | 3.25 | 1.65E-11 |
| neuronal cell body<br>(GO:0043025) | 3.17 | 2.38E-03 |
| external side of plasma<br>membrane (GO:0009897) | 3.16 | 2.61E-02 |
| synapse (GO:0045202) | 3.16 | 4.79E-08 |
| plasma membrane bounded cell<br>projection (GO:0120025) | 3.08 | 1.43E-11 |
| cell junction (GO:0030054) | 3.04 | 2.97E-10 |
| cell surface (GO:0009986) | 2.87 | 1.04E-03 |
| neuron projection<br>(GO:0043005) | 2.72 | 9.47E-05 |
| cytosol (GO:0005829) | 2.65 | 1.36E-13 |
| dendrite (GO:0030425) | 2.61 | 2.85E-02 |
| dendritic tree (GO:0097447) | 2.61 | 2.88E-02 |
| axon (GO:0030424) | 2.59 | 3.04E-02 |
| presynapse (GO:0098793) | 2.58 | 3.14E-02 |
| somatodendritic compartment<br>(GO:0036477) | 2.57 | 5.72E-03 |
| vesicle (GO:0031982) | 2.55 | 6.19E-06 |
| membrane protein complex<br>(GO:0098796) | 2.46 | 8.98E-03 |
| cytoplasmic vesicle<br>(GO:0031410) | 2.41 | 1.52E-04 |
| intracellular vesicle<br>(GO:0097708) | 2.4 | 1.55E-04 |
| plasma membrane region<br>(GO:0098590) | 2.33 | 7.91E-03 |
| mitochondrion (GO:0005739) | 2.2 | 3.92E-03 |
| intracellular membraneless<br>organelle (GO:0043232) | 2.16 | 9.04E-09 |
| membraneless organelle<br>(GO:0043228) | 2.16 | 1.45E-08 |
| extracellular region<br>(GO:0005576) | 1.96 | 9.58E-03 |
| cell periphery (GO:0071944) | 1.84 | 1.78E-07 |
| plasma membrane<br>(GO:0005886) | 1.77 | 1.19E-05 |
| protein-containing complex<br>(GO:0032991) | 1.77 | 2.45E-05 |
| cytoplasm (GO:0005737) | 1.58 | 1.37E-11 |

**Supplementary Table 3.** Overrepresentation analysis for PANTHER pathway terms of labeled proteins only identified in neutrophils sorted from co-culture conditions. Analysis was conducted using the PANTHER overrepresentation test tool (released 2024-08-07)^52^, assessing the PANTHER pathway database (version: 19.0, released 2024-06-20) and as a reference list, the genome of *Mus musculus* provided in the PANTHER database. P-values: Fisher’s exact test, FDR-adjusted (α<0.05).

| PANTHER pathway | Fold-enrichment | Adj. P value |
| --- | --- | --- |
| Pentose phosphate pathway (P02762) | 57.31 | 5.07E-04 |
| ATP synthesis (P02721) | 42.98 | 1.48E-02 |
| Glycolysis (P00024) | 22.43 | 6.29E-03 |
| Nicotine pharmacodynamics pathway (P06587) | 15.17 | 1.48E-02 |
| Cytoskeletal regulation by Rho GTPase (P00016) | 13.4 | 2.34E-04 |
| Dopamine receptor mediated signaling pathway (P05912) | 12.28 | 5.67E-03 |
| Integrin signalling pathway (P00034) | 10.92 | 9.06E-08 |
| PI3 kinase pathway (P00048) | 9.73 | 4.19E-02 |
| Parkinson disease (P00049) | 9.34 | 5.41E-03 |
| Huntington disease (P00029) | 7.07 | 4.93E-03 |
| Inflammation mediated by chemokine and cytokine signaling pathway (P00031) | 6.64 | 1.54E-04 |
| EGF receptor signaling pathway (P00018) | 6.46 | 1.46E-02 |
| Heterotrimeric G-protein signaling pathway-Gi alpha and Gs alpha mediated pathway (P00026) | 5.44 | 2.89E-02 |

**Supplementary Tables 4-8: Transcriptomic analysis**

**Transcriptomic analysis of neutrophils following co-culture with MKs.** Murine marrow neutrophils were co-cultured with MKs for 3 hours (emperipolesis) and control neutrophils were co-cultured with MKs separated by 0.45 µm transwell inserts (control). Neutrophils were FACS-sorted as single, live, CD45+CD11b+Ly6G+CD41- cells, lysed, and isolated RNA was sent for bulk transcriptomic analysis. Differential gene expression analysis was performed using DSeq2 (v3.20) (Wald test, with mouse ID included as a blocking factor for paired design, FDR: Padj<0.10) (**Tables S4,5**).^107^ Gene Ontology (GO) term enrichment was done using clusterProfiler (v4.14.3)^108^ (FDR: Padj<0.10), with annotation based on org.Mm.eg.db (v3.20.0) (**Tables S6,7**). N=8 paired samples (from 4F/4M mice).

**Supplementary Table 4.**
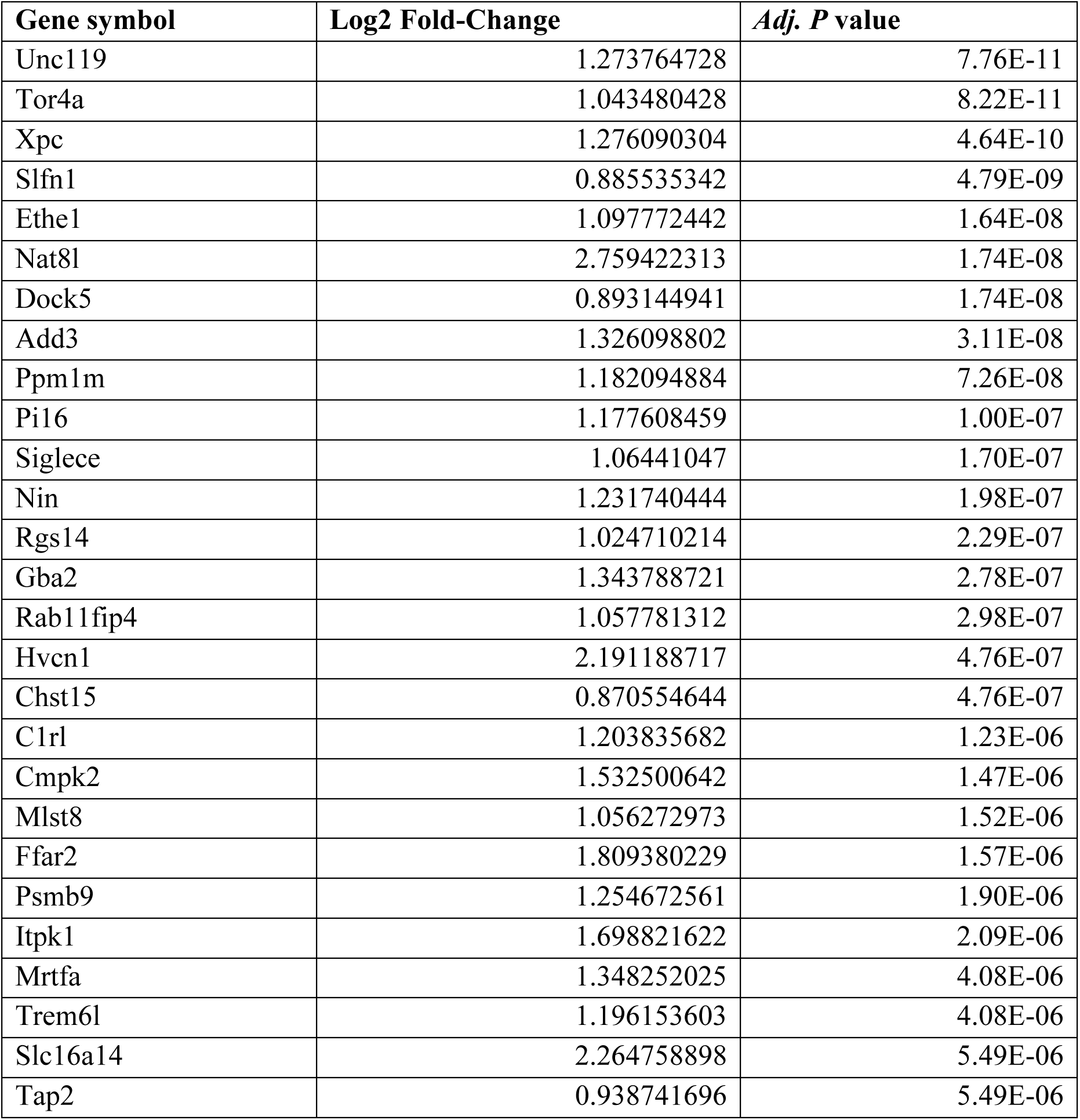
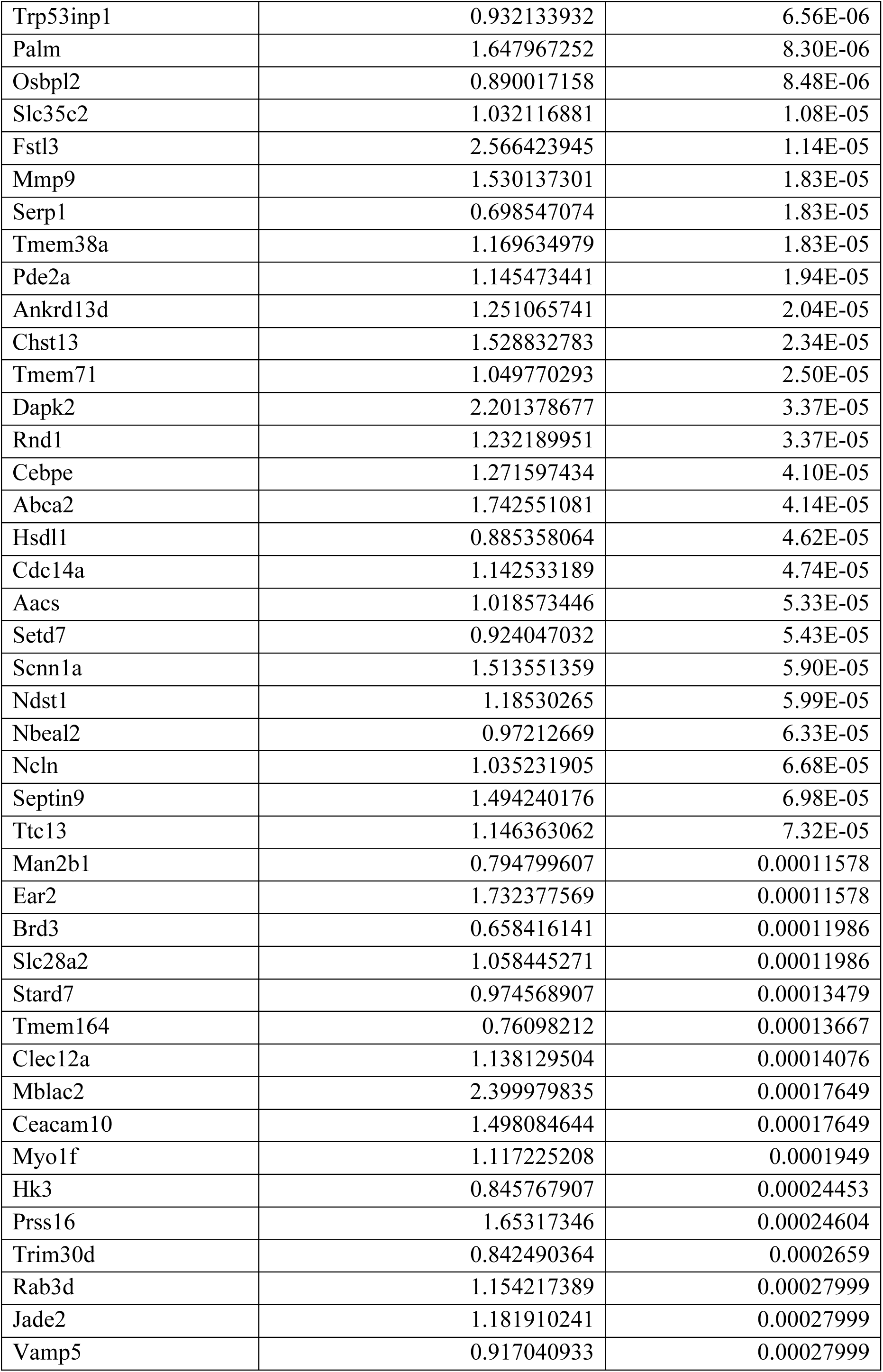
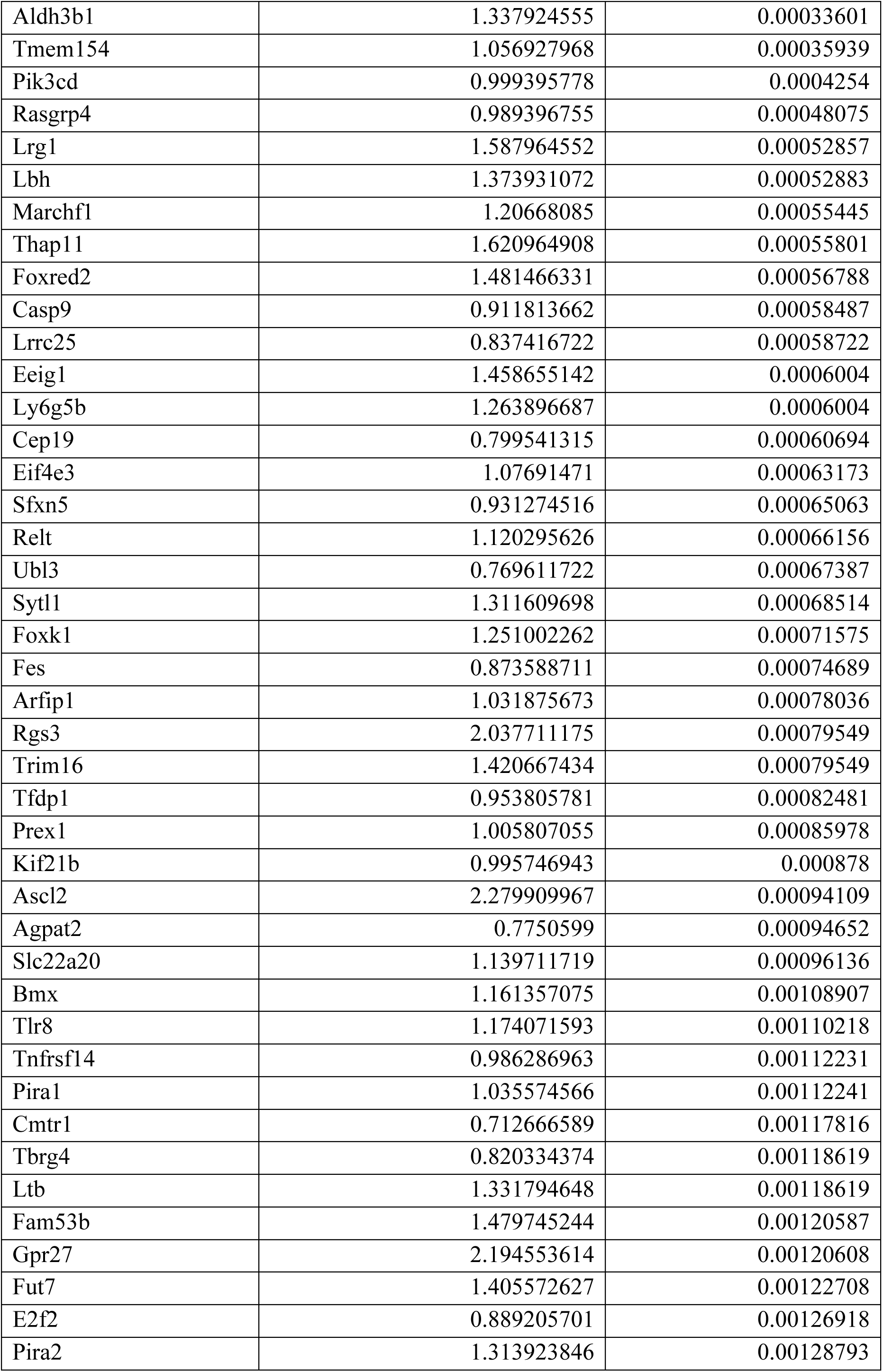
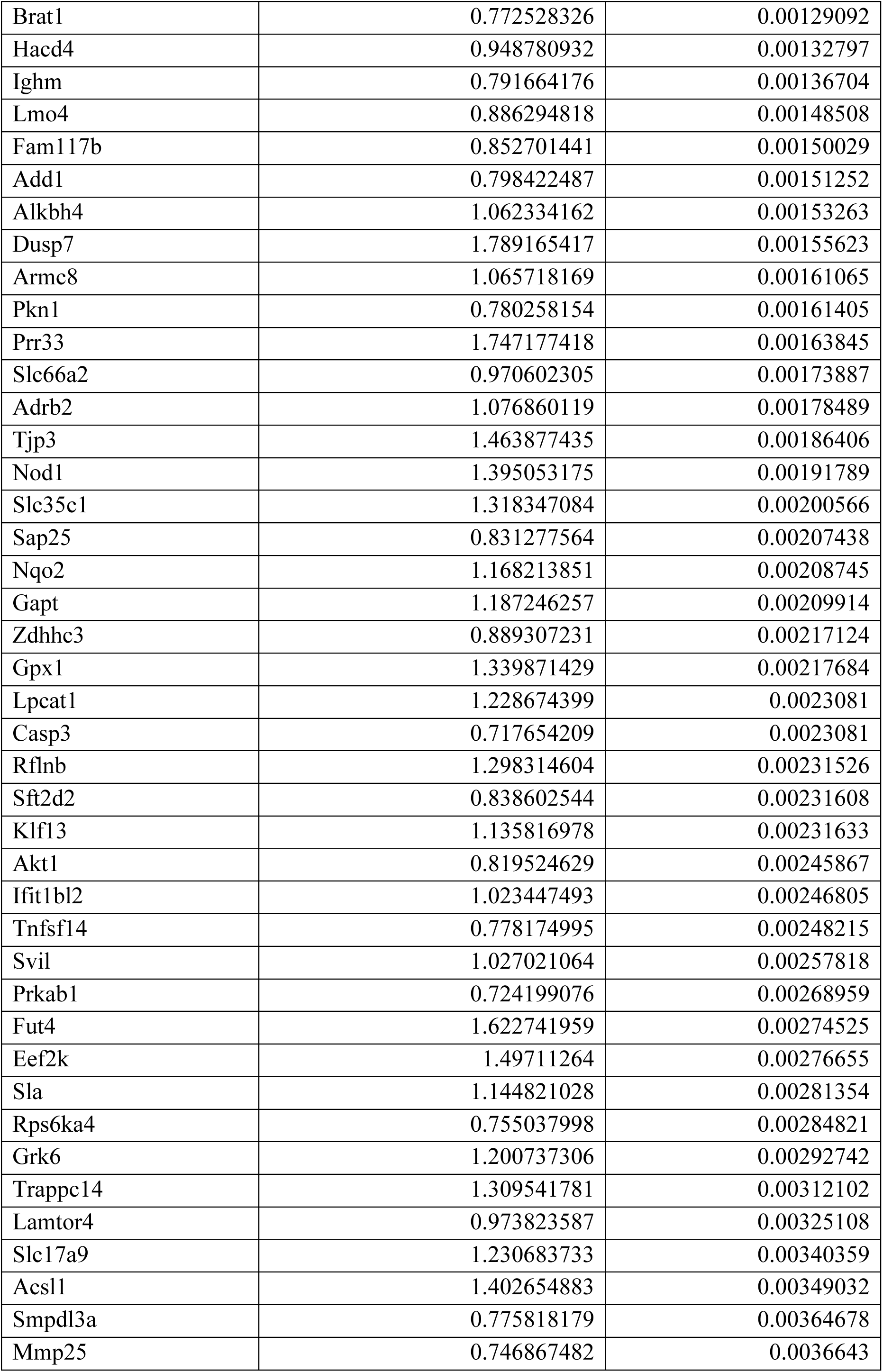
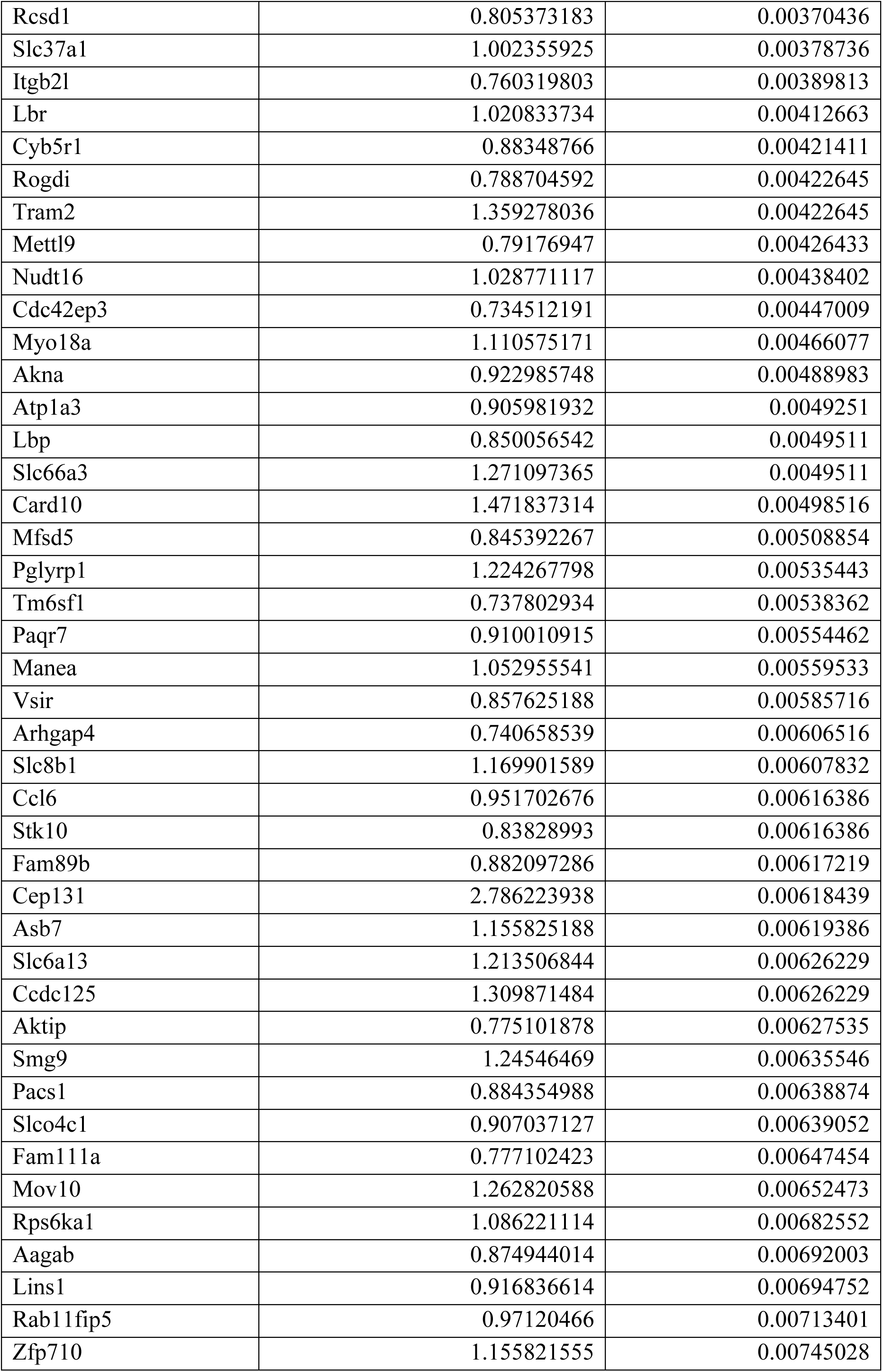
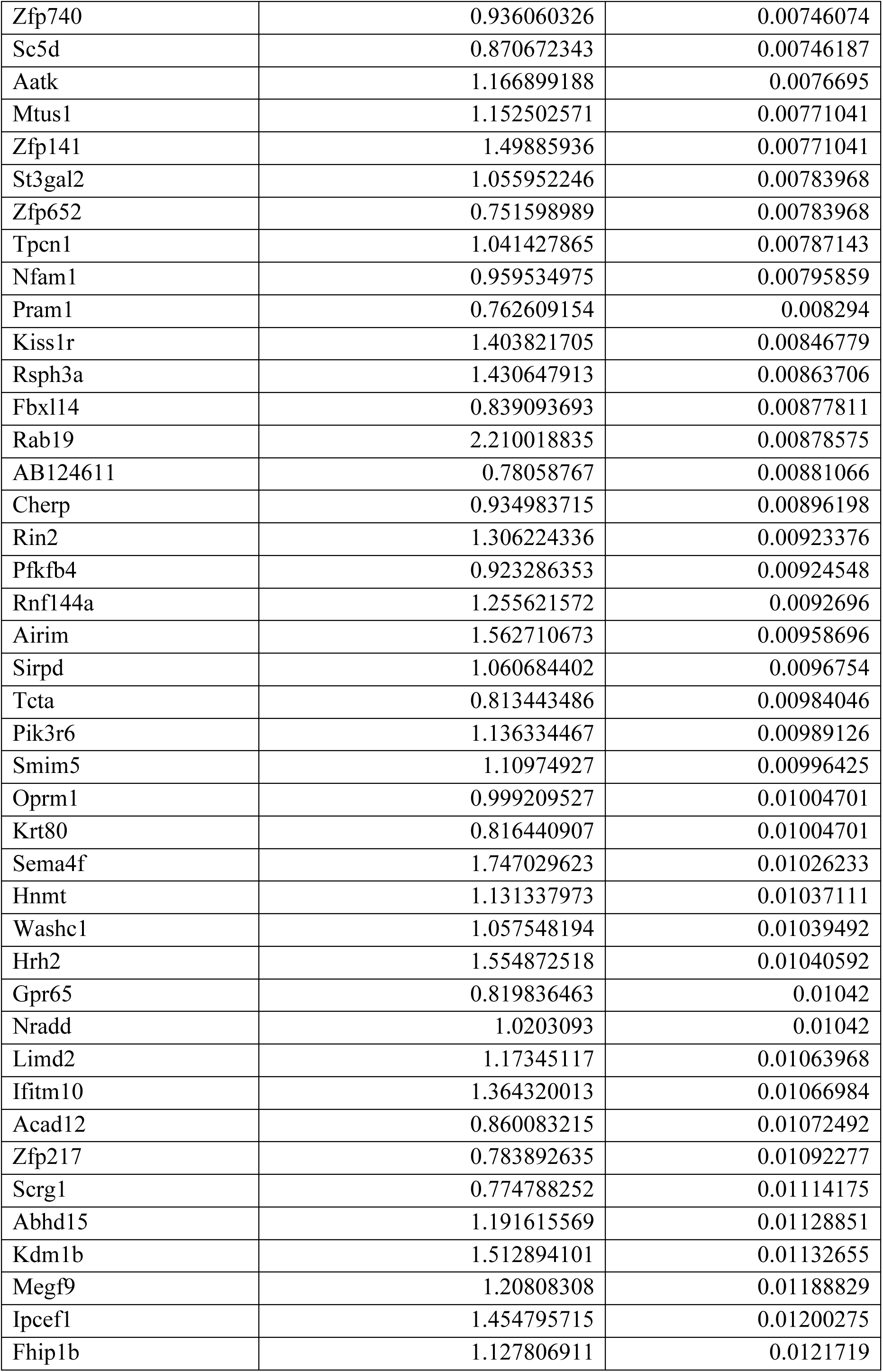
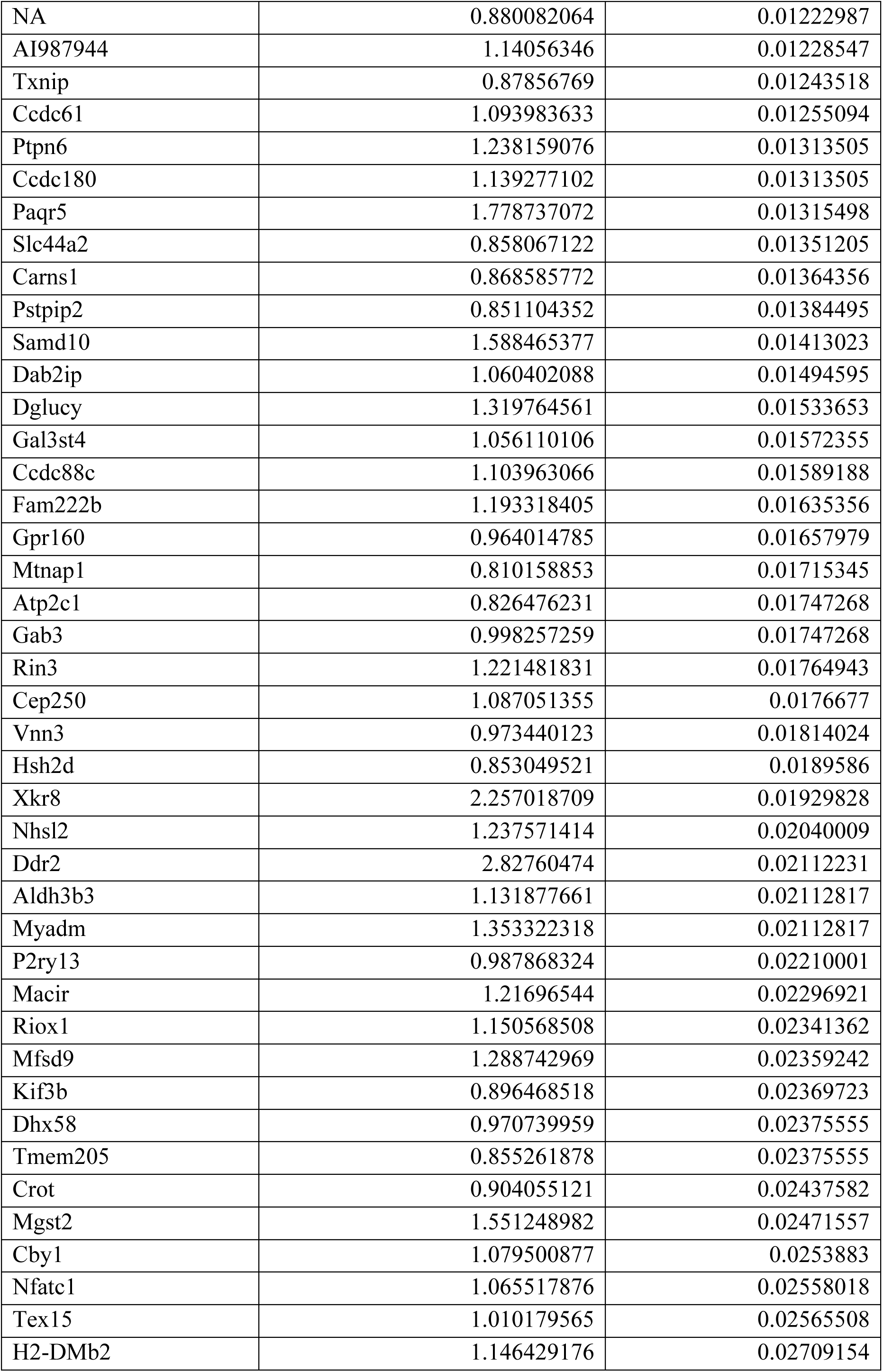
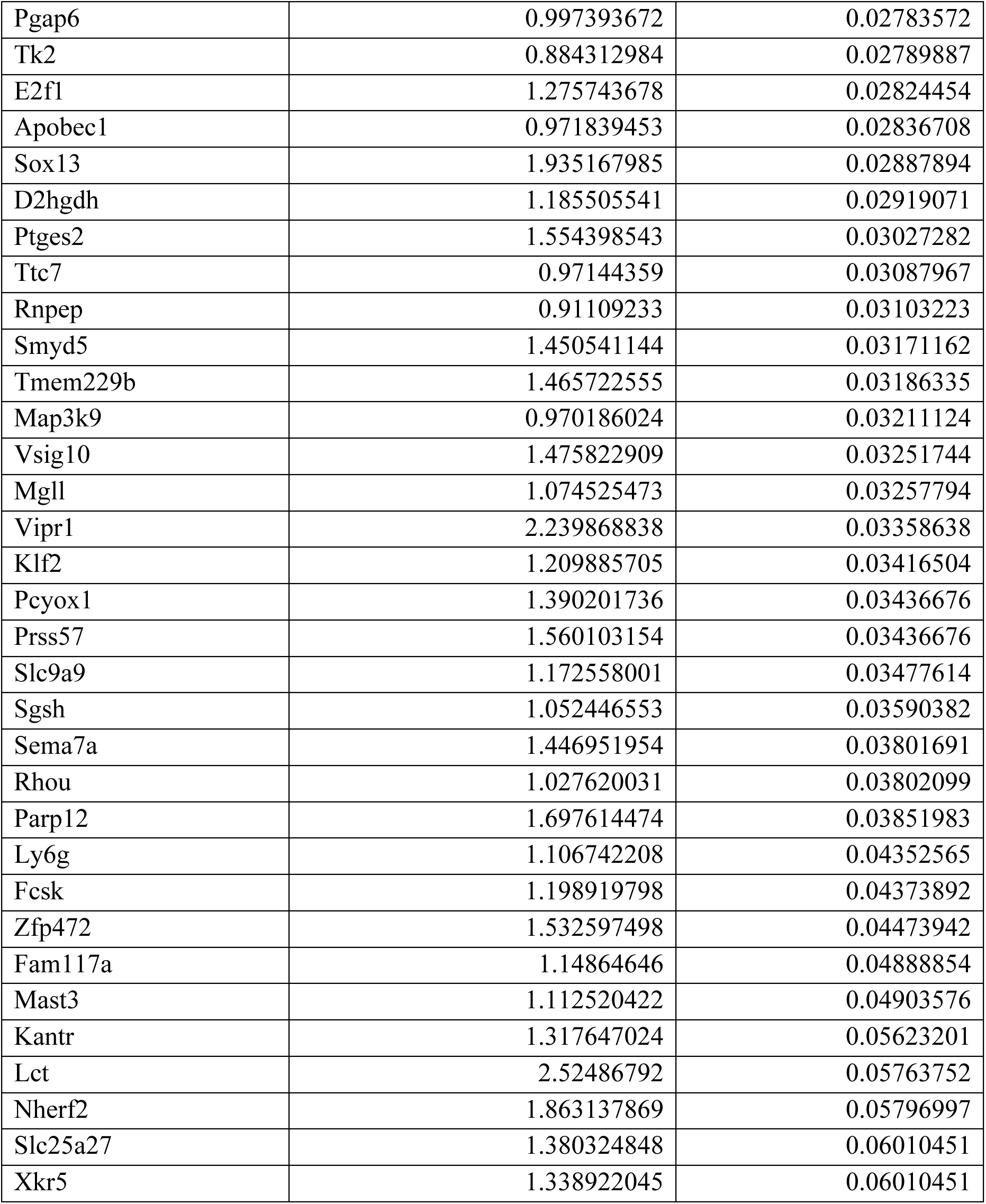
Transcripts with high abundance in co-cultured versus transwell control neutrophils. Displayed by P values, adjusted P-values <0.10 (Wald test, FDR-adjusted) and fold-change > 1.5.

**Supplementary Table 5.** Top 40 GO Cellular Biological Process annotation terms enriched in co-cultured versus transwell control neutrophils. Displayed by adjusted P values (FDR).

| **GO annotation term ID** | **GO annotation term, Biological Process Ontology** | **Fold-enrichment** | **Adj. P value** |
| --- | --- | --- | --- |
| GO:0008610 | lipid biosynthetic process | 1.88967213 | 0.03721421 |
| GO:0030833 | regulation of actin filament polymerization | 3.07386667 | 0.03721421 |
| GO:0050900 | leukocyte migration | 2.13552842 | 0.04090195 |
| GO:0030041 | actin filament polymerization | 2.78111746 | 0.04674717 |
| GO:0002685 | regulation of leukocyte migration | 2.42396343 | 0.05351982 |
| GO:0002697 | regulation of immune effector process | 1.9643645 | 0.05351982 |
| GO:0008154 | actin polymerization or depolymerization | 2.52646575 | 0.05612883 |
| GO:0008064 | regulation of actin polymerization or depolymerization | 2.67723871 | 0.05612883 |
| GO:0030832 | regulation of actin filament length | 2.6558208 | 0.05612883 |
| GO:0043123 | positive regulation of canonical NF-kappaB signal transduction | 2.49232432 | 0.05612883 |
| GO:0002683 | negative regulation of immune system process | 1.79724919 | 0.05612883 |
| GO:0006643 | membrane lipid metabolic process | 2.44280795 | 0.05612883 |
| GO:0006664 | glycolipid metabolic process | 3.15475789 | 0.05612883 |
| GO:0046467 | membrane lipid biosynthetic process | 2.73232593 | 0.05612883 |
| GO:1903509 | liposaccharide metabolic process | 3.11378701 | 0.05612883 |
| GO:0042119 | neutrophil activation | 4.14972 | 0.05612883 |
| GO:0007015 | actin filament organization | 1.90020848 | 0.05612883 |
| GO:0002695 | negative regulation of leukocyte activation | 2.37976774 | 0.05742758 |
| GO:0006935 | chemotaxis | 1.96902491 | 0.05742758 |
| GO:0042330 | taxis | 1.96902491 | 0.05742758 |
| GO:0042554 | superoxide anion generation | 4.47107879 | 0.05742758 |
| GO:0032271 | regulation of protein polymerization | 2.34945223 | 0.05831347 |
| GO:0050866 | negative regulation of cell activation | 2.27827765 | 0.06101897 |
| GO:0043408 | regulation of MAPK cascade | 1.72519911 | 0.06317253 |
| GO:0051250 | negative regulation of lymphocyte activation | 2.44101176 | 0.06317253 |
| GO:0032535 | regulation of cellular component size | 1.92786063 | 0.06360936 |
| GO:0022407 | regulation of cell-cell adhesion | 1.78638325 | 0.06360936 |
| GO:0097529 | myeloid leukocyte migration | 2.23876994 | 0.06360936 |
| GO:0043410 | positive regulation of MAPK cascade | 1.89317616 | 0.06536883 |
| GO:0032956 | regulation of actin cytoskeleton organization | 1.9635346 | 0.06574797 |
| GO:0032970 | regulation of actin filament-based process | 1.90791724 | 0.06598831 |
| GO:0007159 | leukocyte cell-cell adhesion | 1.83874804 | 0.06636462 |
| GO:0050672 | negative regulation of lymphocyte proliferation | 3.12115692 | 0.0827824 |
| GO:0007249 | canonical NF-kappaB signal transduction | 1.94518125 | 0.08407461 |
| GO:0009247 | glycolipid biosynthetic process | 3.29342857 | 0.08517106 |
| GO:0036230 | granulocyte activation | 3.5316766 | 0.08517106 |
| GO:0006687 | glycosphingolipid metabolic process | 3.88277895 | 0.08517106 |
| GO:0022408 | negative regulation of cell-cell adhesion | 2.23197962 | 0.08517106 |
| GO:0110053 | regulation of actin filament organization | 2.04924444 | 0.08517106 |
| GO:0097530 | granulocyte migration | 2.51498182 | 0.08517106 |

**Supplementary Table 6.**
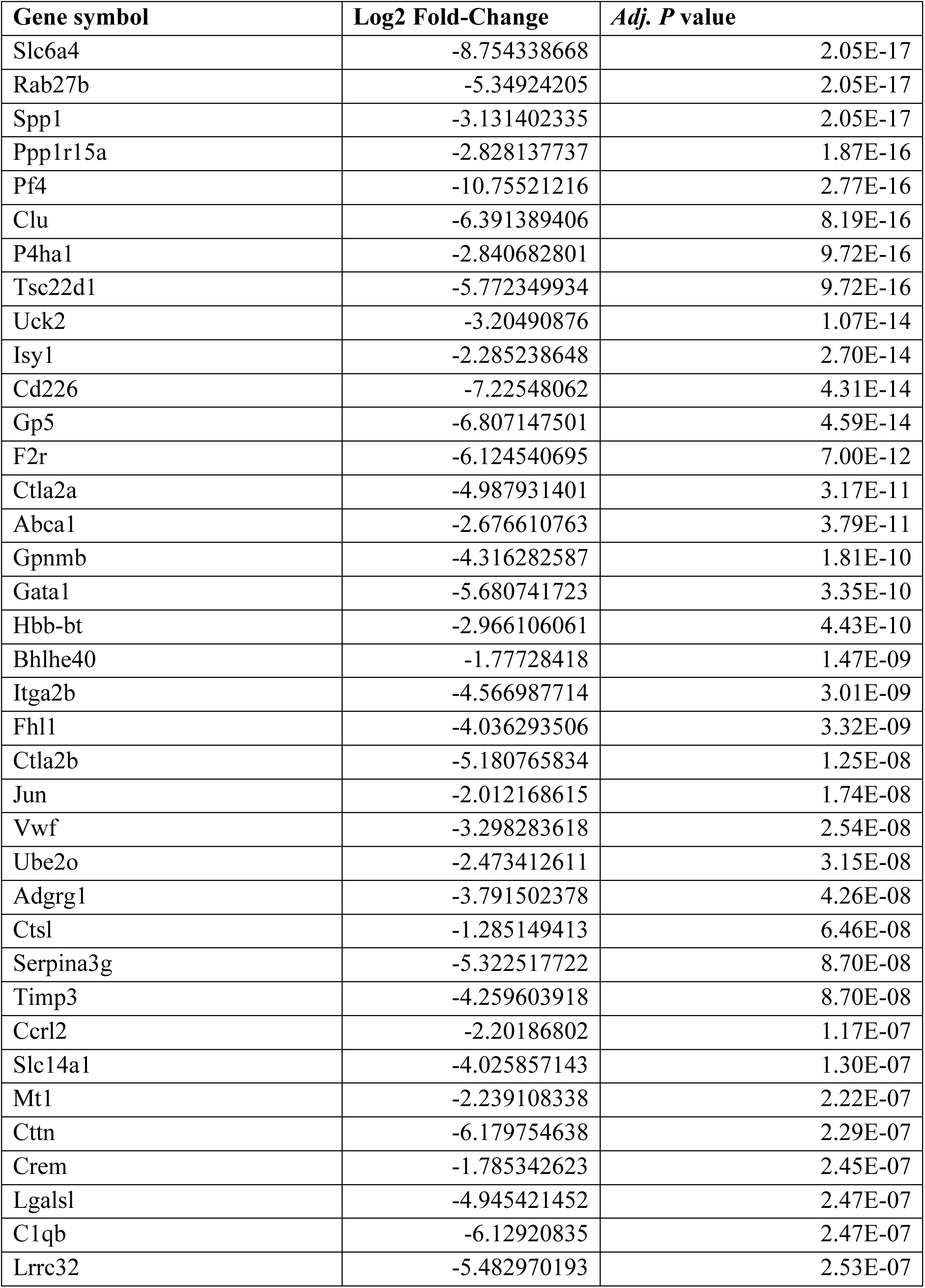
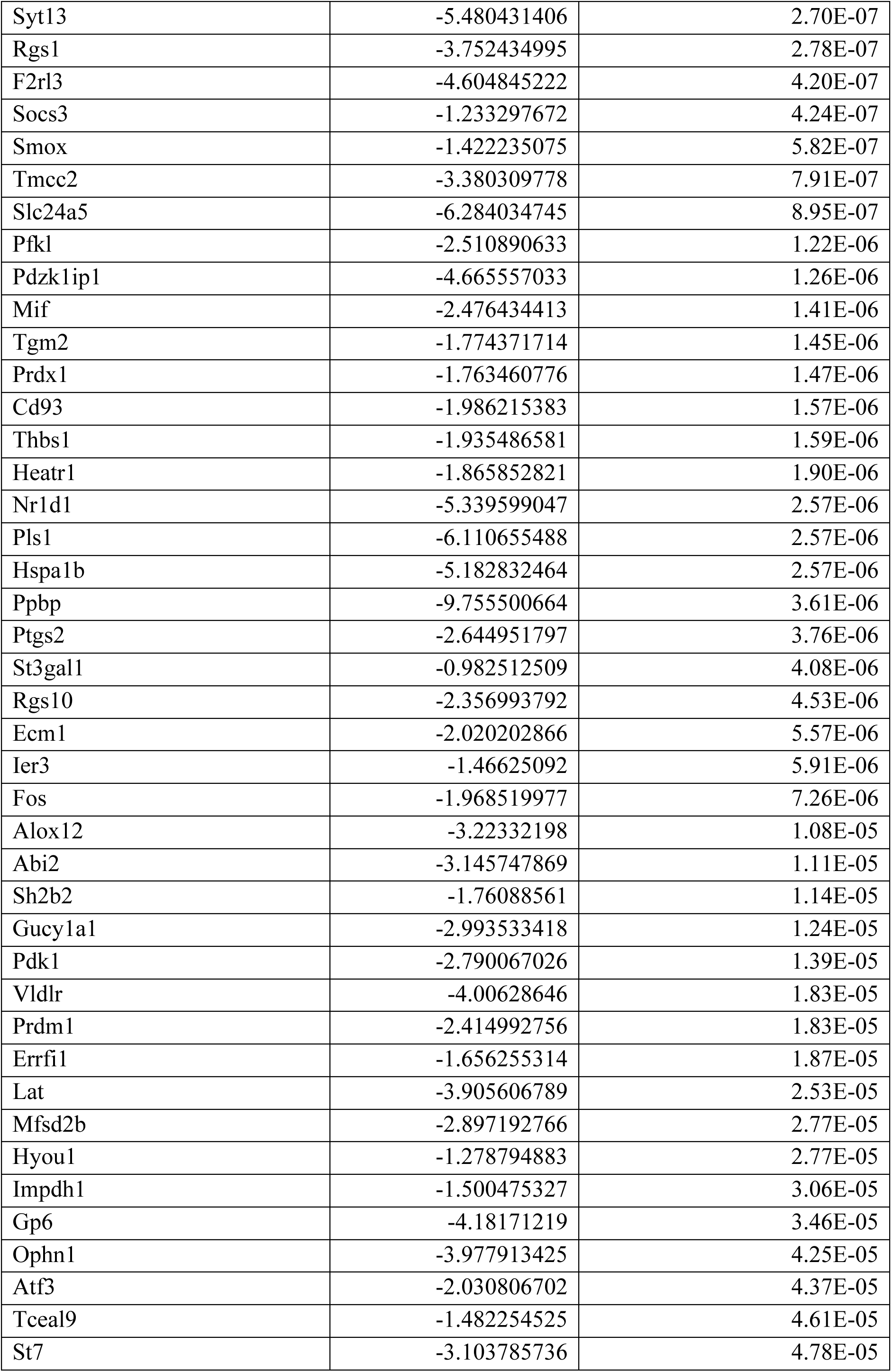
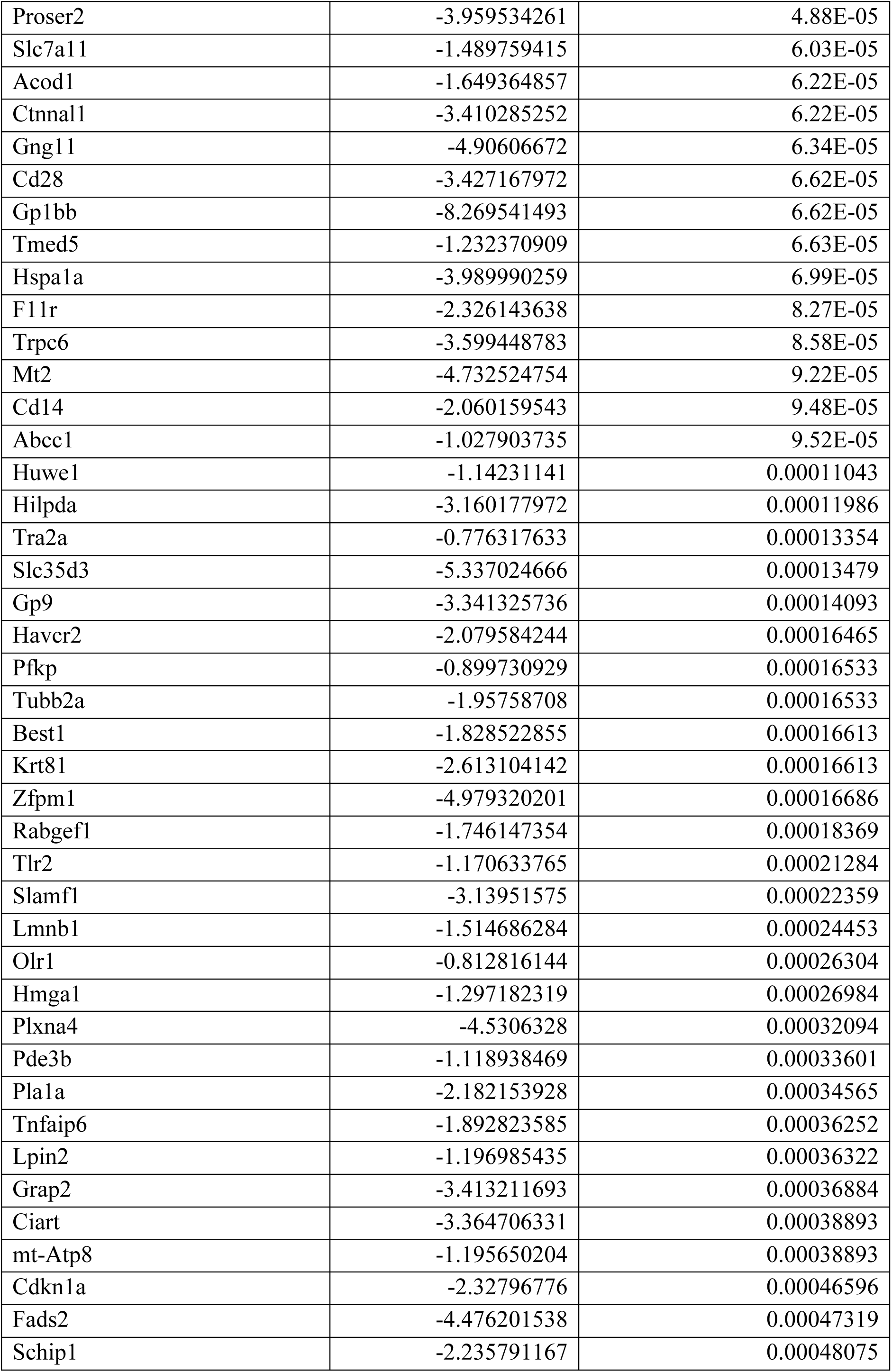
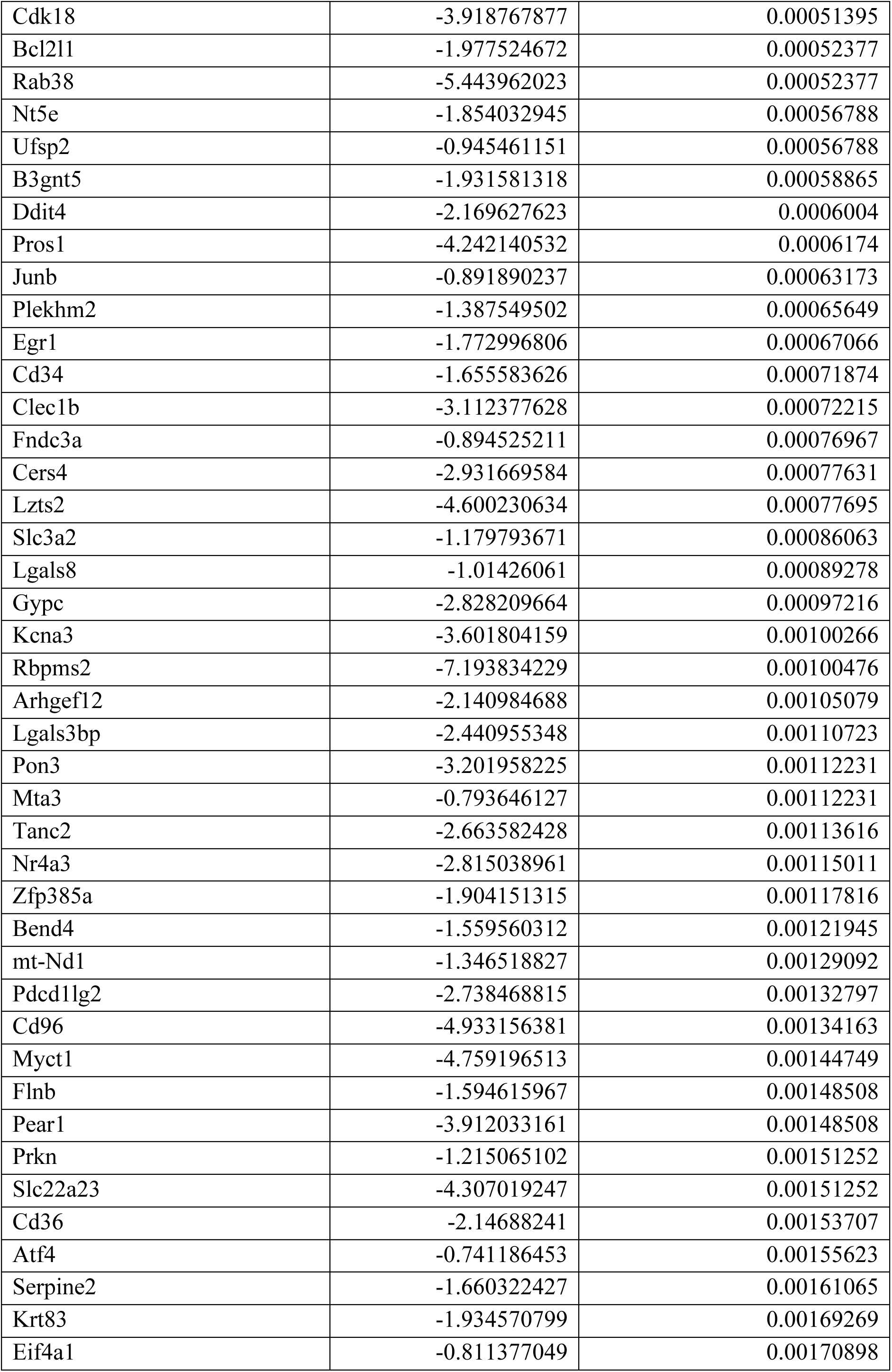
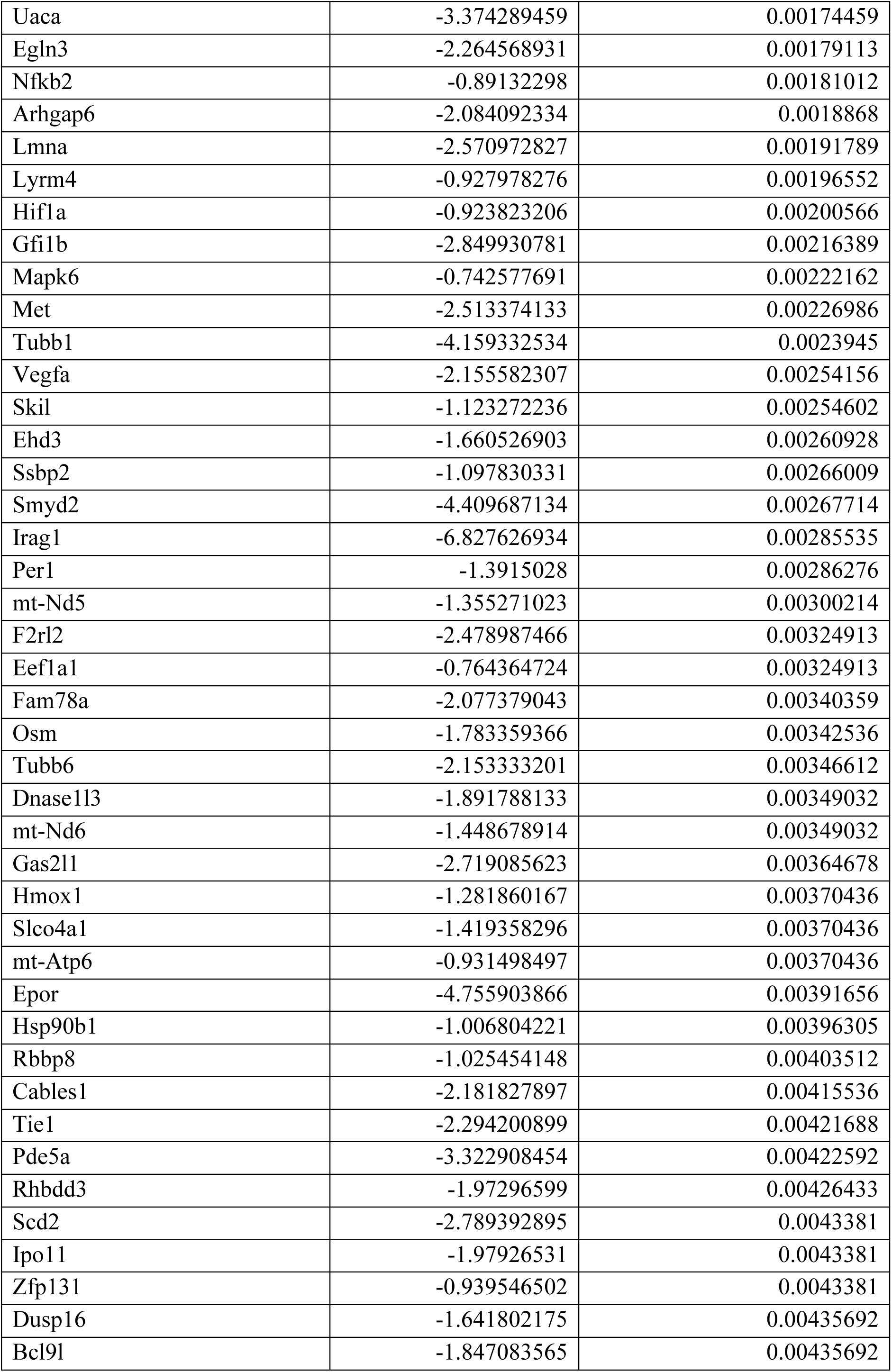
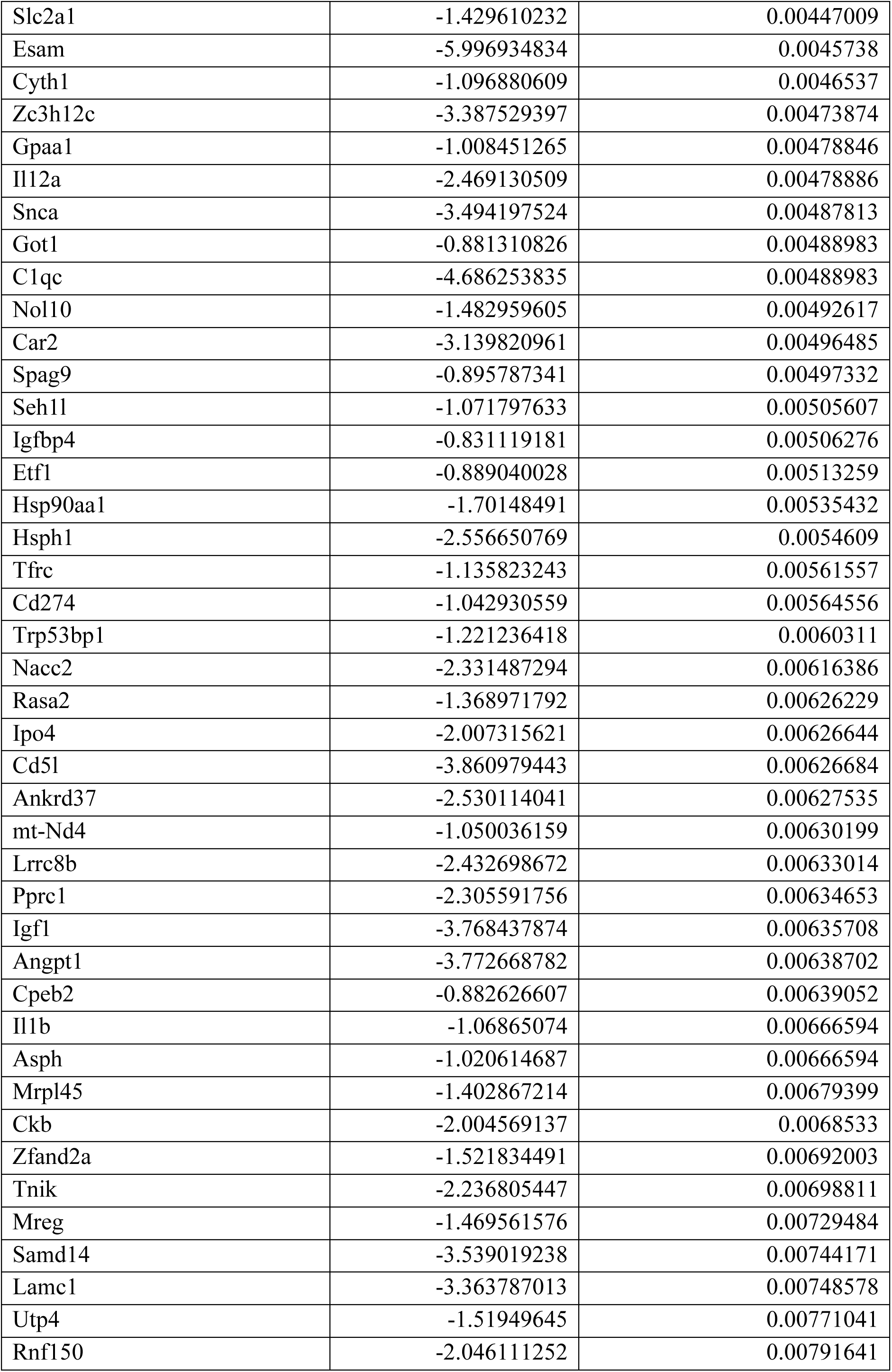
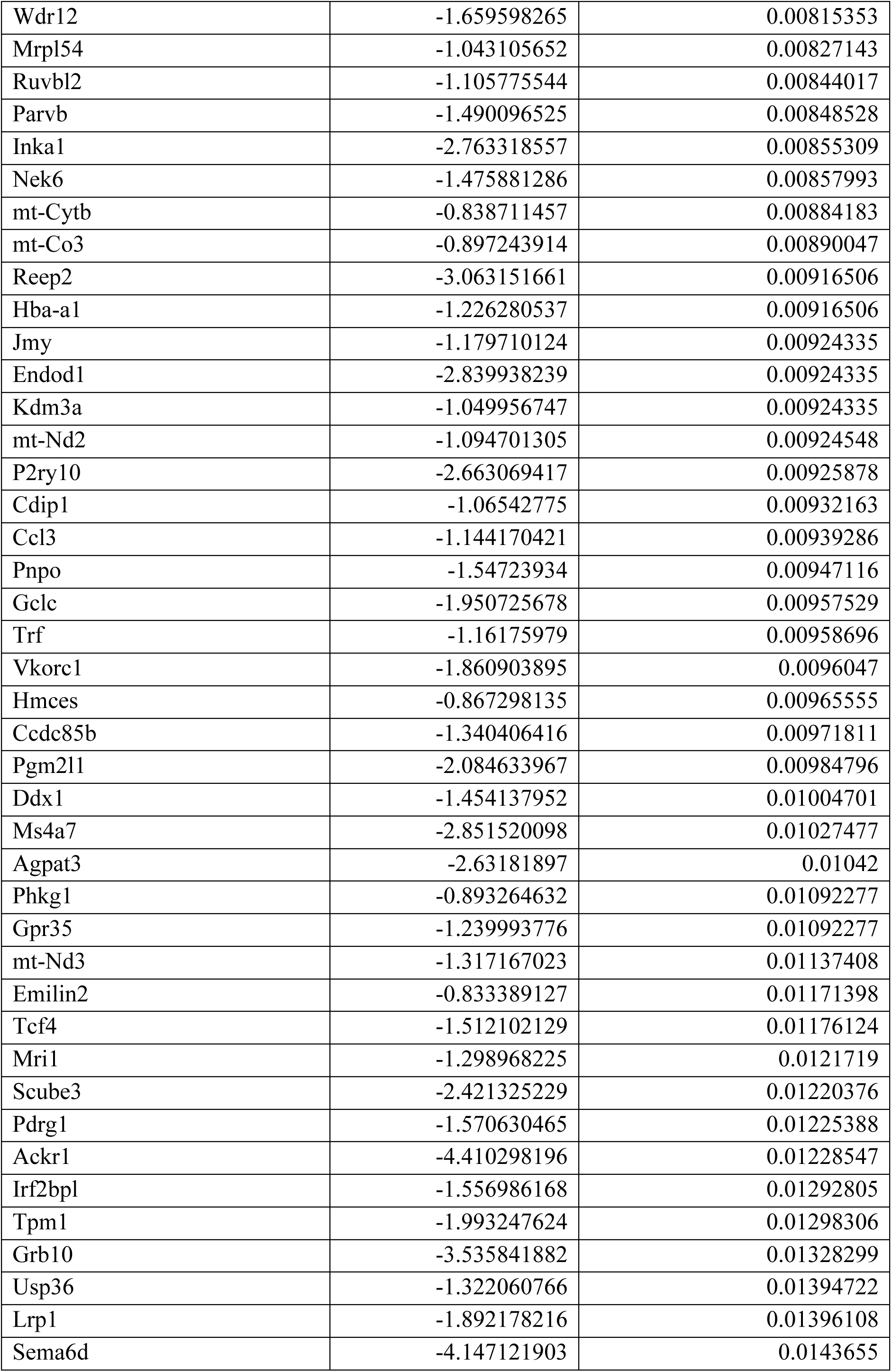
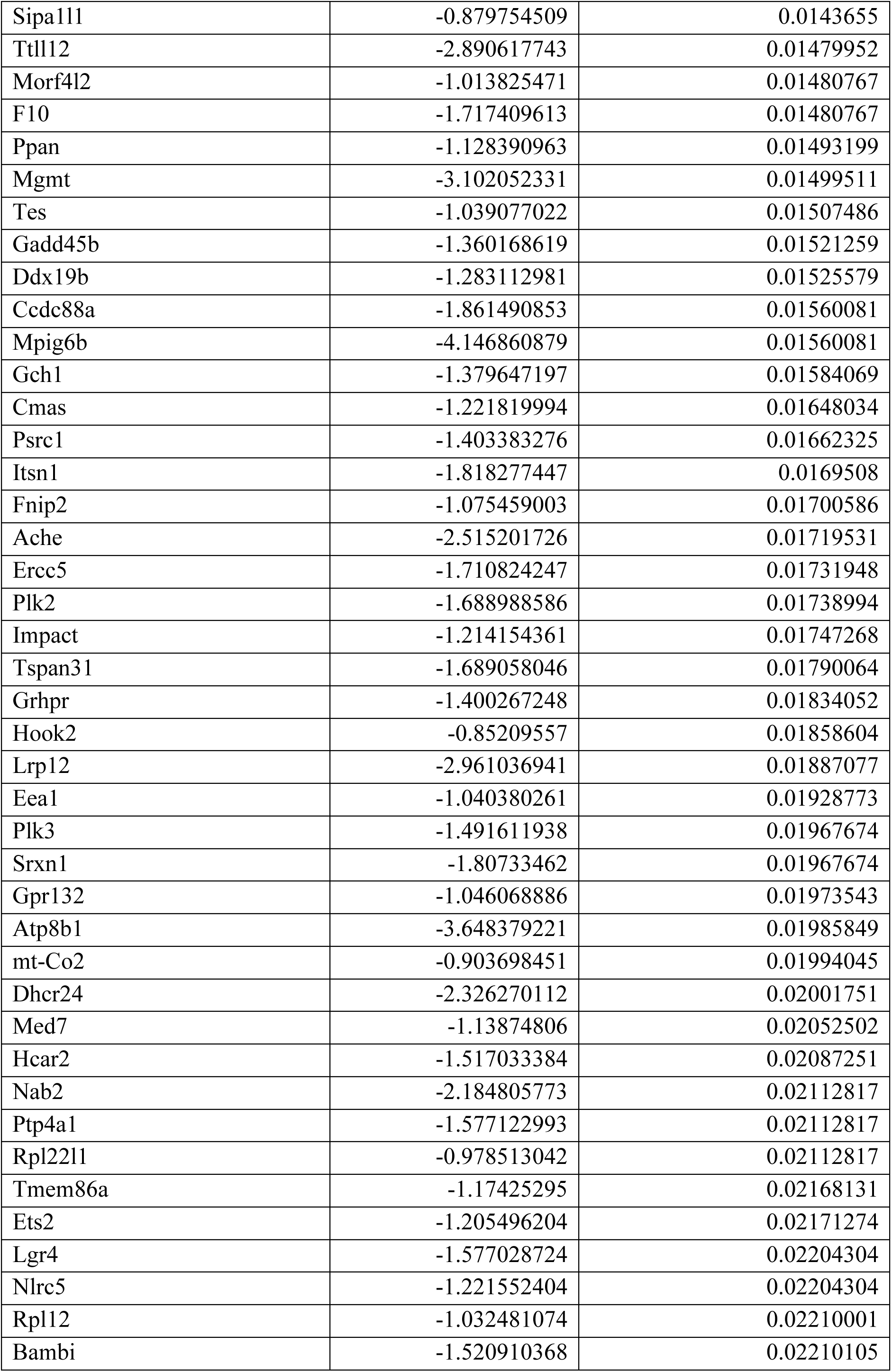
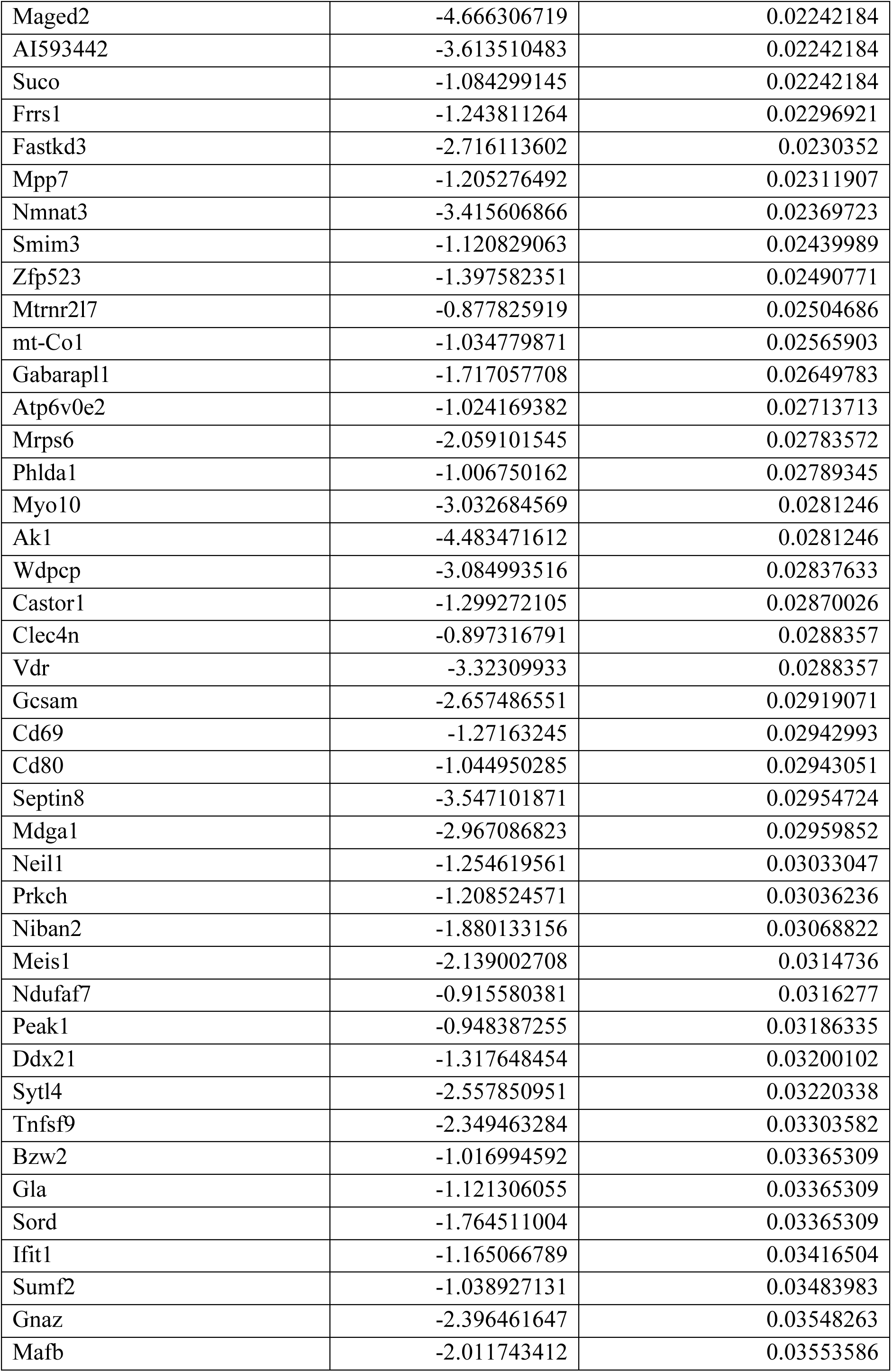
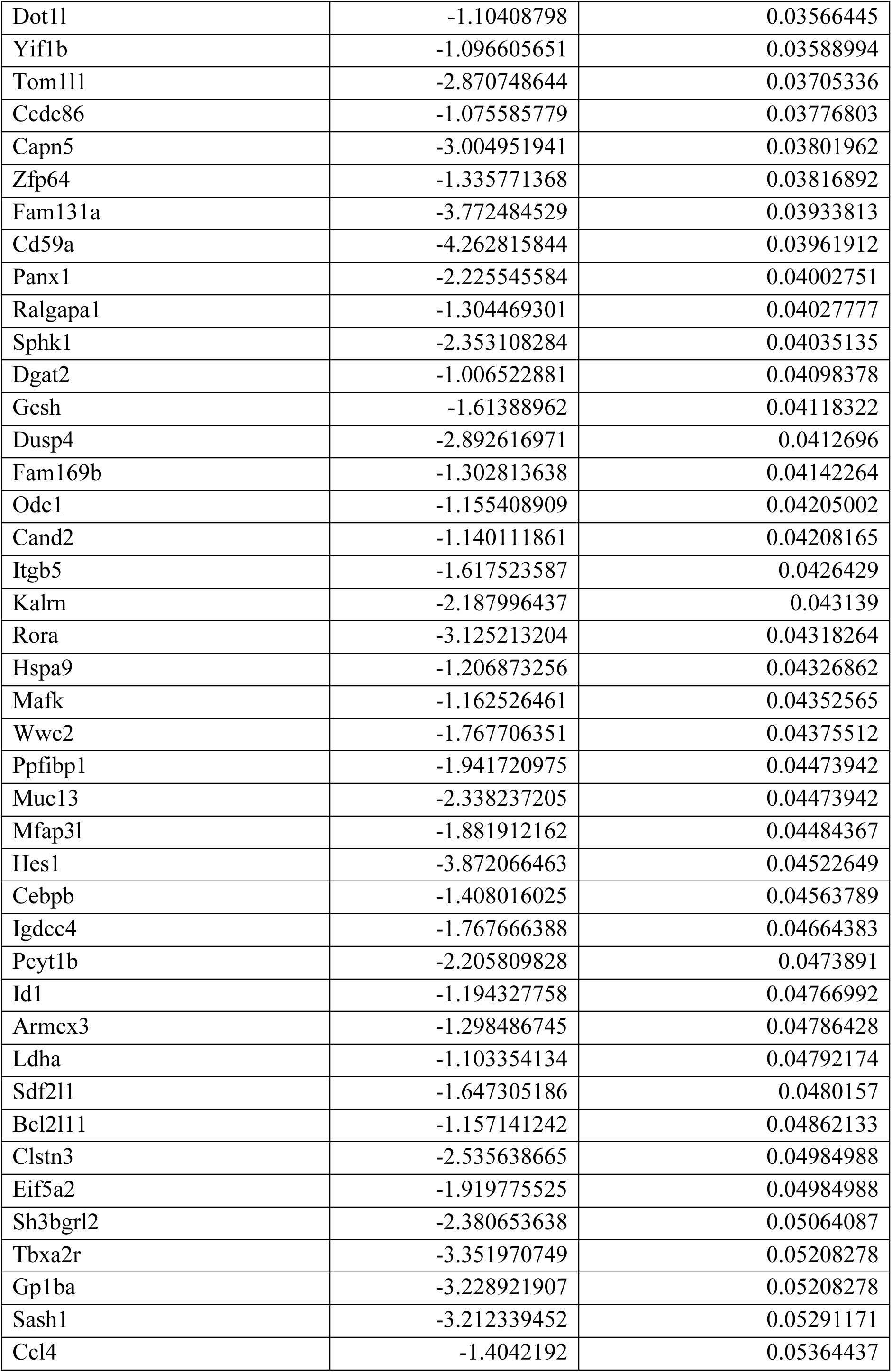
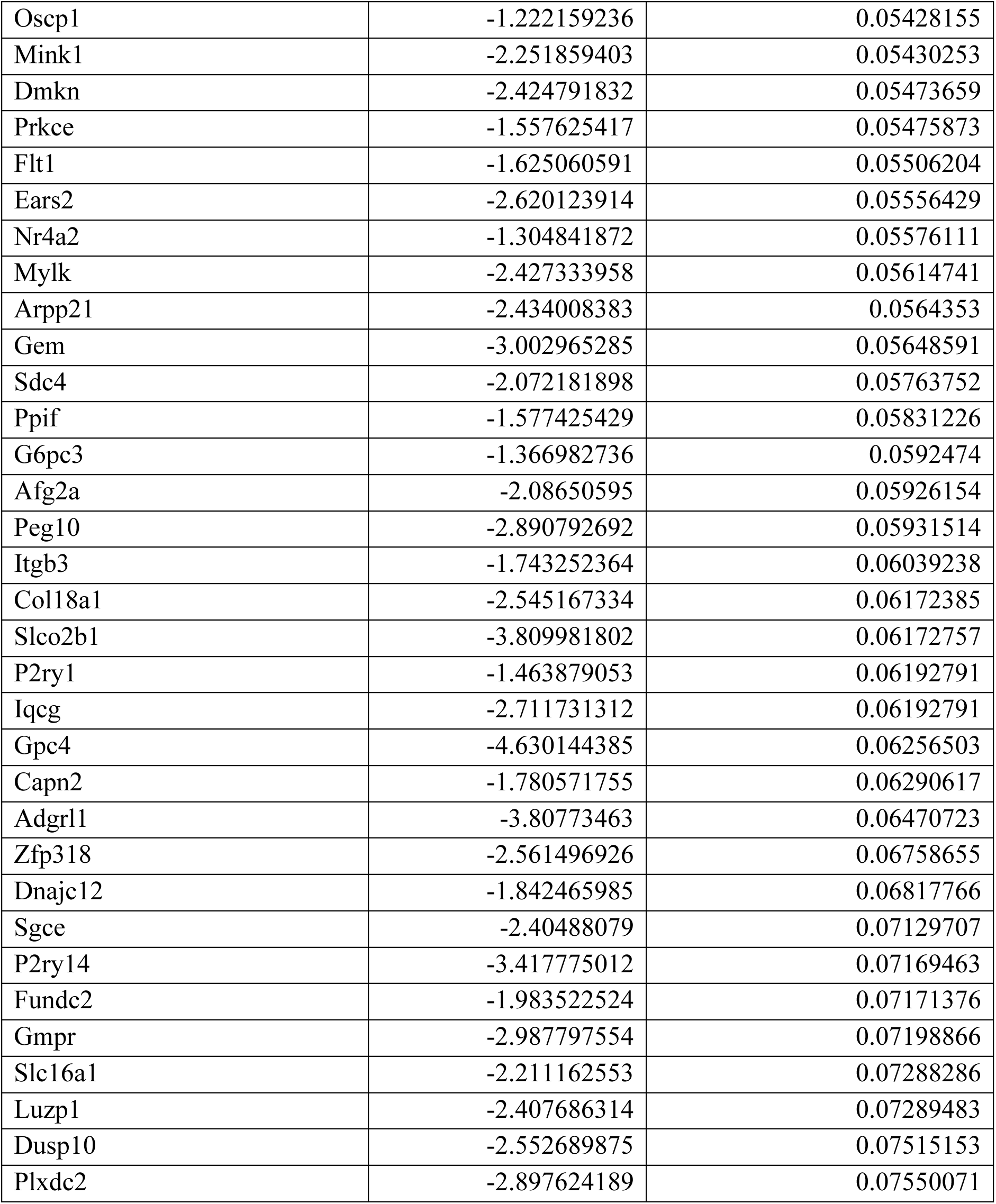
Transcripts with low abundance in co-cultured versus transwell control neutrophils. Displayed by P values, adjusted P-values <0.10 (Wald test, FDR-adjusted) and fold-change < -1.5.

**Supplementary Table 7.** Top 40 GO Biological Process Ontology annotation terms enriched in transwell control versus co-culture neutrophils. Displayed by adjusted P values (FDR).

| **GO annotation term ID** | **GO annotation term, Biological Process Ontology** | **Fold-enrichment** | **Adj. P value** |
| --- | --- | --- | --- |
| GO:0007599 | hemostasis | 5.08889518 | 3.44E-14 |
| GO:0007596 | blood coagulation | 5.06698902 | 5.40E-14 |
| GO:0050817 | coagulation | 4.99408271 | 6.16E-14 |
| GO:0030168 | platelet activation | 6.23779521 | 1.12E-12 |
| GO:0042060 | wound healing | 3.43331535 | 1.12E-12 |
| GO:0050878 | regulation of body fluid levels | 3.74929238 | 1.23E-12 |
| GO:0009611 | response to wounding | 2.97951219 | 5.16E-12 |
| GO:0008285 | negative regulation of cell population proliferation | 2.51832214 | 4.01E-09 |
| GO:0070527 | platelet aggregation | 6.08927628 | 3.73E-07 |
| GO:0001666 | response to hypoxia | 2.99075074 | 7.66E-07 |
| GO:0048514 | blood vessel morphogenesis | 2.37701869 | 8.64E-07 |
| GO:0010543 | regulation of platelet activation | 6.37249843 | 1.38E-06 |
| GO:0001568 | blood vessel development | 2.21314559 | 1.98E-06 |
| GO:0036293 | response to decreased oxygen levels | 2.82807392 | 2.64E-06 |
| GO:0070482 | response to oxygen levels | 2.72967557 | 2.68E-06 |
| GO:0030193 | regulation of blood coagulation | 5.9476652 | 8.96E-06 |
| GO:1900046 | regulation of hemostasis | 5.9476652 | 8.96E-06 |
| GO:0001525 | angiogenesis | 2.34726292 | 1.28E-05 |
| GO:0050818 | regulation of coagulation | 5.68332453 | 1.54E-05 |
| GO:0061041 | regulation of wound healing | 4.18179214 | 1.74E-05 |
| GO:0034109 | homotypic cell-cell adhesion | 4.37398719 | 4.01E-05 |
| GO:0040017 | positive regulation of locomotion | 2.12707596 | 4.76E-05 |
| GO:0030099 | myeloid cell differentiation | 2.18123329 | 4.94E-05 |
| GO:1903530 | regulation of secretion by cell | 2.12911758 | 5.57E-05 |
| GO:0097193 | intrinsic apoptotic signaling pathway | 2.42719404 | 5.57E-05 |
| GO:0043523 | regulation of neuron apoptotic process | 2.62103631 | 5.57E-05 |
| GO:2001233 | regulation of apoptotic signaling pathway | 2.27685983 | 5.57E-05 |
| GO:0048660 | regulation of smooth muscle cell proliferation | 3.20020359 | 5.72E-05 |
| GO:0030335 | positive regulation of cell migration | 2.1312467 | 5.93E-05 |
| GO:0043408 | regulation of MAPK cascade | 2.07496497 | 7.55E-05 |
| GO:0071363 | cellular response to growth factor stimulus | 2.06576335 | 8.37E-05 |
| GO:2000147 | positive regulation of cell motility | 2.08653523 | 0.00010136 |
| GO:0070848 | response to growth factor | 2.02975876 | 0.00010455 |
| GO:0048659 | smooth muscle cell proliferation | 3.06592932 | 0.00010619 |
| GO:0043065 | positive regulation of apoptotic process | 1.984006 | 0.00015295 |
| GO:0007169 | cell surface receptor protein tyrosine kinase signaling pathway | 2.08395135 | 0.00015295 |
| GO:0032496 | response to lipopolysaccharide | 2.33072299 | 0.00015295 |
| GO:0043410 | positive regulation of MAPK cascade | 2.29860098 | 0.00015295 |
| GO:0043524 | negative regulation of neuron apoptotic process | 2.98250267 | 0.00015477 |
| GO:0071219 | cellular response to molecule of bacterial origin | 2.64883518 | 0.00015609 |

**Supplementary Tables 8-10: Proteomic analysis of neutrophils following co-culture with MKs.** Murine marrow neutrophils were co-cultured with MKs for 3 hours (emperipolesis) and control neutrophils were co-cultured with MKs separated by 0.45 µm transwell inserts (control). Neutrophils were FACS-sorted as single, live, CD45+CD11b+Ly6G+ cells, lysed, and TCA-precipitated proteins identified by mass spectrometry. Results were analyzed for differentially abundant proteins using proDA (version 1.20.0)^110^ (**S8, S9**), followed by analysis for enriched pathways using PathfindR (version 2.4.1)^111^ based on mouse KEGG pathway terms (**S10**). N=3 paired samples.

**Supplementary Table 8.**
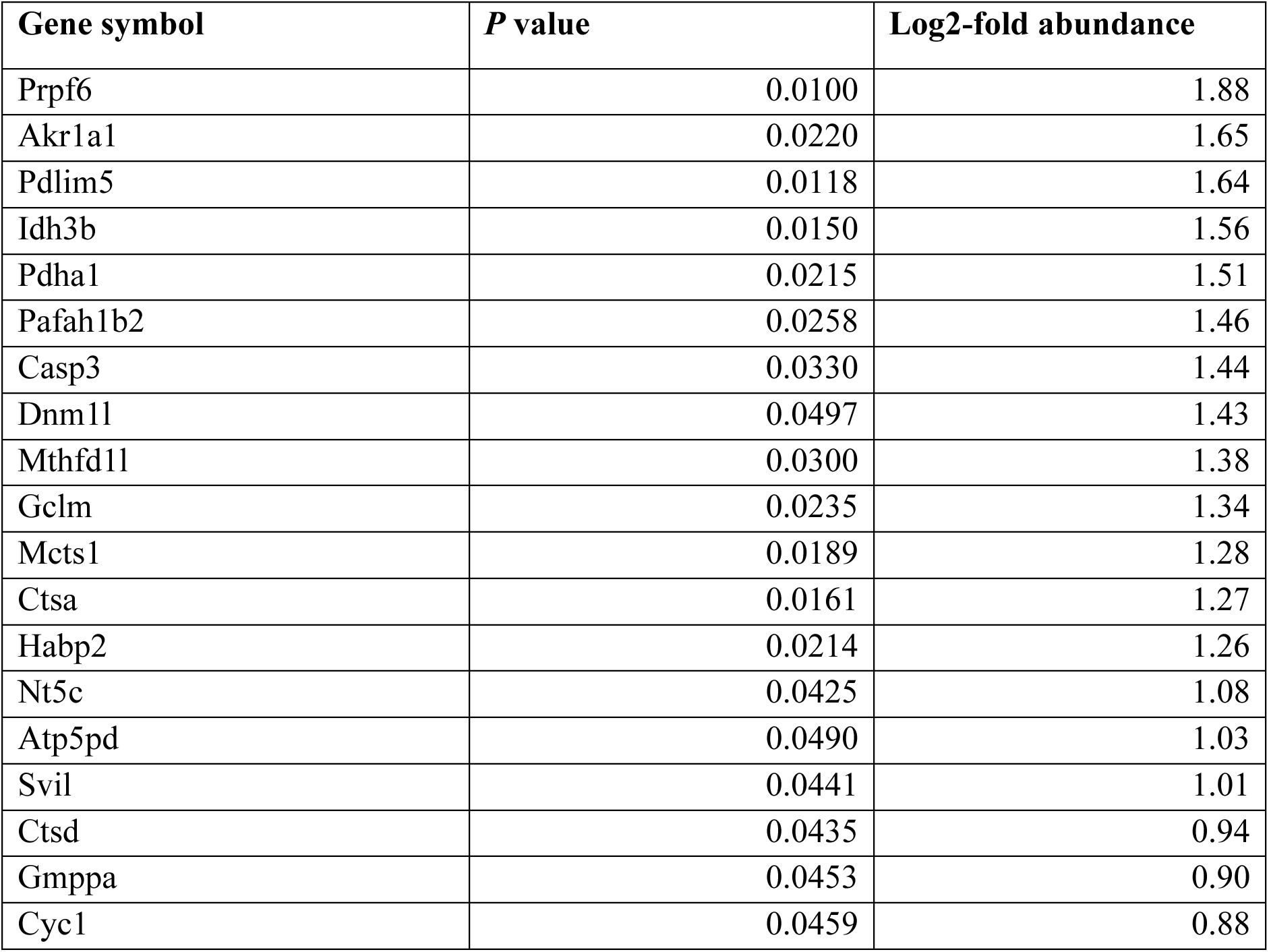
Proteins with high abundance in co-cultured versus transwell control neutrophils. Nominal P-values <0.05 (Wald-test, after a linear probabilistic dropout model fit to median-normalized protein abundance data).

**Supplementary Table 9.**
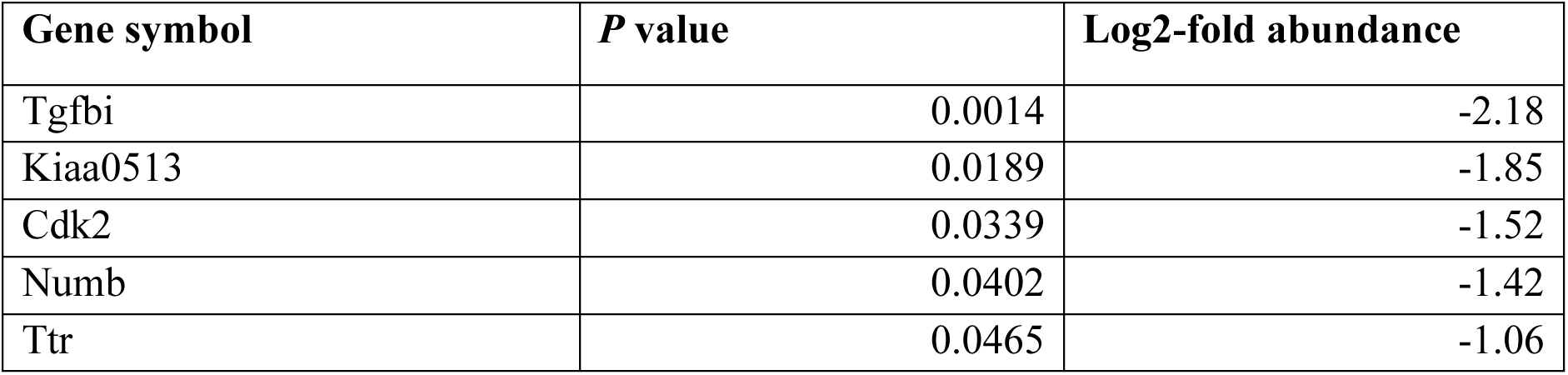
Proteins with low abundance in co-cultured versus transwell control neutrophils. Nominal P-values <0.05 (Wald-test, after a linear probabilistic dropout model fit to median-normalized protein abundance data).

**Supplementary Table 10.**
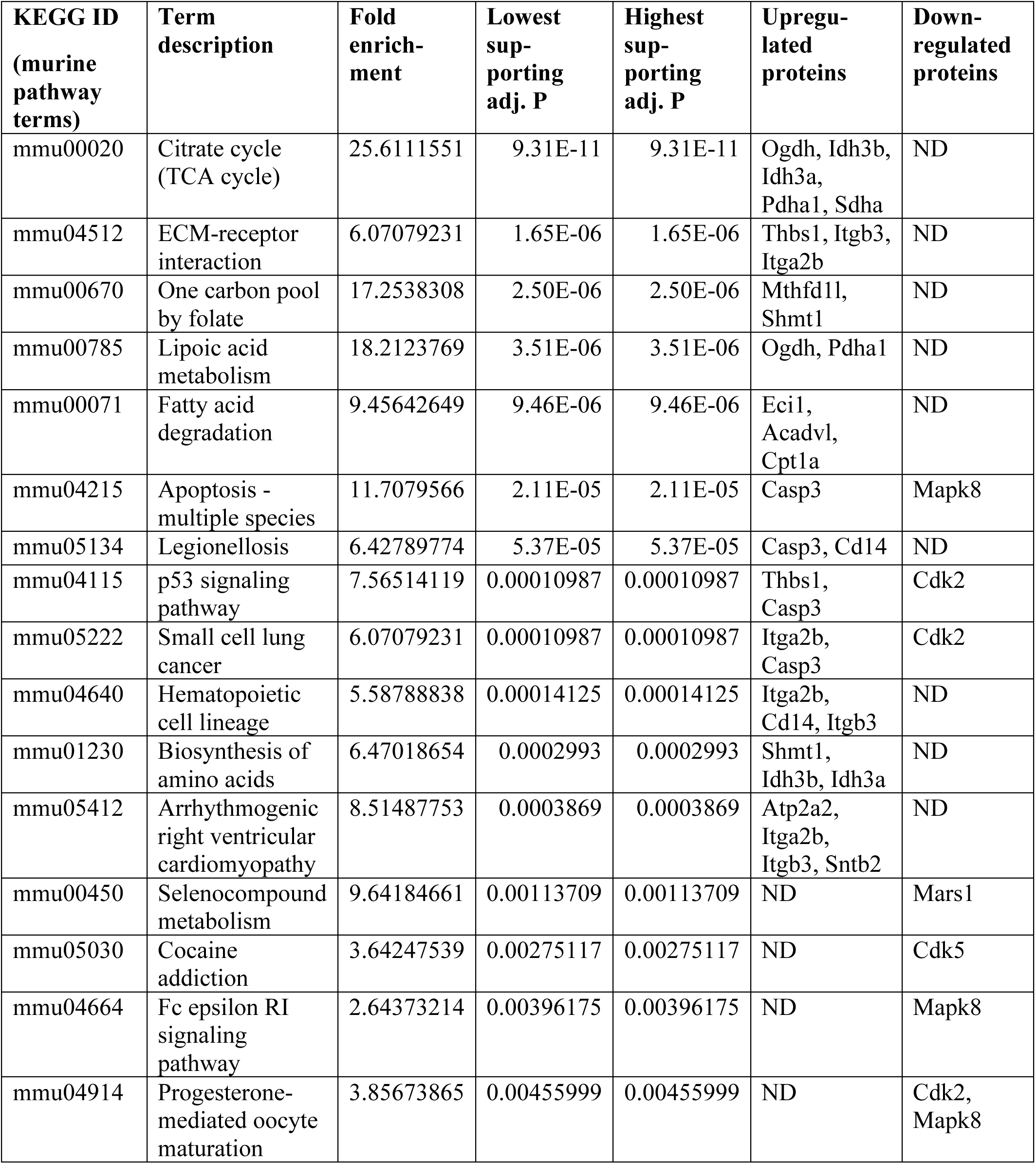
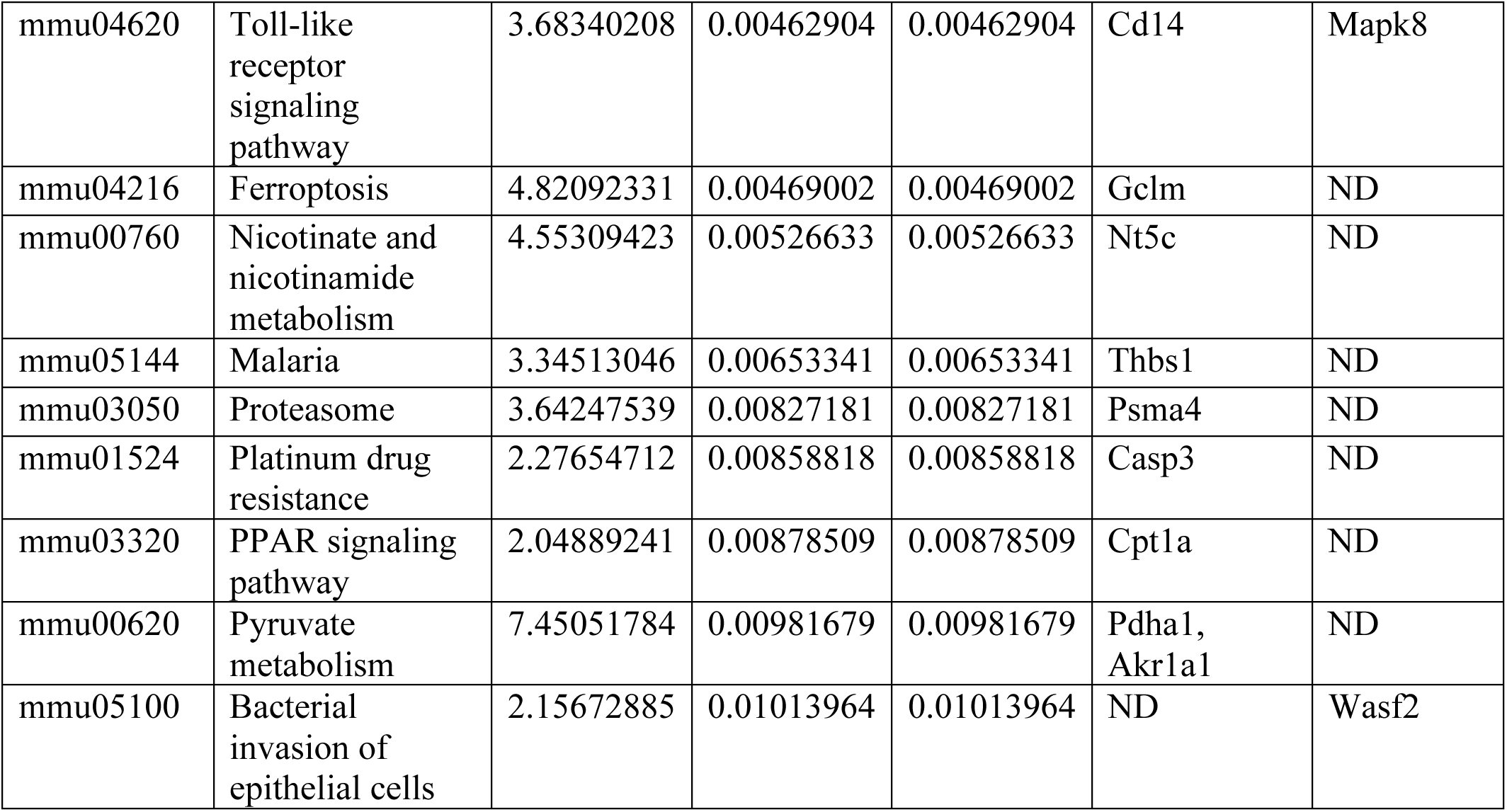
Representative members of the 25 most enriched pathway clusters in co-cultured versus transwell control neutrophils. Selected by lowest reported supporting P value. Lowest and highest supporting adj. P refer to the P values occurred over 10 iterations of active-subnetwork-enrichment analysis, calculated via one-sided hypergeometric testing, FDR-adjusted (α<0.10). Terms were clustered by hierarchal clustering methods.

**Supplementary Tables 11-17: Transcriptomic analysis of circulating neutrophils in MK-depleted mice.** PF4-Cre+ x iDTR^fl/fl^ mice vs. PF4-Cre- x iDTR^fl/fl^ littermate control mice were injected for 3 days with 250 ng diphtheria toxin (DTx) i.p. to deplete MKs, followed by treatment with vehicle or with LPS (2 mg/kg i.p., O111:B4, E. coli) to induce peritonitis. Two hours after induction of peritonitis, blood and marrow neutrophils were FACS-sorted as single, live, CD45+CD11b+Ly6G+ cells, lysed, and sent for bulk transcriptomic analysis. Differential gene expression analysis was performed with DSeq2 (v3.20)^107^ (Wald test, FDR: Padj<0.10) using a full genotype x treatment model (genotype, treatment, and genotypextreatment interaction); results shown report the main genotype effect (PF4-Cre+ vs PF4-Cre-) adjusted for treatment (i.e., averaged across LPS/vehicle treatment within genotype), while interaction terms were included to allow genotype effects to differ by treatment but are not shown to focus on the overall genotype effect across both conditions (**S11,S12,S15,S16**). Gene Ontology (GO) term enrichment was done using clusterProfiler (v4.14.3)^108^ (FDR: Padj<0.10), with annotation based on org.Mm.eg.db (v3.20.0) (**S13,14,S17**). N=3-5 (F/M) / genotype and treatment group.

**Supplementary Table 11.** Transcripts with high abundance in blood neutrophils in MK-sufficient versus MK-depleted mice. Displayed by P values, adjusted P-values <0.10 (Wald test, FDR-adjusted) and fold-change > 1.5.

| Gene symbol | Log2 Fold-Change | Adj. P value |
| --- | --- | --- |
| Pfn1 | 2.85950573 | 2.19E-05 |
| Zfp36 | 3.0628107 | 3.50E-05 |
| Nr4a3 | 4.60035268 | 6.87E-05 |
| Actb | 2.32301647 | 0.00029842 |
| Trib1 | 2.21923495 | 0.00046341 |
| Vsir | 2.06464964 | 0.0016882 |
| Degs1 | 2.40347026 | 0.00180981 |
| Fth1 | 1.15811897 | 0.00198301 |
| Sbno2 | 2.18489182 | 0.00198301 |
| Tapbp | 1.94346198 | 0.00218105 |
| Map4 | 2.65976919 | 0.00261416 |
| Jun | 2.91675295 | 0.00261416 |
| Ccl3 | 1.89825592 | 0.00284511 |
| Laptm5 | 1.62482556 | 0.0032252 |
| Rin3 | 3.30800924 | 0.00416517 |
| Stat1 | 1.14713147 | 0.0045277 |
| Itgb2 | 1.34781696 | 0.00456692 |
| Ptafr | 1.17311446 | 0.00531517 |
| Trex1 | 2.23639161 | 0.00877278 |
| Pycard | 2.47344943 | 0.00959274 |

**Supplementary Table 12.**
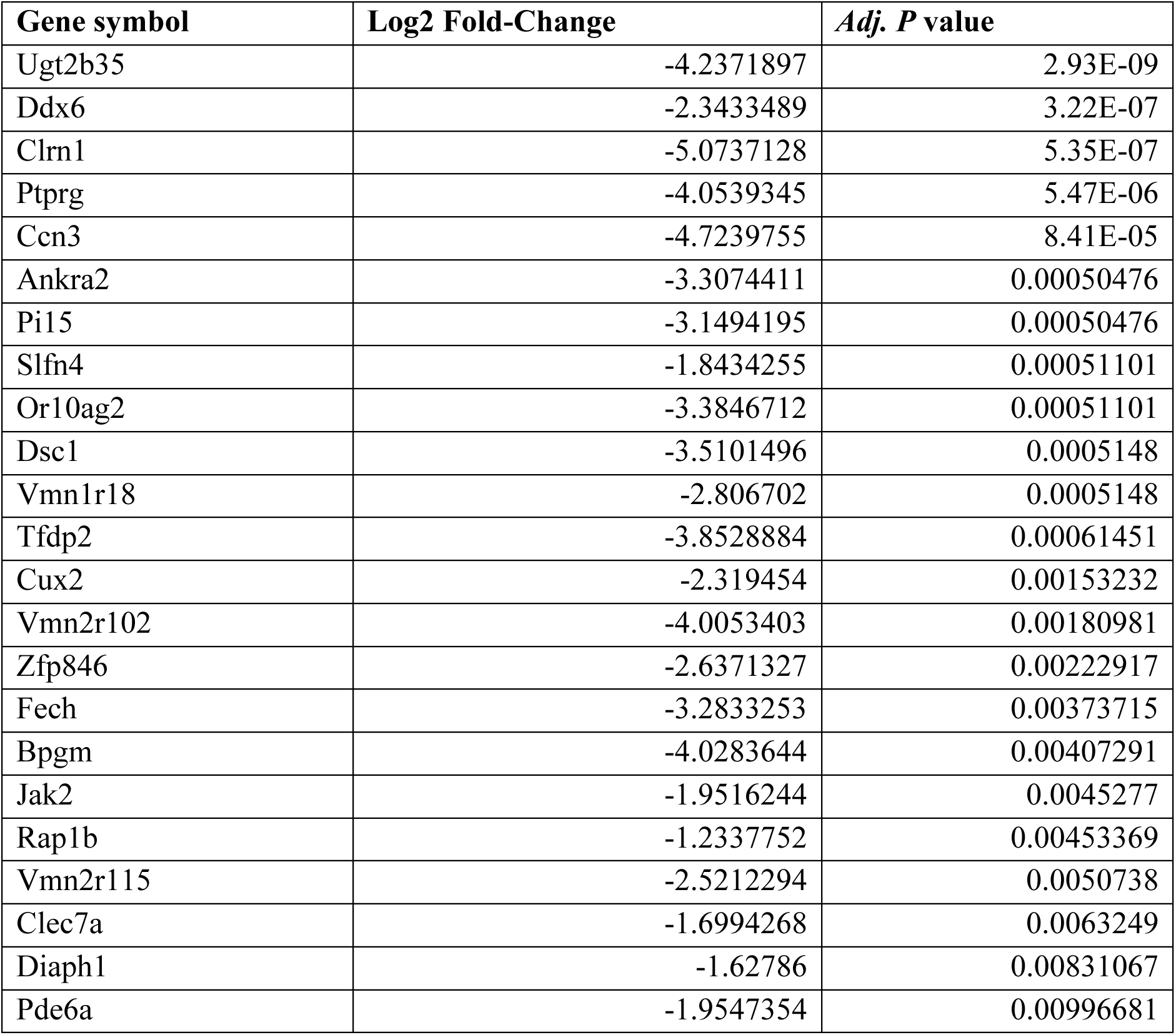
Transcripts with low abundance in blood neutrophils in MK-sufficient versus MK-depleted mice. Displayed by P values, adjusted P-values <0.10 (Wald test, FDR-adjusted) and fold-change < -1.5.

| Gene symbol | Log2 Fold-Change | Adj. P value |
| --- | --- | --- |
| Ugt2b35 | -4.2371897 | 2.93E-09 |
| Ddx6 | -2.3433489 | 3.22E-07 |
| Clrn1 | -5.0737128 | 5.35E-07 |
| Ptprg | -4.0539345 | 5.47E-06 |
| Ccn3 | -4.7239755 | 8.41E-05 |
| Ankra2 | -3.3074411 | 0.00050476 |
| Pi15 | -3.1494195 | 0.00050476 |
| Slfn4 | -1.8434255 | 0.00051101 |
| Or10ag2 | -3.3846712 | 0.00051101 |
| Dsc1 | -3.5101496 | 0.0005148 |
| Vmn1r18 | -2.806702 | 0.0005148 |
| Tfdp2 | -3.8528884 | 0.00061451 |
| Cux2 | -2.319454 | 0.00153232 |
| Vmn2r102 | -4.0053403 | 0.00180981 |
| Zfp846 | -2.6371327 | 0.00222917 |
| Fech | -3.2833253 | 0.00373715 |
| Bpgm | -4.0283644 | 0.00407291 |
| Jak2 | -1.9516244 | 0.0045277 |
| Rap1b | -1.2337752 | 0.00453369 |
| Vmn2r115 | -2.5212294 | 0.0050738 |
| Clec7a | -1.6994268 | 0.0063249 |
| Diaph1 | -1.62786 | 0.00831067 |
| Pde6a | -1.9547354 | 0.00996681 |

**Supplementary Table 13.** Top 40 GO Cellular Biological Process annotation terms enriched in blood neutrophils in MK-sufficient mice. Displayed by adjusted P values (FDR).

| **GO annotation term ID** | **GO annotation term, Biological Process Ontology** | **Fold-enrichment** | **Adj. P value** |
| --- | --- | --- | --- |
| GO:0048661 | positive regulation of smooth muscle cell proliferation | 23.9829303 | 0.00046681 |
| GO:0050727 | regulation of inflammatory response | 23.6631579 | 0.00046681 |
| GO:0032496 | response to lipopolysaccharide | 13.6518219 | 0.00046681 |
| GO:0071222 | cellular response to lipopolysaccharide | 13.2278522 | 0.00046681 |
| GO:0042127 | regulation of cell population proliferation | 13.1461988 | 0.00046681 |
| GO:0002281 | macrophage activation involved in immune response | 81.9109312 | 0.00046681 |
| GO:0033209 | tumor necrosis factor-mediated signaling pathway | 26.2923977 | 0.00105856 |
| GO:0006954 | inflammatory response | 7.64502024 | 0.00174944 |
| GO:0060339 | negative regulation of type I interferon-mediated signaling pathway | 46.2974828 | 0.00174944 |
| GO:0006952 | defense response | 21.1908877 | 0.00174944 |
| GO:1900016 | negative regulation of cytokine production involved in inflammatory response | 31.3188854 | 0.00511424 |
| GO:0032755 | positive regulation of interleukin-6 production | 15.2665535 | 0.00526988 |
| GO:0031663 | lipopolysaccharide-mediated signaling pathway | 28.0221607 | 0.00567242 |
| GO:0043124 | negative regulation of canonical NF-kappaB signal transduction | 14.1978947 | 0.00567242 |
| GO:0030316 | osteoclast differentiation | 27.3036437 | 0.00567242 |
| GO:1904707 | positive regulation of vascular associated smooth muscle cell proliferation | 26.6210526 | 0.00573792 |
| GO:0060337 | type I interferon-mediated signaling pathway | 23.1487414 | 0.0082012 |
| GO:0007259 | cell surface receptor signaling pathway via JAK-STAT | 20.4777328 | 0.01114586 |
| GO:1901224 | positive regulation of non-canonical NF-kappaB signal transduction | 19.7192982 | 0.0113499 |
| GO:0043922 | host-mediated suppression of viral transcription | 59.1578947 | 0.0113499 |
| GO:1904996 | positive regulation of leukocyte adhesion to vascular endothelial cell | 59.1578947 | 0.0113499 |
| GO:0032731 | positive regulation of interleukin-1 beta production | 19.0150376 | 0.0113499 |
| GO:0009617 | response to bacterium | 10.4396285 | 0.01153623 |
| GO:0097190 | apoptotic signaling pathway | 18.3593466 | 0.01153623 |
| GO:0045638 | negative regulation of myeloid cell differentiation | 44.3684211 | 0.0168406 |
| GO:0061635 | regulation of protein complex stability | 44.3684211 | 0.0168406 |
| GO:0035994 | response to muscle stretch | 35.4947368 | 0.0254949 |
| GO:0043525 | positive regulation of neuron apoptotic process | 12.6766917 | 0.02803273 |
| GO:0071356 | cellular response to tumor necrosis factor | 12.6766917 | 0.02803273 |
| GO:0050729 | positive regulation of inflammatory response | 12.1004785 | 0.02889409 |
| GO:0043065 | positive regulation of apoptotic process | 5.52877521 | 0.02889409 |
| GO:0001937 | negative regulation of endothelial cell proliferation | 29.5789474 | 0.02889409 |
| GO:0032703 | negative regulation of interleukin-2 production | 29.5789474 | 0.02889409 |
| GO:0043330 | response to exogenous dsRNA | 29.5789474 | 0.02889409 |
| GO:0051607 | defense response to virus | 7.31850244 | 0.02889409 |
| GO:0032680 | regulation of tumor necrosis factor production | 28.3957895 | 0.02987149 |
| GO:0042981 | regulation of apoptotic process | 7.09894737 | 0.02987149 |
| GO:0038066 | p38MAPK cascade | 27.3036437 | 0.02987149 |
| GO:0048545 | response to steroid hormone | 27.3036437 | 0.02987149 |
| GO:0060907 | positive regulation of macrophage cytokine production | 26.2923977 | 0.03139892 |

**Supplementary Table 14.** Top GO Cellular Biological Process annotation terms enriched in blood neutrophils in MK-depleted mice. . Displayed by adjusted P values (FDR).

| GO annotation term ID | GO annotation term, Biological Process Ontology | Fold-enrichment | Adj. P value |
| --- | --- | --- | --- |
| GO:0045494 | photoreceptor cell maintenance | 28.3957895 | 0.00459682 |
| GO:0045597 | positive regulation of cell differentiation | 20.3592453 | 0.00807609 |
| GO:0007605 | sensory perception of sound | 11.9362832 | 0.00807609 |

**Supplementary Table 15.** Transcripts with high abundance in marrow neutrophils in MK-sufficient versus MK-depleted mice. Displayed by P values, adjusted P-values <0.10 (Wald test, FDR-adjusted) and fold-change > 1.5.

| Gene symbol | Log2 Fold-Change | *Adj. P value* |
| --- | --- | --- |
| Atp6-ps | -22.347784 | 2.11E-07 |
| Cnbd2 | -1.2486085 | 2.11E-06 |
| Il12b | -3.0997804 | 0.00021298 |
| Tnfrsf26 | -0.9496207 | 0.00195586 |
| Epx | -1.839965 | 0.00264247 |
| Chd9 | -0.6955772 | 0.00490912 |
| Serinc3 | -1.0895468 | 0.02545702 |
| Cdk5rap1 | -1.7557634 | 0.03343685 |
| Zfp933 | -1.3322734 | 0.04445215 |
| H4c18 | -1.5134998 | 0.0711394 |
| Tubb1 | -3.4430552 | 0.09782179 |
| Prg2 | -1.4989064 | 0.09782179 |
| Cep85 | -1.2213884 | 0.09782179 |
| Szrd1 | -0.7479852 | 0.09782179 |

**Supplementary Table 16.**
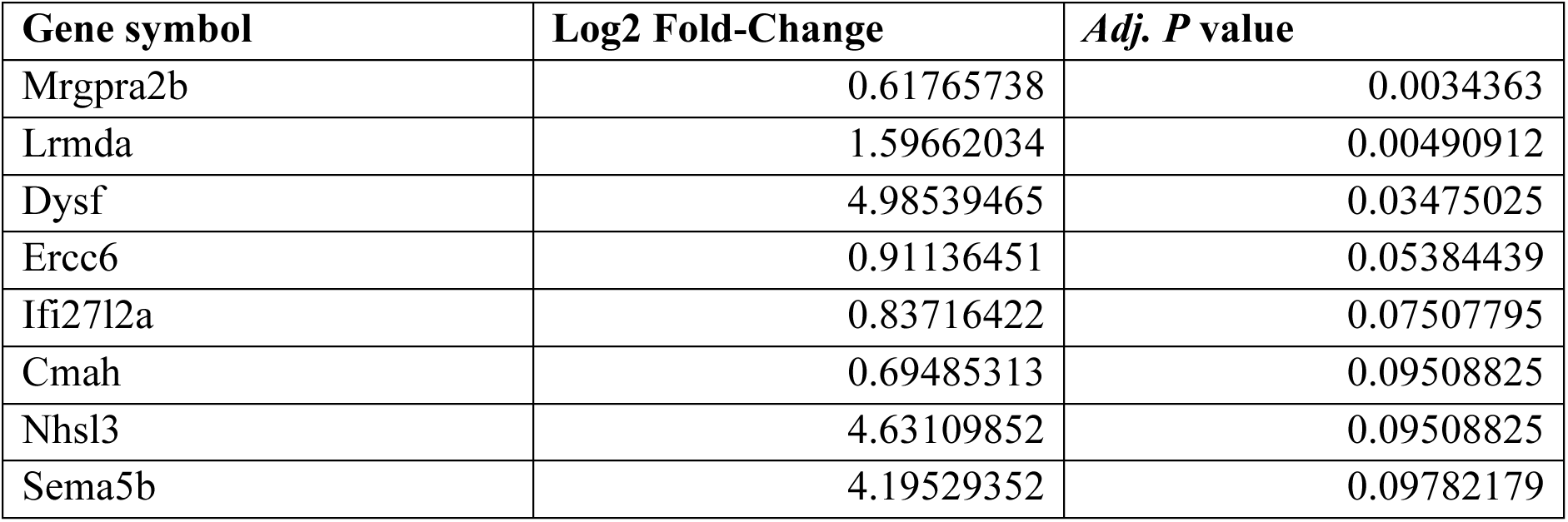
Transcripts with low abundance in marrow neutrophils in MK-sufficient versus MK-depleted mice. . Displayed by P values, adjusted P-values <0.10 (Wald test, FDR-adjusted) and fold-change < -1.5.

**Supplementary Table 17.**
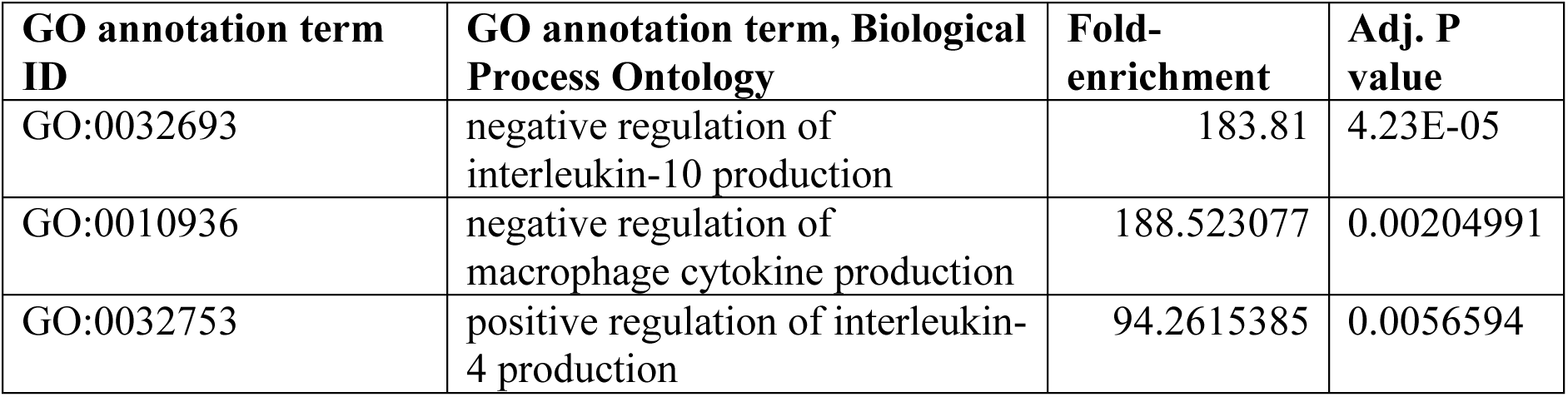
Top GO Cellular Biological Process annotation terms enriched in marrow neutrophils in MK-sufficient mice. . Displayed by adjusted P values (FDR).

